# The Test Used to Choose Between Pesticide Survival Models Tracks Lesion Persistence, Not Hazard Structure

**DOI:** 10.64898/2026.11.03.764107

**Authors:** Jong-Hyeon Lee, Chan Young Joe

## Abstract

Aquatic pesticide risk assessment predicts survival with toxicokinetic–toxicodynamic models that offer two hazard structures, cumulative or peak-based, which disagree about which exposure patterns are dangerous; for imidacloprid, published choices differ across eight species. The choice is made from one observation: deaths after transfer to clean water are taken to refute the peak-based structure. That inference holds only if damage is falling, which is not measured in these tests but taken from a fitted model. The observation responds to the compound and design, not the hazard structure. Across thirty-two *Gammarus pulex* treatments the post-exposure death rate places anticholinesterases above uncouplers, in the order of lesion persistence. For diazinon, one series returns opposite verdicts over its sixteen-day window and its first four days. For imidacloprid in *Daphnia magna*, one damage stage favours the cumulative structure by 78 likelihood units and two stages the peak-based one by 7. Four conditions make a design informative; thirteen of thirty-six treatments in a thirteen-compound archive meet them. Spreading a fixed exposure over longer intervals raises mortality in one of five species, which no repair law reproduces with the models’ kinetics. For neonicotinoids, predictions for such patterns are unvalidated, not conservative.

## 1. Introduction

Aquatic risk assessment for pesticides predicts survival from a measured exposure profile with toxicokinetic–toxicodynamic models; in the European Union (EU) the prescribed model is the General Unified Threshold model of Survival (GUTS).^1–3^ GUTS offers two reduced variants, called here the cumulative and the peak-based structure: in the first, hazard accumulates with the excess of damage over a threshold, integrated over time; in the second, an individual dies when damage first exceeds its tolerance. The two are not settings of one model but two dose metrics computed from one damage trajectory — one integrates it, the other takes its running maximum — so a long, low exposure produces more hazard under the first and a brief spike under the second, and neither is always the more conservative. Because both read the same trajectory, an error in it shifts both predictions the same way, so the true survival need not lie between them.^4^ Guidance recommends fitting both and taking the more protective result,^3^ which defers the choice rather than removing it. The choice therefore decides which field exposure profiles an assessment passes as safe. A companion manuscript submitted alongside this one classifies the two structures and asks which predicts better;^4^ this paper asks what is measured by the observation used in practice to choose between them: deaths after exposure ends.

Nothing in the standard data settles the choice. For imidacloprid the published verdicts split three ways across eight species (§4.1). ^5–8^ These verdicts rest on one observation: animals that went on dying after transfer to clean water were taken to refute a peak-based hazard. Nor does the regulatory battery contain a test that separates the structures: they diverge only after damage falls, and a four-day acute test ends before that (§S9.5, Figure S6a), so in practice the choice rests on that single observation in experiments run for other purposes.

That observation has a sound logic and an unobserved premise. The structures diverge because a running maximum stops once damage falls while an integral keeps growing (§2); deaths in clean water therefore refute the peak-based structure *if damage is falling there* . Damage is never measured. Whether it was falling is inferred from a model fitted to the same survival counts, over whatever window the design provides. The eight-species disagreement therefore admits two readings: the correct structure is a property of the species, in which case the verdicts should follow taxonomy, and they do not (§4.1); or the published comparisons fixed the damage model at a single stage, and each verdict reflects how well that single stage fits the species.

The premise that damage is falling is least secure for neonicotinoids, the class for which its failure would matter most. They reach surface water as repeated pulses separated by days — the exposure pattern on which the structures disagree — and they are the class these models are most often calibrated on. ^5,8–11^ Their receptor binding shows no measurable dissociation over eight days, ^12^ so the chemistry gives no ground for assuming that the lesion, and with it damage, falls once exposure ends. The question survives the 2018 EU restriction on outdoor use of three neonicotinoids: the butenolides and sulfoximines replacing them are agonists at the same receptor.^13,14^

The conditions under which each structure can produce post-exposure deaths have been derived,^15^ but they are conditions on the unobserved damage trajectory; what the observation itself responds to has been asked neither in that derivation nor in the assignments that rely on it.^5^ The question divides into three.

1. What does the clean-water observation respond to: the hazard structure it is taken to discriminate, or how long the lesion persists and how long the animals were watched?
2. If the latter, which of the published verdicts survive, and can any published design decide between the two structures?
3. Is there a pattern in the published pulsed-exposure data that neither structure can produce, whatever damage model and repair kinetics are assumed?

The analysis is confined to what published data can settle (§S2). Most is arithmetic on published counts: the whole archive of pulsed-exposure survival experiments (thirteen compounds, four modes of action) is examined to find what the observation responds to, and each treatment is scored against the conditions a design must meet to separate the structures. Two results need more than arithmetic: one refit of a *Daphnia magna* series and three propositions about the running maximum, two applied to a five-species experiment varying the pulse interval. The consequences for the model choice are worked out for neonicotinoids, for which the published designs cannot settle it.

The short answer to the first question is how long the lesion persists and how long the animals were watched; a verdict read from that observation therefore records the compound and the design, not the organism. For neonicotinoids the choice cannot be made. Two consequences for practice do not wait on that choice: a minimum clean-water window set by compound class, and a rule for how deep the damage model must be, both computable before a test is run (§4.7, §5).

## 2. The Clean-Water Observation and What It Assumes

### Why the observation is a test

Under a peak-based hazard the survival probability depends on the damage trajectory only through its running maximum, and damage is driven by an input *u* minus repair,

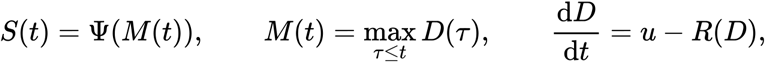

for a decreasing *Ψ* from the tolerance distribution and any non-negative *R*.^4^ The consequence is immediate: *M* is constant whenever *D* is falling, so survival must be flat apart from background mortality during any interval in which damage declines. That is what makes the clean-water observation a test: it can refute the peak-based structure, but only if damage is falling while the counts are taken.

### Freezing holds for any repair law

Once *u* reaches zero, damage cannot rise, the running maximum is constant, and any hazard depending on the trajectory only through that maximum is frozen. Neither the form nor any parameter of the repair law enters (Proposition 1, §S1). Jager’s derivation of when each structure can produce post-exposure deaths is not in dispute;^15^ what this paper adds is how the condition that derivation depends on (that damage is falling) is established in practice — by a fitted model.

### Three facts about the running maximum organise what follows

The first is the freezing: once the input stops, *M* cannot rise (Proposition 1). The other two bound how *M* responds to the shape of an exposure: separating two equal pulses cannot raise it, for any repair law meeting the conditions of §S1 (Proposition 2; inputs as in §4.6), and, for convex repair, cannot lower it by more than half (Proposition 3); concave repair can go below the half (§S10.2). Together they bound what a peak-based structure can express before anything is fitted: the first makes the clean-water observation a test, the second shows a pattern in these data that neither structure can produce with the kinetics the framework uses, and the third gives the peak-based structure a bound an observation could violate (§4.6). Proposition 1 holds for any non-negative repair law, Proposition 2 under the conditions of §S1, and Proposition 3 needs it convex; those conditions, with that input shape, are everything the paper assumes about structure.

### The operative quantity is *u*, and *u* is not the parent compound

In a single-stage model the input to damage is the internal concentration, so clearance fixes when *u* reaches zero. In a multi-stage model it is the preceding state, which need not vanish with the parent: receptor occupancy or subcellular stress is invisible to a total-residue measurement. How much such a state recovers between pulses can still be bounded from the counts alone (§S8). The refit of §4.3 changes neither *Ψ* nor *R*; it gives *u* one more stage, so the input persists after the parent has cleared. Under a peak-based hazard survival keeps falling only while *M* is still rising, and *M* rises only while *u* exceeds the repair term. The post-exposure decline rate of §4.5 therefore measures how long the input persists, not how damage becomes death. That persistence is set by the compound’s chemistry: how long the adduct or the occupancy lasts.

### What the assignments in the literature did

Focks and colleagues assigned structures from the clean-water behaviour: flat survival to individual tolerance (the peak-based structure), declining survival to stochastic death (the cumulative one).^5^ Whether that assignment records the death mechanism or the damage model is what §4.3 and §4.5 test.

### What the companion does with the same counts

The companion shares this freezing result and one archive but asks a different question. It measures the carry-over ratio between paired pulses to see which structure predicts survival better;^4^ this paper measures the rate of decline over the clean water that follows the last pulse. The two quantities are read from the same counts but bear on different things: one on the structures, the other on the test.

## 3. Materials and Methods

### Data

All data are published. Survival and exposure series for imidacloprid in *D. magna* are from Li *et al*.;^6^ three treatments share a first 24-h pulse and differ only in the timing of the second (2, 7 or 15 d), giving seventeen survival observations over 22 d. Survivor counts for five macroinvertebrate species under three splittings of one total exposure (§4.6), and for three neonicotinoids under two intervals in *Cloeon dipterum*, are from Focks *et al*.^5^ and their raw data file.^16^ Daily survival counts with concurrent controls, and measured exposure profiles, for thirteen compounds in *G. pulex* are from the data supplement of Ashauer *et al*.^17^ A fourteenth sheet in that supplement reproduces the chlorpyrifos experiment of ref 21, counted as a separate experiment (§S3). Sources and provenance are in §S2.

### Models

Toxicokinetics follow the published one-compartment fits.^6^ Damage is either single-stage, d*D*/d*t* = *k*_*a*_*C*_*b*_ − *R*(*D*), or two-stage, with a state upstream of damage, d*D*_1_/d*t* = *k*_*a*_*C*_*b*_ − *k*_off_*D*_1_ and d*D*_2_/d*t* = *k*_12_*D*_1_ − *R*(*D*_2_) with *k*_12_ fixed at 1 h^−1^ because only its product with *k*_*a*_ is identified. Repair is either proportional, *R* = *k*_*r*_*D*, or Michaelis–Menten, *R* = *R*_max_*D*/(*K*+*D*). Hazard is cumulative, *H*_*A*_ = *b*_*w*_∫[*D*−*z*]^+^d*t*, or peak-based, *H*_*B*_ = *k*[*M*−*θ*]^+^. Background survival enters multiplicatively as the measured control series (§S9).^6^

### Fitting and model comparison

Survival counts are fitted by conditional binomial likelihood, seven variants (§S9, Table S29) compared by AICc with Akaike weights. Each variant is fitted with one parameter set to all three treatments, so the dependence on the pulse interval is predicted rather than fitted. Estimation details are in §S9; the likelihood surface has multiple local optima, and randomised starts stop about eight units short of the optimum, so each fit starts from the optimum of the simpler variant nested in it. The DNOC series is fitted with four variants — single- and two-stage damage under each hazard structure, the two-stage form with saturating repair — with toxicokinetics from the measured half-life and survival relative to day 1 (§S12).

### Analyses from counts alone

For each treatment the clean-water window opens two days after measured exposure last exceeds 5% of its maximum (the allowance a multi-stage cascade needs to reach its peak — 1.6 to 1.9 d under the companion’s three-compartment diazinon model; §S3, §S6) and closes when observation ends. For diazinon, where a trajectory has been fitted, the window opens at the fitted damage maximum instead; the two rules agree to within a day (§S6). Survival at each end is control-corrected as (*n*_trt_/*N*_trt_)/(*n*_ctl_/ *N*_ctl_) and the change tested by Fisher’s exact test, with Benjamini–Hochberg control of the false discovery rate. Designs differ in length, so the quantity compared across them is the decline rate

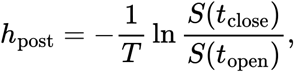

where *T* is the length of the window. Treatments are nested in compounds, so the modes of action are compared at compound level by the Mann–Whitney test, each compound summarised by the median of its treatments (§S4.2), and the class difference is tested for confounding by the mortality already accumulated when the window opens (§S4.3). Intervals on the rate are profile-likelihood limits from the two binomials, treatment and control, over the window. The rate serves two purposes: Fisher’s test on the window asks whether the fall it summarises is significant — the literature’s flat-or-declining verdict — and the design criterion of §4.7 is its product with the window length. Nothing in §4.5 or §4.7 is fitted, and the excess-damage ratio of §S8 is also a ratio of counts, with a zero count replaced by one-half.

### Robustness

Three checks cover the comparison of §4.3 (profile likelihood, leave-one-out prediction, a bootstrap under three descriptions of the reading error; §S14–§S15), and the census is repeated under nine variants of the window rule (§S15.3). The propositions of §2 are stated and explained in §S1, which proves the first and third and cites a technical report for the general proof of the second; §S10 checks them numerically.

## 4. Results

### 4.1 The problem: one compound, eight species, no agreement

A hazard structure should be a property of the organism, not of the compound. The published verdicts for imidacloprid do not show that pattern: five species are assigned to the cumulative structure,^5,6,8^ two to the peak-based structure,^5,7^ and one to neither, with no taxonomic pattern. One laboratory reached opposite conclusions for two mayflies in the same experiment.^5^ Figure 1 shows the series behind six of the eight; the other two report no counts. Either a hazard structure is not a property of the organism after all, or these verdicts are not measuring it. Two papers in a 2024 exchange, arguing opposite sides, agree that the verdicts are not measuring it: slower repair can substitute for individual tolerance within one fit,^18^ and a repair rate inferred from survival is an assumption rather than a measurement.^19^

**Figure 1.**
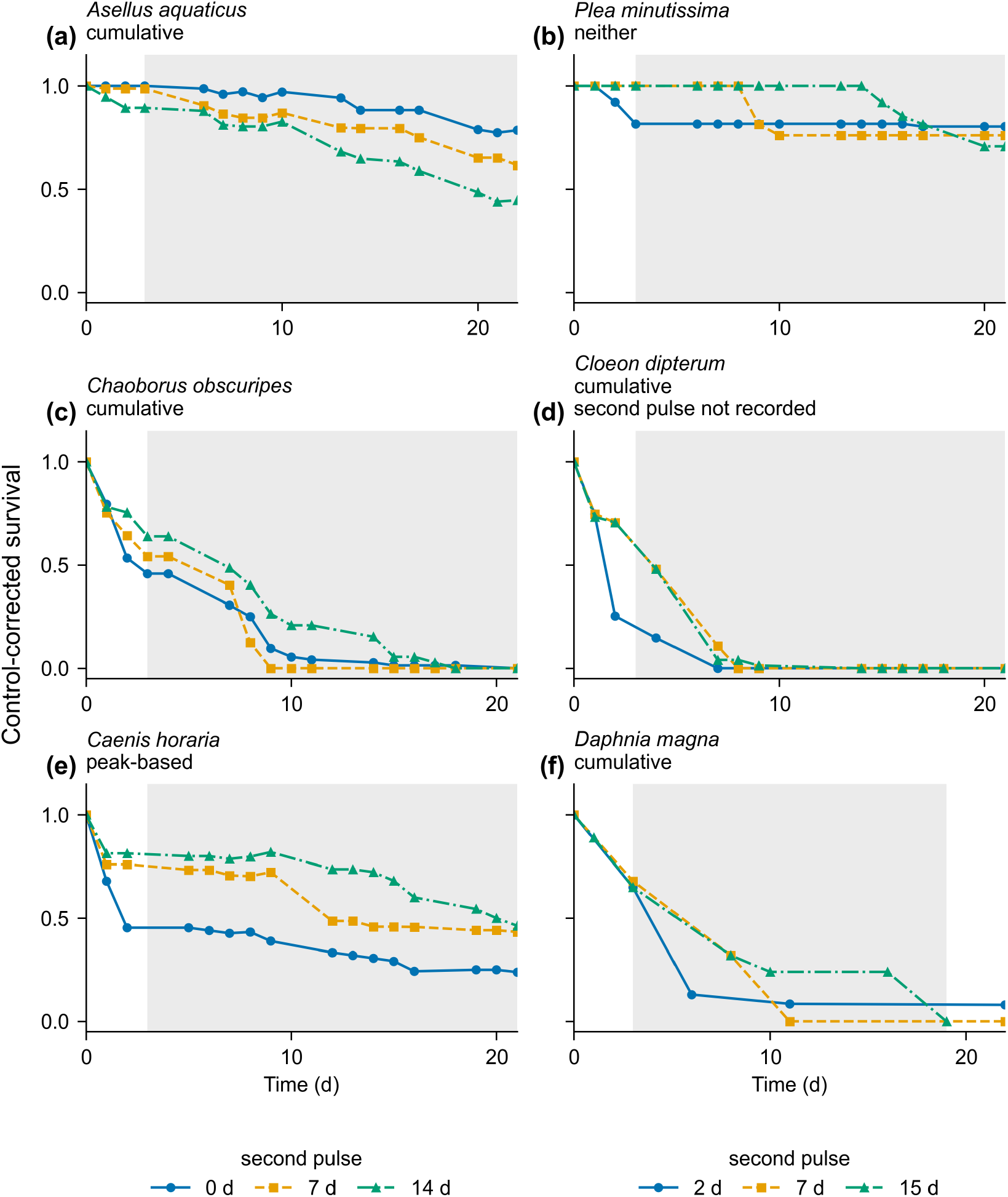
Imidacloprid across every species with published survival counts. Control-corrected survival for the five macroinvertebrates of the interval design^5,16^ and for *Daphnia magna*,^6^ each panel labelled with the structure the literature assigns. The five interval-design panels split one total exposure as in §4.6, a second pulse at 0 d being the single 48-hour one, and shading marks where a peak-based hazard predicts flat survival (§2). The *Daphnia magna* panel is the three treatments of §3 and carries its own legend; its shading closes at day 19, where the 15-day treatment was last observed. *Gammarus pulex* and *Deleatidium* spp. have verdicts but no counts.

### 4.2 On one damage stage, the peak-based structure fails — by counts and by fit

In the *D. magna* treatment with a 15-d interval, survival fell from 89% on day 1 to 24% on day 10 and then stayed constant until the second pulse. On the published single-stage trajectory damage peaks on day 1.14, so until the second pulse the running maximum, and with it a peak-based hazard, is frozen (§S11.2; Figure 2b): survival can fall only as fast as the controls. Placing all 22-day background mortality (control survival 76%) before day 10, most generous to the peak-based structure, predicts 89% × 0.76 = 68% on day 10 against 24% observed, a factor of 2.8. No fit can close the gap: the parameters cancel from the ratio while the maximum is frozen (§S11.2).

**Figure 2.**
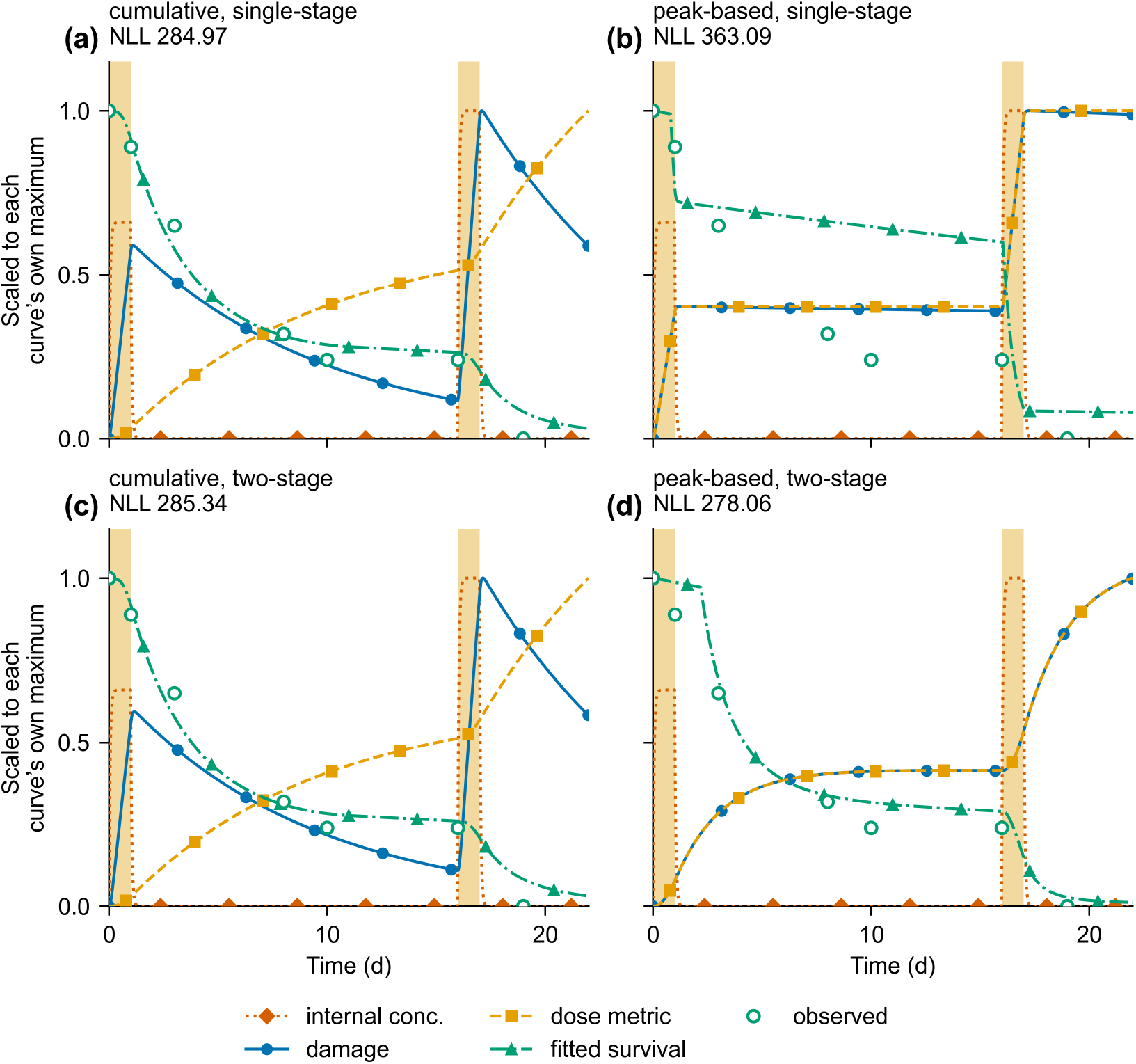
One treatment, two damage models, both hazard structures. *Daphnia magna*, imidacloprid, the 15-day treatment, fitted as in §3. Internal concentration, damage, the dose metric each structure reads from it, the fitted survival and the counts, each scaled to its own maximum; gold bands are the pulses. (a, b) On a single-stage trajectory damage peaks early (day 1.1 under the cumulative fit, day 1.3 under the peak-based one) so the cumulative metric keeps rising while the running maximum is fixed for the next fourteen days. The peak-based fit in (b) then falls only as fast as the controls, which is the comparison of §4.2. (c, d) With a state that outlasts the parent the first-interval peak moves to day 13.8 in (d), and the maximum is still rising when the test ends. Parameters are in Table S30.

A fit gives the same answer (Figure 2a,b): fitted across all three treatments with a single parameter set, the peak-based structure on the published single-stage trajectory gives NLL 363.1 against 285.0 for the cumulative structure. Two toxicokinetic changes that could favour the peak-based structure by delaying the damage peak fail. No metabolite can be invoked: three were measured and none detected.^6^ The exoskeleton holds half the imidacloprid mass during depuration,^6^ but at its measured release rate delays the damage peak by one hour. Neither closes a gap of 78 likelihood units.

### 4.3 A persisting upstream state reverses the verdict

One extra state in the damage model turns a 78-likelihood-unit deficit for the peak-based structure into a 7-unit advantage (Figure 2b,d). §4.2 rejected one hazard structure on one damage model, and the reduced variants of §1 omit two extensions that the companion’s classification allows: an upstream state through which damage forms, and saturating repair.^4^

Fitting the extensions separately shows which one changes the verdict (Table S31). Against the cumulative structure refitted at the same damage model, the peak-based structure trails by 78.1 likelihood units on a single-stage trajectory and leads by 7.3 with an upstream stage — an improvement of 85 units for one added parameter, which, once penalised, leaves it 12.0 AIC units ahead of the single-stage cumulative model and at least twelve ahead of every cumulative variant (Table S29). The extension does not help every variant: with the added stage the cumulative fit reduces to its single-stage form, gaining nothing from a later peak, and adding saturating repair on top worsens the information criterion (§S11). The saturation this class does show is in formation, not repair: receptor binding is capacity-limited, so it bounds how much damage forms, not how fast it is removed.^12^

What the upstream stage does is move the damage peak. On a single-stage trajectory damage peaks within a day of each pulse, so the running maximum is updated only in two brief episodes and the peak-based hazard must account for 22 days of deaths with them. A state that outlasts the parent keeps the input positive for days, and the fitted damage rises almost to the end of the test: the maximum is frozen over only a small part of it (Figure 2d, §S11), and while the maximum is still rising the two structures respond to the trajectory in the same way (§S12). How far the peak moves is set by the upstream rate alone, and the fitted value falls between the rate at which the compound leaves the animal and the rate at which it dissociates from its receptor: the range in which a stage must be represented explicitly (§S9.5).

The data identify the persistence, not the repair law: the profile likelihood localises the upstream half-life to 1.53 d (95% interval 1.13–2.41 d) and puts terminal repair beyond 10^2^ d, while the half-saturation constant is not identified at all (§S15.2). The companion paper reaches the same conclusion for organophosphates.^4^ Which repair law the model carries makes no difference to any comparison here. This is the pattern §4.5 measures across compounds: what survival identifies is how long damage persists.

Two features of the design make the comparison informative. First, one parameter set must reproduce three interval-dependent curves (§3). Fitted to one treatment alone the verdict reverses — the simplest cumulative model then has the lowest information criterion and the jointly winning peak-based model trails every cumulative variant (Table S29) — because a single curve constrains only its own shape, whereas three intervals require the model to predict how survival depends on the interval, which the single-stage trajectory cannot. The result also holds out of sample: holding each treatment out and predicting it from the other two, the two peak-based variants with the upstream stage are the best two of the six and the same hazard structure on a single-stage trajectory is the worst — a swing of 74 predictive likelihood units from the damage model alone (Table S36, §S14); the cumulative structure again gains nothing from the added stage (§S11). Second, the comparison is informative only where damage recovers within the experiment; where it does not, the two structures coincide (§S12).

### 4.4 One compound, one experiment: the verdict follows the window length

The clean-water verdict flips for two reasons, and neither is the hazard structure. One has just been shown: §4.2 and §4.3 reach opposite conclusions from the same counts, differing only in the damage model. The other is the length of the window a design provides, and it can be shown within one compound.

That compound is diazinon in *G. pulex*, in its one published pulsed experiment: two one-day pulses 2, 7 or 15 days apart, seventy animals per treatment, with concurrent controls.^17,20^

One test applied to its three treatments (Figure S3, Table S14) shows the window deciding the result. Survival declines over both post-exposure windows, sixteen days in treatment A and eleven in treatment B. Shortening the window within one treatment removes the decline: over the full sixteen-day window, treatment A declines at 0.059 d^−1^ with an interval excluding zero; over its first four days, the rate is 0.049 and the interval includes zero (−0.005 to 0.123; §S4). A design leaving four days of clean water would have recorded no decline where sixteen days record one. Window placement depends on the damage model; the result does not (§S6).

That disposes of the first reading of §1: compound, mode of action, species and experiment are all held fixed, and only the window differs. The test measures the adequacy of the damage model over the window the design provides, and since that adequacy depends on the toxicokinetics, it varies between species for a fixed compound, the pattern of §4.1.

### 4.5 What the test tracks is how long the lesion persists

Counting significant declines will not show which compounds separate, because significance depends on window length, and windows are short: across the thirty-nine treatments recording both survival and exposure the median post-exposure window is four days (Table S3; §4.7 scores the thirty-six of the archive itself). The decline rate *h*_post_ of §3 — decline divided by window length — is not confounded in this way: it makes designs of different length comparable and admits windows too short for the flat-or-declining reading, and what then separates treatments is the compound. Pooling the thirteen-compound archive with the second series^21^ gives thirty-two treatments in eleven compounds; diazinon, whose windows are placed by a different rule (§S6), is reported separately and makes a twelfth compound (Figure 3a,b; Table S8).

**Figure 3.**
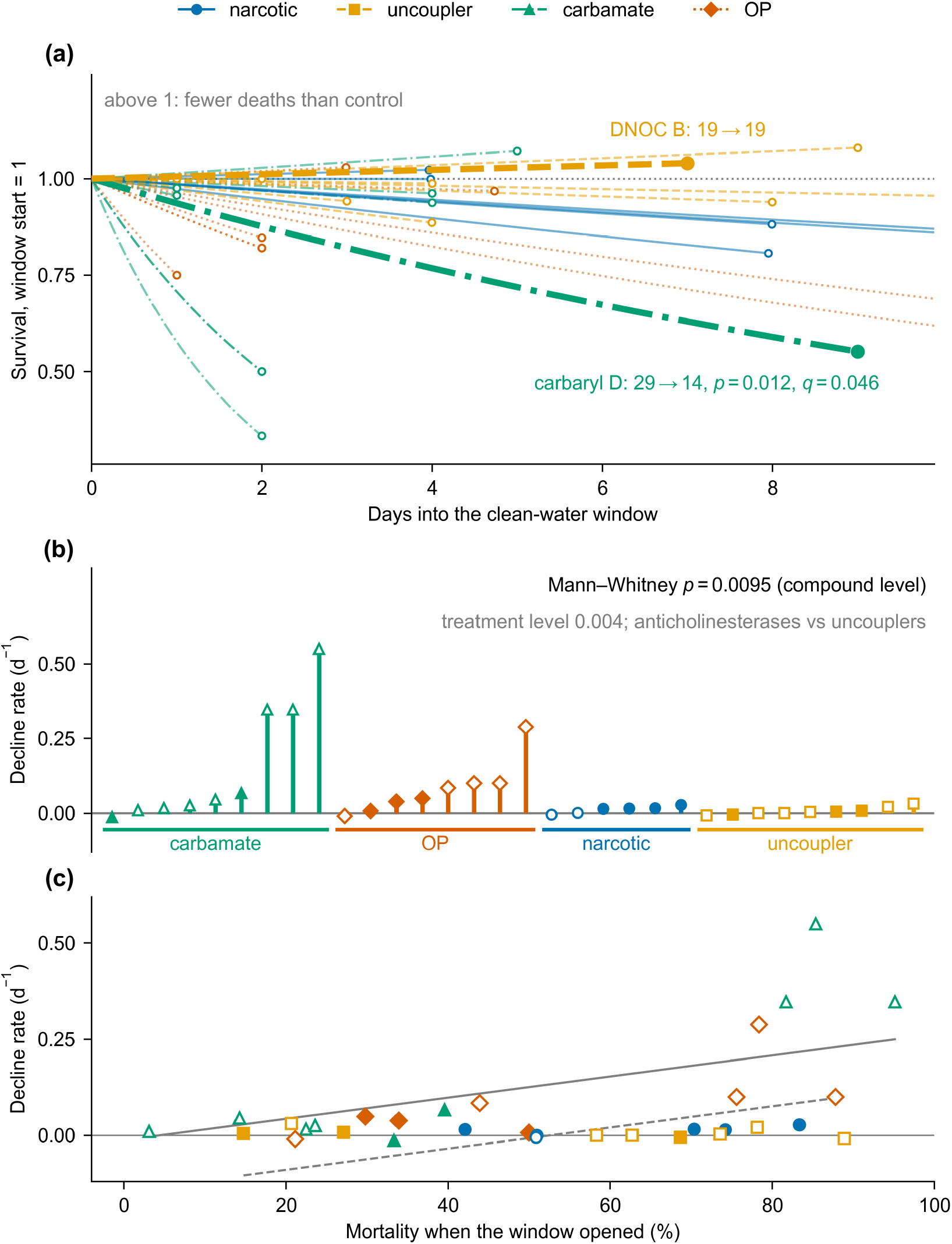
The post-exposure decline in every treatment that leaves a window, three ways. (a) Each treatment across its own clean-water window. Circles are the control-corrected survival at its close; the line between is the exponential the counts imply, not an observed path. Above 1 means fewer deaths than in the control. Bold: the matched pair of §4.5, where windows and group sizes are comparable. (b) The same treatments as decline rates *h*_post_, grouped by mode of action; filled markers meet §4.7. (c) The same rates against the mortality when each window opened; lines are the fit of Table S12. Depletion of the frail predicts a downward slope; the observed slope is upward.

The two mode-of-action classes separate (Table 1), the classes being assigned from pharmacology rather than from these data: nine uncoupler treatments have a median of 0.003 d^−1^, while fifteen of seventeen anticholinesterase treatments are positive (Tables S8, S9). The compound is the unit (§3): the ten compound medians of the two classes separate completely, with the smallest *p* that six compounds against four can return (0.0095, Mann–Whitney; §S4.2). The two narcotics, 1,2,3-trichlorobenzene and 2,4-dichloroaniline, lie between the classes at 0.015 and 0.014 d^−1^, and none of their six treatments differs from zero (Table S8). By lesion chemistry, neonicotinoids lie at the persistent end of the same ordering,^12^ and the same ordering appears in three further species (§S4.1, §S5).^22–24^

**Table 1.** What the five compound classes do once exposure ends. Brackets mark the entry the design cannot check. Internal half-lives are ln 2/*k*_out_ for whole-body elimination in *Gammarus pulex*^25–27^ (thiacloprid: the structure compartment of ref 12); *h*_post_ is the median across the treatments of §4.5, except the last row, three species of the interval series (§S7). The organophosphate median is over the five chlorpyrifos and three malathion treatments; with the two diazinon treatments of §4.4 it is 0.054, the compound-pooled value §4.7 uses (Table S5).

| **Class** | **Target** | **Recovery of the lesion** | ***t*<sub>1/2</sub> (d)** | ***h*<sub>post</sub> (d<sup>−1</sup>)** | ***n*** |
|---|---|---|---|---|---|
| Uncoupler | inner membrane, non-covalent | gradient re-forms as the compound clears | 0.39–3.33 | + 0.003 | 9 |
| Narcotic | membranes, non-specific | reverses as the compound clears | 0.18–0.29 | + 0.015 | 6 |
| Carbamate | AChE, carbamylation | spontaneous, within hours | 0.11–4.62 | + 0.045 | 9 |
| Organophosphate | AChE, phosphorylation | none; ageing makes it permanent | 0.20–1.54 | + 0.066 | 8 |
| **Neonicotinoid** | nAChR, agonist binding | **none measurable in 8 d**<sup>12</sup> | 0.13–2.60 | **( + 0.052)** | **(3)** |

The neonicotinoid entry is bracketed because its position cannot be checked. Its lesion is the longest-lived of the five, so its rate should be the highest; instead it falls among the anticholinesterases. Either the damage decays faster than the binding suggests, or the window, which opens the day after the pulse is removed (§S7), opens before damage has peaked and the rate understates the eventual decline.

The compounds the flat-or-declining reading calls silent are not counterexamples: aldicarb, carbofuran and chlorpyrifos all have positive rates on windows too short to resolve them (§S4, Table S8).

The decisive comparison is at matched window length and group size (Figure 3a, bold). A DNOC treatment loses none of the nineteen animals entering its seven-day window; a carbaryl treatment loses half of the twenty-nine entering its nine-day one (*q* = 0.046). The pattern repeats across the other uncoupler and carbamate treatments (Table S8; the carbaryl treatments in Figure S2).

Nor is the difference selection of the hardy: the uncoupler pulse was the harsher of the pair (Table S7), and across the thirty-two treatments the rate rises with prior mortality rather than falling as selection requires (Figure 3c, §S4.3).

Nor is time-reinforced toxicity — toxicity rising with exposure duration — the explanation of the ordering: in *C. dipterum* the ratio of four- to twenty-eight-day median effect concentration spreads fivefold between generations and only 1.3-fold across three neonicotinoids,^9^ so it follows the animal’s state, not the compound.

What separates the compounds is how long their damage persists, not their toxicokinetics. Internal half-lives overlap across these classes in the same species (0.11–4.62 d; Table 1), while the lifetime of the lesion runs from minutes for an uncoupler to no measurable dissociation in eight days for a neonicotinoid. The uncoupler, carbamate and organophosphate classes fall in the order their lesion chemistry predicts, a consistency rather than a test (Table 1). Fitting the same four variants to the DNOC series shows the same contrast: the upstream stage the neonicotinoid needs does not improve the uncoupler fit, and no cumulative variant comes close for either compound (Figure 4, §S12).

**Figure 4.**
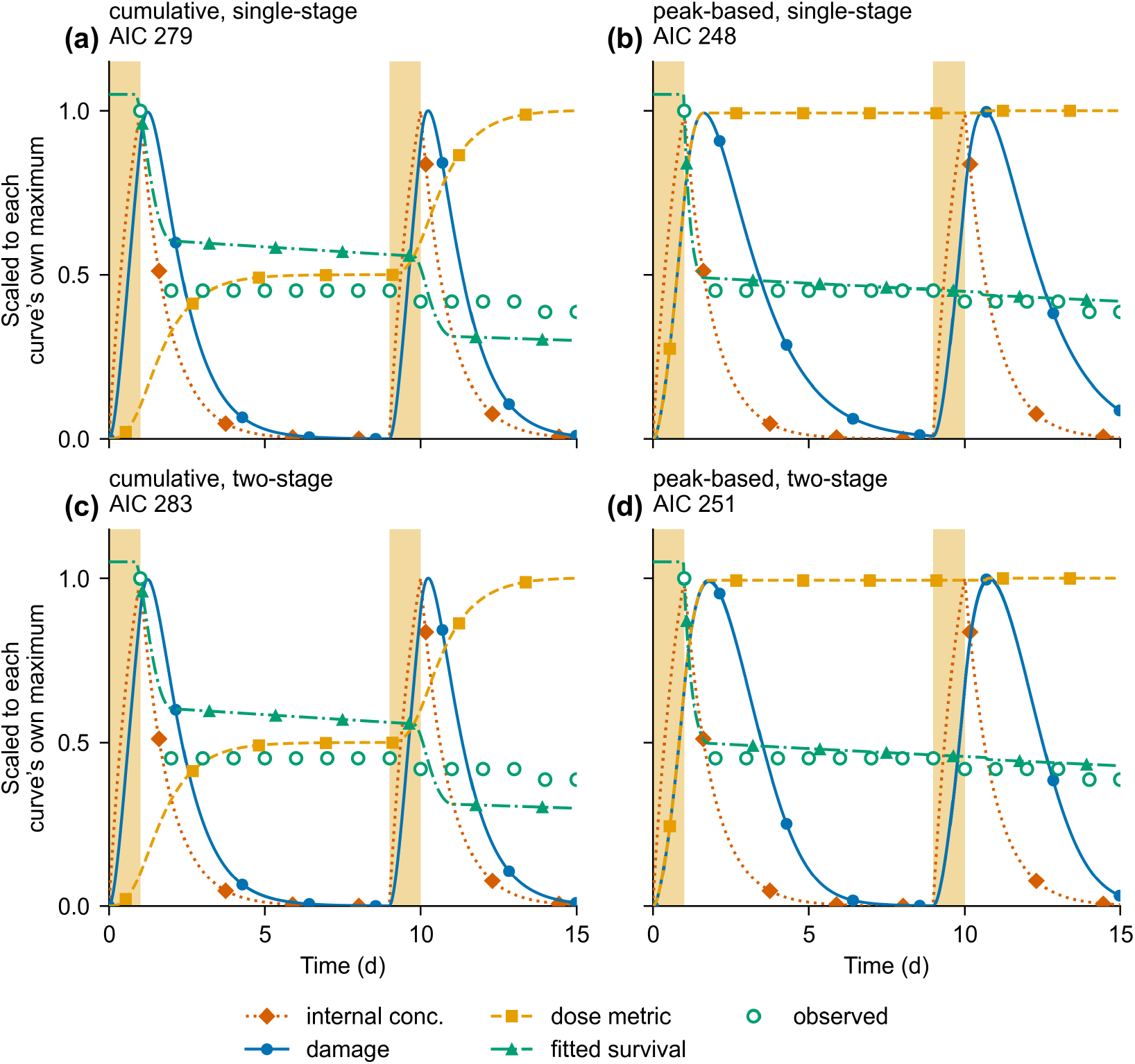
The same four-way comparison on an uncoupler. DNOC in *Gammarus pulex*, ref 17: two 24-hour pulses two, four or eight days apart, daily counts to day 15. The eight-day treatment is shown; the other two are fitted with the same parameters (§S12). Curves as in Figure 2, fitting as in §3. No mortality is observed across the eight clean-water days, and no variant predicts any. Over the whole series the cumulative structure is thirty-one AIC units worse, and the upstream stage the neonicotinoid needed costs 3.4 units here against a gain of 166 for imidacloprid.

### 4.6 A pattern no fitted variant reproduces, and the recovery a peak-based hazard can express

Focks *et al*. included a design isolating exposure pattern from exposure amount (the splitting shown in Figure 1) in *Asellus, Caenis, Chaoborus, Cloeon* and *Plea*, observed to day 21–28.^5,16^ Both structures predict that spreading a fixed exposure allows more recovery and so causes less mortality. One of five species shows the opposite (Figure 1, top left; Figure S4b): survivors fall from 51 to 40 to 29 in *A. aquaticus* as the interval lengthens (trend *p* = 3 × 10^−4^; Table S16). In *Plea minutissima* they fall from 61 to 57 to 53 by day 21 (*p* = 0.17); all its arms received a third pulse on day 21, so later counts do not compare equal exposures.

For log-concave damage inputs — every series of first-order stages, condition (V3) of §S1 — the peak damage from two identical pulses cannot rise as they are moved apart, for any repair law meeting the conditions of §S1 (Proposition 2; Figure S4a). Outside that class a numerical search reproduces the ordering only with two-compartment elimination and threshold-type repair (proportional repair cannot produce the rise for a single-peaked input), and the largest peak ratio on the grid, 1.241, is below the bound of 1.515 that linear integrations give for that configuration; matching the observed excess-damage ratio further requires a killing threshold at 0.895 of the single-pulse peak for *A. aquaticus* (3.3), four unmeasured properties in all (§S10.1). Nor does the cumulative metric, for which no proof exists: a numerical scan of both metrics over 162 parameter combinations reproduces the ordering in none (§S10, Figure S7).

Neither recorded endpoint shows what the first pulse left in the survivors: in *P. minutissima* it immobilises sixty-one animals in two days, after which immobilisation is zero and survival complete until the second pulse (§S7, Table S17). A developmental explanation can be set aside: the species that reverses, *A. aquaticus*, neither emerges nor reproduces within the test (§S7).^28^

An eroding tolerance does not reproduce the reversal either: at best it turns the ordering over by 1.3%, where the observed excess-damage ratio is 3.3 (§S13). What the data require is a non-monotone state (depleted during exposure and only partly restored in clean water) and the sublethal literature on this class reports that.^10,11^

#### How much recovery there was, and how much the structures can express

The counts alone bound how much recovery there was. In a second Focks design *C. dipterum* received three neonicotinoids with the second pulse at 7 or 14 d,^5^ and the excess-damage ratio of §S8 — a lower bound on the peak-damage ratio, near one meaning no recovery between 7 and 14 d — is 0.91 for imidacloprid and at most 0.27 for thiacloprid, while thiamethoxam kills every animal in both arms and leaves it undetermined (Table S19, Figure S5): imidacloprid recovers almost nothing in seven further days, and the cyano-substituted thiacloprid is the one that recovers (§S8).

The same argument bounds how much recovery a peak-based metric can express: separating two equal pulses reduces the peak by at most a factor of two, *M*(*Δ*′)/*M*(*Δ*) ≥ 1/2, however long the interval, for convex repair (Proposition 3, §S1). Concave repair breaks it only at rates far above those fitted here (§S10.2).

The observed 0.27 for thiacloprid lies below the bound of one half, and this is not a violation. Survival reports excess damage, whose ratio lies below the peak-damage ratio whenever that ratio is below one and the threshold below both peaks, so an excess ratio of 0.27 is what a peak ratio at or above one half looks like once a threshold is subtracted; Figure S5 shows the upstream half-life each observed ratio requires.

Each structure carries bounds that could refute it. The cumulative structure’s — a carry-over ratio of at most unity, which no choice of parameters can move — has been tested by the companion: fourteen of fifteen comparisons reach or exceed unity, and seven exclude it.^4^ Of the peak-based structure’s three, the recovery bound just derived has never been tested; the ceiling 1/*S*_1_ on the carry-over ratio, *S*_1_ being survival over the first pulse, is approached in five of six compounds;^4^ and the flat-survival bound the clean-water test reads is the least informative, because whether it appears violated depends on the window length (§4.4).

### 4.7 What a design must supply, and how many treatments supply it

The preceding sections specify what a design must supply for the clean-water observation to bear on the hazard structure. The damage peak must be locatable in time: without a measured exposure profile the window cannot be placed. The change across it must be correctable: without concurrent controls a decline cannot be separated from background mortality. The window must be long relative to the recovery timescale — three upstream half-lives, 4.6 d for imidacloprid (§4.3) — because a test resolves the product of the decline rate and the window length.^4^ The group must be large enough: ten animals is the smallest in which a halving of survival reaches significance (§S3.1).

That product can be computed before an experiment is run: at the median anticholinesterase rate, 0.048 d^−1^, a 4.6-day window gives 0.22, a 20% loss, short of the 0.35 (a 30% loss) that twenty animals can detect, whereas the longest-window chlorpyrifos treatment gives 0.03, and is silent. The requirement therefore follows from lesion persistence alone: the window each class needs is 0.35 divided by its median rate — 6.5 d for an organophosphate, 7.9 d for a carbamate, 24 d for a narcotic — and for an uncoupler, whose median rate is 0.003 d^−1^, it would be 111 d, longer than any test (Table S5, §S3.2).

Applied to the archive, thirteen of the thirty-six treatments meet all four: two each of 1,2,3-trichlorobenzene, 2,4-dichloroaniline, 2,4-dichlorophenol, carbaryl, malathion and diazinon, and one of DNOC (Figure S1, Table S4). Against the class windows of Table S5, five of the thirteen can also bear on the structural question — carbaryl D and both malathion and diazinon treatments; the narcotic windows (8–11 d) fall short of the 24 d their class needs, carbaryl E falls short of its 7.9 d, and the three uncoupler treatments measure rates near zero, leaving nothing for the structures to disagree about (§S3.2). What most fail is the window, not the controls: thirty-four have controls, sixteen leave a window long enough. The interval series of ref 5, the one neonicotinoid design with survival counts, measured concentrations in every replicate, and under the same rule eleven of its fifteen treatments meet all four; survival falls across the window in eight of them, in four of the five species, and stays flat in both *P. minutissima* treatments (§S7). These declines do not decide the structure. The rule opens the window two days after exposure, the allowance the diazinon cascade needs; for imidacloprid the upstream half-life is 1.53 d in *D. magna* (§4.3) and at least 10 d in *C. dipterum* (§S8). Opened three *D. magna* half-lives after exposure, seven treatments qualify and four still decline; at the *C. dipterum* bound, three half-lives exceed the 21-day test. The companion’s carry-over ratio cannot be formed on this series, whose intervals are shorter than the recovery they need.^4^

## 5. Discussion and Environmental Implications

The clean-water test measures the damage model, not the hazard structure. The same test gave opposite answers to one compound in one species, and what differed was the window (§4.4); across the archive it sorts by compound, not by hazard structure (§4.5). For *D. magna* the peak-based structure fails on a single-stage trajectory and passes once damage formation carries a persisting state: both structures read a trajectory the framework supplies, and here it supplied one stage where the data need two.^4^

The inference from deaths after exposure ends is valid; what two decades of use missed is that its premise — damage falling while the counts are taken — is supplied by the fitted damage model, not observed in the animal. A verdict therefore rests on two choices — hazard structure and damage model — and the observation cannot separate them; the companion’s test rests on the same fitted damage model.^4^ Because the verdict turns on the depth of that model, which depends on how long the lesion persists and how long the animals were watched (§4.3), assignments differing only in compound or window length have looked like disagreements about mechanism.

What the framework cannot express shows up in the neonicotinoids. Delivering a fixed exposure as pulses further apart increases mortality in one of five species, and no hazard structure and repair law within the framework’s kinetics reproduce that ordering (Proposition 2, §S10). The slow *de novo* synthesis of the target enzyme proposed for diazinon recovery in *D. magna*^29^ is a further non-negative repair term and so is covered by Proposition 2; an eroding tolerance is not, and §S13 excludes it separately (§4.6).

What remains is the non-monotone state of §4.6, which the framework does not carry and an energy budget describes. Two measurements of energy use support that reading, and a feeding manipulation would separate an energetic explanation from a damage-cascade one (§S13).

Spreading a fixed load over a longer period is what lengthening an application interval does, so a framework that cannot express the ordering fails the comparison regulators most often make. The framework’s predictions for such patterns are unvalidated, not conservative: a conservative model errs on the safe side, an unvalidated one in a direction not determined. Assessment for neonicotinoids should not rest on a choice between the reduced variants; it needs a model coupling survival to the sublethal state.

### What this does not show

The bound of §4.6 holds for convex repair, and §S10.2 shows what concave repair does at rates this archive does not reach. §4.3 rests on one series and four parameters, and the out-of-sample check withholds a treatment, not a dataset (§S14). The persisting state is inferred from survival, not measured, though it falls between two measured rates, elimination and receptor dissociation (§S11.1).^12^ The reversal is established for one of five species, and Propositions 2 and 3 assume equal pulses. None of these limits touches the design census, which uses counts alone.

### Implications for assessment

#### Fewer designs can bear on the structural question than the published verdicts imply

Five of the eight published assignments rest on a series in which most treatments meet §4.7 as the archive applies it, but none on a design whose window has been shown to open after the damage maximum for its species (§4.7); thirteen of thirty-six treatments in the archive meet §4.7, and five of nineteen studies outside it (§S5). What a guideline can adopt is the class window itself: 6.5 d for an organophosphate, 7.9 d for a carbamate (Table S5).

#### What can be changed in the aquatic guidance now

The specification of §4.7 needs no new measurements: a minimum window set from the compound class’s own decline rate, imposed as concurrent controls already are — the guidance requires the controls but not the window.^3^ The same is true of the damage model. Whether a stage between exposure and death must be carried is settled before fitting, by its measured rate constant against the test length (§4.3). A model chosen that way costs one parameter and removes the commonest way of crediting the hazard structure with what the damage model did. Neither recommendation depends on which hazard structure is chosen.

#### How to read a qualifying design

A qualifying design returns one of two outcomes, neither a verdict on the hazard structure. If survival declines, damage was either still rising or the peak-based structure is wrong, and §4.3 shows the first is not unusual. A decline licenses a structural conclusion only where the damage model has been shown deep enough, by refitting with an added stage (§S9.5, §S11). If survival is flat, the peak-based structure is not refuted and the lesion is short-lived; the rate separates the two classes of lesion (§4.5). Either way the rate is the datum worth reporting: it sets the window for the next compound in that class.

#### What a peak-based default would cost

Survival becomes a one-way instrument (Proposition 1): it records the running maximum and is flat while damage falls, so recovery between applications cannot be read from the counts, and any statement about damage must come from a design meeting §4.7.

#### How the published assignments should be re-read

A model comparison within the reduced framework says nothing about mechanism: published assignments report the adequacy of a single-stage damage model (§4.3). They are not invalidated; what changes is what they are evidence for. What settles the question is not a better fit but a quantity one structure cannot produce: the companion tests the carry-over ratio, which a cumulative hazard cannot raise above unity, and finds it at or above unity in fourteen of fifteen comparisons, seven with intervals excluding unity.^4^

Assessment gains more from measuring how long the lesion persists than from choosing between the structures.

## 6. Associated Content

### Supporting Information

The Supporting Information is available with this preprint.

- Statements of Propositions 1–3 with proofs of Propositions 1 and 3; data sources and post-exposure window census; clean-water tests, decline rates and compound-level tests; recovery ratios for three neonicotinoids; model configurations, fits, numerical checks of both dose metrics, out-of-sample prediction and robustness checks; script inventory; references cited only in the Supporting Information; Tables S1–S42 and Figures S1–S8 (PDF)

### Data and code availability

Reference 4 supplies the classification and the companion results quoted in §4.3–§4.7 and §5, each cited at the point of use; every other number here is computed from the archived scripts. All data are from published sources listed in Table S1. Analysis scripts reproducing every reported number and figure, including the parsers for every spreadsheet layout and the digitised coordinates extracted from ref 6, are archived with the data at Zenodo (https://doi.org/10.5281/zenodo.23488395), ^30^ a deposit the Supporting Information joins on acceptance. The general proof of Proposition 2, and of the results cited in §S10, is in Technical Report T0, version 1.0 (Zenodo, 2026; https://doi.org/10.5281/zenodo.23356185), cited as ref 35 in the Supporting Information.

## Author Information

### Notes

No new experiments were performed and no animals were handled by the authors; all analyses use published data. The authors declare no competing financial interest.

### Author contributions

Jong-Hyeon Lee: conceptualization, methodology, software, formal analysis, investigation, data curation, writing — original draft, writing — review and editing, visualization. Chan Young Joe: validation, formal analysis, writing — review and editing.

## 8. Acknowledgments

This work was supported by the Korea Environmental Industry & Technology Institute (KEITI) with funding from the Korea Ministry of Environment (MOE) through the projects “Development of chemical emission factors for consumer products” (RS-2020-KE000904) and “Development of cumulative exposure assessment and prediction for biocide” (RS-2018-KE000099). AI-assisted tools were used during manuscript preparation for editorial refinement and for implementing, running and checking analysis code; all scientific content, analysis and interpretation are the responsibility of the authors.

