## Supporting Information for "The Test Used to Choose Between Pesticide Survival Models Tracks Lesion Persistence, Not Hazard Structure"

#### Contents


| Section | Tables | Figures |
|---|---|---|
| **Part I. Theory** | | |
| **S1** The Three Propositions: What They Say, What They Assume, and How They Are Proved | — | — |
| **Part II. Data and Designs** | | |
| **S2** Data Sources and Extraction | S1, S2 | — |
| **S3** The Post-Exposure Window Census | S3, S4, S5 | S1 |
| **Part III. What the Test Measures** | | |
| **S4** The Clean-Water Test, Treatment by Treatment | S6, S7, S8, S9, S10, S11, S12 | S2 |
| **S5** The Rate Outside the Archive | S13 | — |
| **S6** The Two Diazinon Experiments: Window Placement, Test Results, and Interval Ordering | S14, S15 | S3 |
| **S7** The Five-Species Imidacloprid Interval Series: Survivor Counts, Immobilisation, and the Window after the Second Pulse | S16, S17, S18 | S4 |
| **S8** How Much Three Neonicotinoids Recover Between Pulses, Read from Survival Counts Alone | S19 | S5 |
| **Part IV. Model Fits and Controls** | | |
| **S9** Materials and Methods for the Model Fits | S20, S21, S22, S23 | S6 |
| **S10** Numerical Verification of Both Dose Metrics | S24, S25, S26, S27, S28 | S7, S8 |
| **S11** The *Daphnia magna* Series: Fits, Parameters, and What Each Extension Contributes | S29, S30, S31, S32, S33 | — |
| **S12** The Two-Stage Extension Applied to Two More Neonicotinoid Series and to an Uncoupler | S34 | — |
| **S13** Why an Eroding Tolerance Cannot Produce the Interval Reversal, and What Could | — | — |
| **S14** Out-of-Sample Prediction: Each Treatment Predicted from the Other Two | S35, S36 | — |
| **S15** Robustness: Digitisation Error, Identifiability, and the Window Rule | S37, S38, S39, S40, S41 | — |
| **Part V. Availability and References** | | |
| **S16** Code and Data Availability | S42 | — |
| **S17** References Cited Only in the Supporting Information | — | — |


Tables: 42 · Figures: 8

### Part I. Theory

#### S1. The Three Propositions: What They Say, What They Assume, and How They Are Proved

The main text states three results about the peak of damage in words. This section restates them for a reader without training in analysis, gives their exact statements, and proves Propositions 1 and 3 in full. The general proof of Proposition 2 is Theorem 5 of a deposited technical report;[35] its idea is set out here, and the proposition is also checked numerically (§S10).

**One picture for all three.** Think of damage as the water level in a tank. Exposure is a tap that fills it, and repair is a drain that empties it. The drain may run faster or slower as the level changes, in any way at all, but it never adds water. In symbols, $\mathrm{d}D/\mathrm{d}t = u - R(D)$, with an input $u \geq 0$ and a repair term $R(D) \geq 0$. A peak-based hazard reads only the highest mark the water has left on the wall, the running maximum $M(t) = \max_{\tau \leq t} D(\tau)$. A cumulative hazard reads how long the water has stood above a line. All three propositions are statements about that high-water mark, and none of them assumes a functional form for $R$ or any parameter value.

- Proposition 1 (freezing): once exposure stops, the high-water mark stops moving, so a peak-based hazard predicts no further deaths. It assumes only that repair never adds damage, it is proved below in two lines, and it is what the clean-water observation of §2 and the parameter-free comparison of §4.2 rest on.
- Proposition 2 (spacing cannot raise the peak): moving two equal pulses further apart never raises the high-water mark, for any repair law meeting the conditions stated below, provided the input from a single pulse fades in the way a chain of first-order steps produces. Its idea is given below and its general proof in ref 35; §S10 checks it numerically, under ten repair laws at short separations and six over a wider range. It is what places the interval reversal of §4.6 outside the framework.
- Proposition 3 (spacing cannot halve the peak): with convex repair, moving two equal pulses apart lowers the high-water mark by at most half. It is proved below in two steps and checked in Tables S27 and S28. It bounds the recovery a peak-based metric can express (§4.6), and, as shown below, it cannot be tested directly on survival counts.

Propositions are numbered here independently of ref 35, which carries its own correspondence table; there, Proposition 2 is Theorem 5 and Proposition 3 is Theorems 6a and 6b.

##### *S1.1 Proposition 1: once exposure stops, the peak is frozen*

**Proposition 1 (freezing).** *Let $dD/dt = u(t) - R(D)$ with an input $u \geq 0$ and any repair function $R \geq 0$, and suppose $u(t) = 0$ for $t \geq t_0$. Then $dD/dt \leq 0$ on $[t_0, \infty)$, so $D$ is non-increasing there, $M(t) = \max_{\tau \leq t} D(\tau)$ is constant, and any hazard depending on the trajectory only through $M$ is frozen.*

*Proof.* For $t \geq t_0$, $\mathrm{d}D/\mathrm{d}t = -R(D) \leq 0$ because $R \geq 0$ by assumption. A non-increasing function attains its supremum at the left endpoint of the interval, so $M(t) = M(t_0)$ for all $t \geq t_0$. Any hazard that depends on the trajectory only through $M$ is then constant. □

**Neither the functional form of *R* nor any of its parameters enters**, so the result covers both repair laws of the classification.[4]

**What Proposition 1 licenses.** A peak-based hazard has no memory of how long damage stayed high, only of how high it got. Once the compound is gone the maximum cannot be exceeded, so the model predicts nothing beyond background mortality — whatever the repair law and whatever the parameter values. That is what makes a death in clean water look like a refutation, and it is also why the refutation depends entirely on the claim that the input has stopped, which §2 shows is not observed but supplied by the fitted damage model.

##### *S1.2 Proposition 2: moving two equal pulses apart cannot raise the peak*

**The claim in words.** Deliver a fixed exposure as two equal pulses. Moving the pulses further apart never raises the highest damage reached, for any repair law meeting the conditions of §S1. A peak-based model therefore has no setting in which wider spacing is more lethal.

**Why it is not obvious.** The designs cannot be compared hour by hour. When the second pulse arrives late, damage at that moment can be higher than in a design whose pulses were closer together, because the closer design has already passed its peak and is recovering. Some other way of lining the designs up is needed.

**The idea of the proof.** Instead of comparing the designs at the same time, compare them at the same amount of exposure received so far. Every design delivers the same total, so each point on this cumulative scale is reached by every design. At any such point, the design with the pulses further apart has received that amount more slowly. Repair has therefore had more time to act, and its damage is no higher. A comparison argument makes this exact, and taking the highest point of each trajectory gives the result.

**Where log-concavity enters.** The step that needs care is the claim that the more separated design always receives exposure more slowly at the same cumulative amount. That holds when the input from a single pulse, once it starts to fade, fades at a relative rate that never slows down. This property is log-concavity, condition (V3) below. A rectangular exposure passed through any chain of first-order steps — uptake, then one or more damage stages — always has it. An input that fades quickly at first and then slowly lacks it, as happens when a compound is eliminated by a fast and a slow pathway in parallel; there the conclusion can fail, and §S10.1 shows by how much.

**Proposition 2 (peak monotonicity under pulse separation).** *Let a fixed dose be delivered as two identical rectangular pulses whose start times are separated by $\Delta \geq 0$, overlapping designs included, and let the damage input $v$ generated by a single pulse satisfy: (V1) $v = 0$ for $t \leq 0$, and $v > 0$ and continuous on $(0,\infty)$; (V2) $V = \int_0^\infty v < \infty$; (V3) $\log v$ concave on $(0,\infty)$. Let $D$ solve $dD/dt = u_\Delta(t) - R(D)$ with $u_\Delta(t) = v(t) + v(t-\Delta)$, $D(0) = 0$, where $R \geq 0$, $R(0) = 0$ and $R$ is locally Lipschitz. Then*

$$M(\Delta) \equiv \sup_t D_\Delta(t)$$

*is non-increasing on $[0, \infty)$, so $M$ is largest at complete superposition, $\Delta = 0$.*

**Where the hypotheses come from.** The additive form $u_\Delta = v(t) + v(t-\Delta)$ is not an assumption about the two-pulse system but a consequence of linearity *upstream* of damage: toxicokinetics and any intermediate damage stages are linear, time-invariant and start at rest, so the input generated by two pulses is the sum of two shifted copies of the input generated by one. Superposition stops at the damage equation, because $R$ is not assumed linear, which is why the two-pulse trajectories have to be compared directly. (V1) and (V2) say that a pulse delivers a finite total input, so that every design can be compared at the same cumulative input. (V3) is automatic for the models used here: a rectangular exposure passed through a series of first-order stages is a convolution of log-concave kernels, and convolution preserves log-concavity, so single-stage, two-stage and longer cascades all qualify. On the repair side, $R \geq 0$ makes repair remove damage rather than add it, $R(0) = 0$ keeps damage non-negative, and local Lipschitz continuity makes solutions unique, which the comparison step needs. Monotonicity of $R$ is not assumed, so repair that slows when damage is severe — when damage impairs the repair machinery itself — is covered, as is threshold repair, $r[D-c]^+$, which is Lipschitz. Constant removal does not vanish at zero damage and so lies outside these conditions, but the conclusion holds for it as the limit of laws that satisfy them (ref 35, §1 and Theorem 5).

**Proof.** The complete argument is given in ref 35 (Theorem 5). For the specialist, its two steps are these. Writing $\sigma$ for cumulative input and $U_\Delta(\sigma)$ for the input rate at cumulative input $\sigma$, damage $y_\Delta(\sigma)$ obeys

$$\frac{\mathrm{d}y_\Delta}{\mathrm{d}\sigma} = 1 - \frac{R(y_\Delta)}{U_\Delta(\sigma)},$$

in which the first term says that input accumulates as damage and the second is the fraction lost to repair per unit of input received. Log-concavity of the single-pulse input is exactly what makes $U_\Delta(\sigma)$ non-increasing in $\Delta$ at each fixed $\sigma$: with $F(t) = \int_0^t v$ and the constraint $F(t) + F(s) = \sigma$, $t - s = \Delta$,

$$\partial_\Delta U = \frac{v(t)\,v(s)}{U}\Big[(\log v)'(t) - (\log v)'(s)\Big] \le 0,$$

so the older pulse loses at least as fast, in relative terms, as the younger one gains. A comparison argument in $\sigma$ then orders the trajectories, $y_{\Delta'} \le y_\Delta$ on $[0, 2V)$, and taking suprema gives the proposition. ■

**Which designs the theorem covers.** The single 48-hour pulse of §4.6 is two 24-hour pulses back to back ($\Delta/\tau = 1$, $\tau$ being the pulse duration), so its second pulse begins before the first has peaked; Theorem 5 of ref 35 covers that comparison as well as the seven- and fourteen-day designs. The proposition is also checked numerically at separations from 0 to 14 days under ten repair laws, threshold and non-monotone ones included; the designs of §4.6 have $\Delta/\tau = 1$, 7 and 14, all three inside that range, and $M$ is non-increasing throughout (§S10).

**How much of the log-concavity hypothesis is needed.** Two results mark its limits: one repair law needs only a single-peaked input, and without the condition a violation can be built.

**Linear repair never produces the reversal for a unimodal input.** For $R(D) = \kappa D$ the peak is non-increasing in the separation whenever the single-pulse input rises to one maximum and then falls, log-convex tails included, so a violation requires a repair law that is nonlinear over the range the trajectory visits. Proof: ref 35, Theorem 1.

**Without log-concavity the conclusion can fail, and this separates series kinetics from parallel kinetics.** If $\log v$ is strictly convex on an interval, a repair law that switches on in a thin band of damage can hold the less separated design at that level while the more separated one passes through, and the conclusion can fail. A *series* of first-order filters produces a difference of exponentials and is log-concave; a *parallel* mixture, $A_1 e^{-k_1 t} + A_2 e^{-k_2 t}$ with $A_1, A_2 > 0$ as arises from two-compartment distribution kinetics, is log-convex in the tail, because the fast component is exhausted first and the decay rate relaxes with time. The distinction matters when the toxicokinetics of a compound are described by distribution rather than by a cascade; §S10.1 shows how large a violation it can produce.

**What Proposition 2 licenses.** Splitting a fixed exposure and moving the halves apart can only lower the peak damage, for any repair law meeting the conditions of §S1. A peak-based model therefore has no setting — no repair function, no parameter value — that makes wider spacing more lethal. §4.6 reports the opposite in one of five species, survivors falling from 51 to 40 to 29 as the interval grows: that ordering is not a poor fit but an outcome the family cannot produce. What the proposition does not cover is a threshold that itself moves with time; §S13 closes that route separately, and by a different kind of argument.

**Why the cumulative metric is checked numerically instead.** The proof orders the designs at equal cumulative input, and that ordering does not carry over to quantities added up over time. Since $dt = d\sigma/U_\Delta$ and $U_\Delta$ is ordered in the opposing direction, the more separated design spends more time at each level of cumulative input; for a cumulative excess statistic $I(\Delta) = \int_0^\infty [D_\Delta - z]^+ \, dt$, with $z$ the hazard threshold, lower damage and longer time compete, and for some repair laws the second dominates. No proof is therefore available on that side, which is why §S10 verifies the cumulative metric numerically, for proportional and Michaelis–Menten repair, over 81 parameter combinations (Table S25) and as a continuous function of the separation (Figure S7). In the linear case the two endpoints of $I$ can still be compared: with $f$ the single-pulse damage trajectory, $I(0) = \int [2f - z]^+ dt \geq 2\int [f - z]^+ dt = \lim_{\Delta \to \infty} I(\Delta)$, with strict inequality whenever $f$ exceeds $z/2$ anywhere; monotonicity between the endpoints is verified numerically in §S10 rather than proved, and no violation was found.

##### *S1.3 Proposition 3: moving two equal pulses apart cannot halve the peak*

**The claim in words.** With convex repair, moving two equal pulses apart lowers the highest damage by at most half. However much the animal recovers between pulses, a peak-based metric cannot express more than a twofold reduction.

**Proposition 3 (a floor on the peak ratio).** *Under the conditions of Proposition 2, and for convex repair, for any two separations $\Delta' > \Delta \geq 0$,*

$$\frac{M(\Delta')}{M(\Delta)} \geq \frac{1}{2}.$$

*Proof.* Both steps rest on the comparison principle for a single differential equation: a solution that starts no lower and is driven no less stays no lower. Let $f$ be the damage trajectory produced by a single pulse, with peak $f_{\max}$.

*Step 1: two pulses never give more than twice the single peak.* A convex repair law with $R(0) = 0$ removes at least as much from a combined level as from its parts separately, $R(a+b) \geq R(a) + R(b)$ for $a, b \geq 0$. The sum of two single-pulse trajectories, $W(t) = f(t) + f(t-\Delta)$, therefore satisfies

$$W' = u_\Delta - R(f(t)) - R(f(t-\Delta)) \geq u_\Delta - R(W),$$

so $W$ can never fall below the two-pulse trajectory $D_\Delta$, which starts from the same value. Hence $M(\Delta) \leq \max_t W(t) \leq 2f_{\max}$ for every $\Delta$. For proportional repair the inequality is an equality and $W$ is the trajectory itself.

*Step 2: the second pulse alone reaches at least the single peak.* When the second pulse begins, damage is at least zero, and from then on the input is at least that of the second pulse alone. A trajectory that starts no lower and receives no less input stays no lower, so $D_{\Delta'}(\Delta'+s) \geq f(s)$ for all $s \geq 0$, and $M(\Delta') \geq f_{\max}$. This step uses no convexity.

Dividing the two bounds gives $M(\Delta')/M(\Delta) \geq f_{\max}/(2f_{\max}) = 1/2$. The same argument, with its regularity conditions, is ref 35, Theorems 6a and 6b. □

**Why convexity is needed.** Step 1 fails when repair is concave. A concave law removes less from a combined level than from its parts, so two overlapping pulses can push damage above twice the single peak, and the floor can be breached (§S10.2). Step 2 holds for any non-negative repair law.

**What Proposition 3 licenses.** However long the interval, the second pulse alone sets a maximum of at least $f_{\max}$, so a peak-based dose metric cannot express more than a twofold reduction in damage between two equal pulses; the bound is attained in the limit of complete recovery between pulses and of complete overlap. It is the peak-based structure's own falsifiable limit — the counterpart on that side to the carry-over ratio on the cumulative side — and §4.6 reports that it has never been tested. The paragraph below shows why survival counts alone cannot test it, and what a design would have to supply.

**Relation to the companion.** The companion states the same bound asymptotically, as the residual from the first pulse approaches the height of the second.[4] The form proved here is uniform: it holds for every separation $\Delta' > \Delta \geq 0$, overlapping designs included, which is what §4.6 needs — an observed ratio has to be compared with a bound that holds at the separation actually used, not with one attained only in a limit.

**Why the bound is not directly observable.** Survival under a peak-based hazard gives $-\ln S = k(M-\theta)$, so the measurable ratio is

$$\frac{-\ln S(\Delta')}{-\ln S(\Delta)} = \frac{M(\Delta')-\theta}{M(\Delta)-\theta} < \frac{M(\Delta')}{M(\Delta)}$$

whenever $M(\Delta') < M(\Delta)$ and $0 < \theta < M(\Delta')$.

An observed ratio below one half is therefore not a violation. Testing the bound requires either a single-pulse arm, which fixes $\theta$ through $M = f_{\max}$, or a design in which the threshold is small relative to peak damage.

##### *S1.4 Terms used in this section*

**Running maximum**, $M(t)$: the highest damage reached up to time $t$. It can rise or stay level, never fall.

**Log-concave input**: an input whose logarithm is concave. In words, once it starts to decline, its relative rate of decline never decreases. A chain of first-order steps always produces one; a mixture of fast and slow exponential decays does not.

**Convex and concave repair**: convex repair removes damage at a rate that grows at least in proportion to damage, proportional repair being the boundary case; concave repair saturates, as Michaelis–Menten repair does.

**Comparison principle**: for a single differential equation, a solution that starts no lower and is driven no less stays no lower. It requires that solutions be unique, which local Lipschitz continuity of $R$ guarantees.

**Supremum**: the least upper bound of a trajectory, equal to its maximum whenever the maximum is reached.

---

### Part II. Data and Designs

#### S2. Data Sources and Extraction

All data are published. No new experiments were performed and no animals were handled by the authors. The table below covers the series this paper reanalyses. Sections §S4.1, §S5 and §S11.1 additionally quote published values — rate constants, recovery times and effect percentages — from studies listed in the references; those are citations rather than extractions, and no data were read from their figures.

**Table S1. Sources and how each was read.**

| Source | Content used | Extraction |
|---|---|---|
| Li *et al.* 2021, main text and SI | Three pulsed treatments of *D. magna*; toxicokinetic and toxicodynamic parameters; model code | Survival from the mortality percentages annotated in Figure 2 and from statements in the text, with two points read from the plotted curve (Table S2); parameters from Table 2 of that paper; model structure from the OpenModel listing in its SI |
| Ashauer *et al.* 2015, data supplement | Daily survival counts, concurrent controls and measured exposure profiles for thirteen compounds in *G. pulex* (a fourteenth sheet reproduces the chlorpyrifos experiment of ref 21; see §S3) | Read directly from the data file by the two parsers described in §S3 and listed in §S16 |
| Ashauer *et al.* 2010 | Three pulsed diazinon treatments, intervals 2, 7 and 15 d, with concurrent controls | Counts and measured exposure read from the data file of that paper, which the 2015 data supplement reproduces; windows placed by the fitted trajectory of §S6 |
| Focks *et al.* 2018, main text | Endpoint survivor counts for five species under three exposure splittings, and for three neonicotinoids under two intervals | Design and endpoint totals from the text and figures; the counts themselves from ESM5 (next row) |
| Focks *et al.* 2018, ESM5 | Raw daily dead / alive / immobile counts, replicate-resolved | Read directly |
| Van den Brink *et al.* 2016 | Acute and chronic effect concentrations, two generations | Read from the tables |

**The three pulsed treatments of Li *et al.*, and where each point comes from.** Seven replicates of ten animals per treatment; control survival 76 ± 5% at 22 d. Most of the survival series is stated numerically in the source rather than shown only in the figure: Figure 2 there annotates the mortality within each interval as a percentage, and the text gives two further values directly. Table S2 sets out which is which.

**Table S2. Provenance of the seventeen survival points.**
"Stated" means the value follows from a percentage printed in Figure 2 of the source or from a sentence in its text; "read" means it was taken from the plotted curve alone; the three day-0 rows are fixed by the design of the test.

| Trt | Day | *S* | Source |
|---|---|---|---|
| A | 0 | 1.00 | start of test |
| A | 3 | 0.65 | 35% mortality in 3 d after first pulse (stated) |
| A | 6 | 0.13 | 80% mortality in 3 d after second pulse (stated) |
| A | 11 | 0.085 | 87% mortality in 8 d after second pulse (stated) |
| **A** | **22** | **0.080** | **read** |
| B | 0 | 1.00 | start of test |
| B | 3 | 0.68 | 32% mortality in 3 d after first pulse (stated) |
| B | 8 | 0.32 | 68% mortality in 8 d after first pulse (stated) |
| B | 11 | 0.00 | 100% mortality in 3 d after second pulse (stated) |
| **B** | **22** | **0.000** | **read** |
| C | 0 | 1.00 | start of test |
| C | 1 | 0.89 | text: survival after the first 1-d exposure was 89 ± 7% |
| C | 3 | 0.65 | 35% mortality in 3 d after first pulse (stated) |
| C | 8 | 0.32 | 68% mortality in 8 d after first pulse (stated) |
| C | 10 | 0.24 | text: dropped to 24 ± 11% on the 10th day |
| C | 16 | 0.24 | text: no mortality between day 9 and the second exposure |
| C | 19 | 0.00 | 100% mortality in 3 d after second pulse (stated) |

Fifteen of the seventeen points follow from the source text or the design of the test; the two read from the plotted curve alone (treatments A and B on day 22) lie at the floor of series already flat on day 11.

**Digitisation error.** The two read points carry an estimated error of $\pm 0.03$ in surviving fraction, about two animals in seventy; §S15.1 reports the bootstrap. The parameter-free comparison of §4.2 uses values stated in the text (89% on day 1, 24% on day 10) and turns on a difference of 0.44 in surviving fraction, fifteen times that error. The window census, the clean-water tests, the interval reversal and the compound recovery timescales all use counts stated numerically in their sources.

---

#### S3. The Post-Exposure Window Census

The clean-water test needs a window between the last exposure and the end of survival observation. Where the data file records both, the window was taken to open two days after measured exposure last exceeded 5% of its maximum — the allowance a multi-stage cascade needs to reach its peak; the three-compartment model of §S6 takes 1.6 to 1.9 days — and to close when observation ends.

The data file records exposure in six distinct sheet layouts, and two sheets carry two survival blocks each rather than one; a parser that stops at the first block misses the second experiment entirely. The parser of §S16 reads all six.

**Table S3. Post-exposure window by treatment.** Rows are from the thirteen-compound archive of ref 17 except the pentachlorophenol rows, which are from the chlorpyrifos–pentachlorophenol series of ref 21; the pentachlorophenol sheet of ref 17 is a separate experiment and appears in Table S4 instead. The archive data file carries a fourteenth sheet, chlorpyrifos, which reproduces the experiment of ref 21 and is counted with that series rather than with the archive; its treatments appear in Table S8. Bold rows have a window of at least one day, concurrent controls and animals alive when it opens, and are tested in §S4.

| Compound | Mode of action | Treatment | Observation ends (d) | Exposure ends (d) | Window (d) |
|---|---|---|---|---|---|
| **1,2,3-trichlorobenzene** | **narcosis** | **A** | 19 | 6.0 | **11** |
| **1,2,3-trichlorobenzene** | **narcosis** | **B** | 19 | 9.0 | **8** |
| **1,2,3-trichlorobenzene** | **narcosis** | **C** | 19 | 13.0 | **4** |
| **2,4-dichloroaniline** | **narcosis** | **A** | 19 | 6.0 | **11** |
| **2,4-dichloroaniline** | **narcosis** | **B** | 19 | 9.0 | **8** |
| **2,4-dichloroaniline** | **narcosis** | **C** | 19 | 13.0 | **4** |
| pentachlorophenol | uncoupling | C: I, II, III | 10 | 9.0 | 0 |
| pentachlorophenol | uncoupling | D: I | 20 | 16.0 | 2 |
| pentachlorophenol | uncoupling | D: II | 20 | 19.0 | 0 |
| pentachlorophenol | uncoupling | D: III | 20 | 20.0 | 0 |
| **2,4,5-trichlorophenol** | **uncoupling** | **A** | 10 | 4.0 | **4** |
| **2,4,5-trichlorophenol** | **uncoupling** | **B** | 10 | 7.0 | **1** |
| 2,4,5-trichlorophenol | uncoupling | C | 10 | 10.1 | 0 |
| **DNOC** | **uncoupling** | **A** | 15 | 4.0 | **9** |
| **DNOC** | **uncoupling** | **B** | 15 | 6.0 | **7** |
| **DNOC** | **uncoupling** | **C** | 15 | 10.0 | **3** |
| **2,4-dichlorophenol** | **uncoupling** | **A** | 19 | 6.0 | **11** |
| **2,4-dichlorophenol** | **uncoupling** | **B** | 19 | 9.0 | **8** |
| **2,4-dichlorophenol** | **uncoupling** | **C** | 19 | 13.0 | **4** |
| **aldicarb** | **carbamate** | **A** | 10 | 4.0 | **4** |
| **aldicarb** | **carbamate** | **B** | 10 | 7.0 | **1** |
| aldicarb | carbamate | C | 10 | 10.1 | 0 |
| **carbofuran** | **carbamate** | **A** | 10 | 4.0 | **4** |
| **carbofuran** | **carbamate** | **B** | 10 | 7.0 | **1** |
| carbofuran | carbamate | C | 10 | 10.0 | 0 |
| **carbaryl** | **carbamate** | **A, B, C** | 15 | 11.0 | **2** |
| **carbaryl** | **carbamate** | **D** | 22 | 11.0 | **9** |
| **carbaryl** | **carbamate** | **E** | 22 | 15.0 | **5** |
| **malathion** | **organophosphate** | **A** | 22 | 7.2 | **13** |
| **malathion** | **organophosphate** | **B** | 22 | 10.0 | **10** |
| **malathion** | **organophosphate** | **C** | 22 | 17.0 | **3** |
| **diazinon** | **organophosphate** | **A** | 22 | 4.0 | **16** |
| **diazinon** | **organophosphate** | **B** | 22 | 9.0 | **11** |
| 4-nitrobenzyl chloride | reactive | A | 6 | 1.0 | 3 |
| Sea-Nine | reactive | A | 14 | 14.1 | 0 |

Thirty-nine treatments; the median window is four days, twenty-three treatments reach three days or more, and nine have no window at all. Ten compounds leave at least one window that can be tested; the 4-nitrobenzyl chloride treatment has none alive when its window opens. The pentachlorophenol and chlorpyrifos sheets of the archive belong to the 2007a series and are counted there (Table S4, Table S8).

##### *S3.1 Per-treatment detail behind the design criterion*

The four requirements are stated in §4.7 of the main text; Table S4 summarises the tally by compound, and Table S3 gives the per-treatment windows on which the third requirement turns.

**Table S4. The four design requirements of §4.7, applied to every treatment in the archive.** A treatment counts toward a requirement only if the data file records what the requirement needs.

| Compound | Treatments | Exposure profile | Concurrent controls | Window ≥ 4.6 d | ≥ 10 alive | All four |
|---|---|---|---|---|---|---|
| 1,2,3-trichlorobenzene | 3 | 3 | 3 | 2 | 3 | **2** |
| 2,4-dichloroaniline | 3 | 3 | 3 | 2 | 3 | **2** |
| 2,4-dichlorophenol | 3 | 3 | 3 | 2 | 3 | **2** |
| pentachlorophenol | 2 | 2 | 0 | 2 | 0 | 0 |
| 2,4,5-trichlorophenol | 3 | 3 | 3 | 0 | 2 | 0 |
| DNOC | 3 | 3 | 3 | 2 | 2 | **1** |
| aldicarb | 3 | 3 | 3 | 0 | 2 | 0 |
| carbofuran | 3 | 3 | 3 | 0 | 2 | 0 |
| carbaryl | 5 | 5 | 5 | 2 | 2 | **2** |
| malathion | 3 | 3 | 3 | 2 | 3 | **2** |
| diazinon* | 3 | 3 | 3 | 2 | 2 | **2** |
| 4-nitrobenzyl chloride | 1 | 1 | 1 | 0 | 0 | 0 |
| Sea-Nine | 1 | 1 | 1 | 0 | 0 | 0 |
| **total** | **36** | **36** | **34** | **16** | **24** | **13** |

* Read by the branch of the census parser written for the layout of the diazinon data file (`s03_window_census.py`, §S16) rather than by the general one.

The thirteen treatments meeting all four requirements are treatments A and B of 1,2,3-trichlorobenzene, 2,4-dichloroaniline, 2,4-dichlorophenol and malathion (8–13-day windows, 10–52 alive at opening), DNOC treatment B (7-day window, 19 alive), carbaryl treatments D and E (9 and 5 days, 29 and 32 alive), and both diazinon treatments (16 and 11 days, 29 and 22 alive).

**One consequence for the matched comparison of §4.5.** The group-size requirement removes DNOC treatment A from the matched comparison of §4.5: seven animals are alive when its window opens, too few to detect a halving of survival, so treatment B is used instead — seven days rather than nine, nineteen animals rather than seven, and no losses across the window. Ten is the smallest group in which a halving of survival reaches significance against a control that loses none (Fisher's exact test, two-sided: $p = 0.033$ for 10 → 5 against 10 → 10; $p = 0.077$ for 8 → 4).

##### *S3.2 A class-specific window requirement, obtained without fitting*

The 4.6 d of §4.7 is the recovery timescale fitted to imidacloprid in *Daphnia magna*, so applying it to the *Gammarus pulex* archive imports a number from a model under dispute. The same requirement can be set from counts alone. What has to be large is the product of the decline rate and the window length; the product at which 30% of the survivors are lost — the smallest loss twenty animals can detect — is 0.35, since $1 - e^{-0.35} = 0.30$. Dividing by the class medians of Table S8 gives the window each class demands (Table S5).

**Table S5. The design requirement by compound class.** Rates are the compound-pooled medians of Table S8; the requirement is 0.35 divided by the rate. No fitted quantity enters. The neonicotinoid row carries no requirement: its rate, from the interval series of §S7, is measured on windows that may open before damage peaks (§4.7), and receptor dissociation exceeds 8 d (ref 12).

| Class | Median $h_{post}$ ($d^{-1}$) | Window required (d) |
|---|---|---|
| organophosphate | 0.054 | 6.5 |
| carbamate | 0.045 | 7.9 |
| narcotic | 0.015 | 23.9 |
| uncoupler | 0.003 | 110.8 |
| neonicotinoid | (0.052) | not set |

Re-tallying the census against these numbers rather than against a single 4.6 d moves the count from thirteen treatments to five: carbaryl D (9 d against 7.9), both malathion treatments (13 and 10 d against 6.5) and both diazinon treatments (16 and 11 d) clear their class requirement; carbaryl E (5 d) and the four narcotic treatments (8–11 d against 24) do not, and the three uncoupler treatments cannot, because the rate an uncoupler shows is near zero. That last case is not a failure of design. DNOC treatment B measures its rate precisely — the interval of Table S8 excludes anything above 0.010 $d^{-1}$ — and it is the measurement §4.5 uses. What it cannot do is discriminate between the hazard structures, because there is no decline for the structures to disagree about. The two counts answer different questions: thirteen designs can measure the post-exposure decline, and five of those can bear on the structural question. Reproduced by `s03_class_window.py` (§S16).

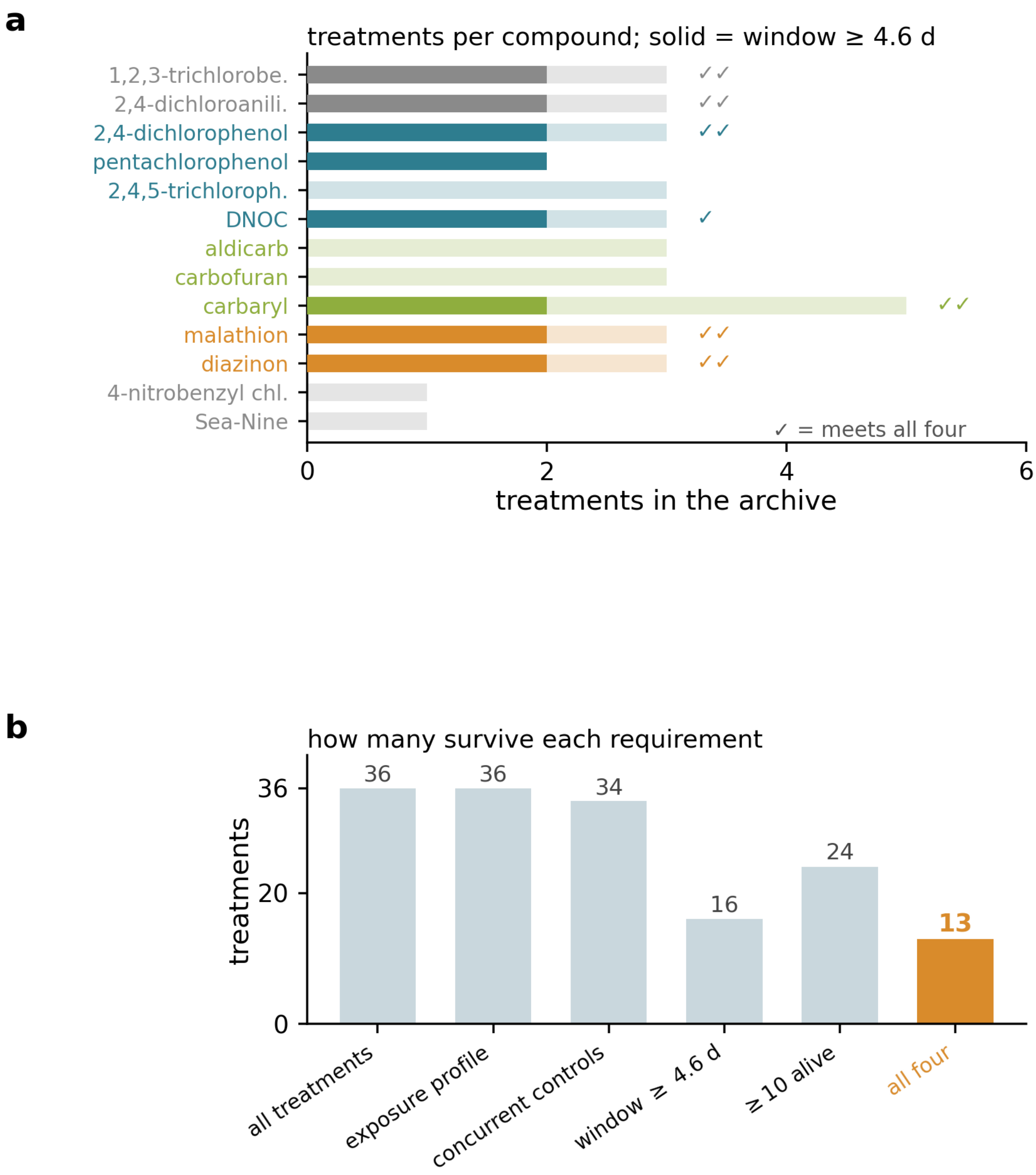


**Figure S1. How many published designs can bear on the question.** (a) The thirteen compounds of the pulsed archive: pale bars give the treatments each contributes, solid bars the number leaving a window of 4.6 d or more, and ticks the treatments meeting all four requirements of §4.7. Colour marks the mode of action. (b) The four requirements applied in turn: 34 of 36 treatments have concurrent controls but only 16 leave a window of 4.6 d or more, and 13 meet all four. The archive holds no neonicotinoid; the one neonicotinoid interval series with survival counts is scored in §4.7 and §S7.

The class windows of Table S5 are detectability windows — long enough for the product of rate and window to be resolved in counts. The companion's Condition 1 is a divergence window — long enough for the two structures to separate dynamically. Diazinon treatment B (11 d) clears the first and falls far short of the second, which for diazinon is one $t_{95}$ of 167 d; the two requirements answer different questions and both must be read.

### Part III. What the Test Measures

#### S4. The Clean-Water Test, Treatment by Treatment

Survival is control-corrected as $(n_{trt}/N_{trt})/(n_{ctl}/N_{ctl})$, evaluated at the opening and close of the window; the correction assumes that background mortality enters multiplicatively, the assumption the fitted models also make. The change is tested by Fisher's exact test on a 2 × 2 table whose rows are the treatment and its concurrent control and whose columns are the animals alive at the close of the window and those that died within it, both counted among the animals alive when it opened. It is not a comparison of two time points within one cohort, which the overlap of the two counts would not permit. Because twenty-eight tests are reported, $p$-values are accompanied by Benjamini–Hochberg false-discovery-rate $q$-values computed across the whole set.

**Table S6. Every window of at least one day with concurrent controls.** Group sizes are treatment / control. Trajectories for the three diazinon treatments are in Figure S3 and for the five carbaryl treatments in Figure S2.

| Compound | Mode of action | Trt | Window (d) | Alive | $N_{trt}/N_{ctl}$ | Corrected survival | $p$ | $q_{BH}$ |
|---|---|---|---|---|---|---|---|---|
| 2,4,5-trichlorophenol | uncoupler | A | 6–10 | 15 → 14 | 70/90 | 0.264 → 0.261 | 1.00 | 1.00 |
| 2,4,5-trichlorophenol | uncoupler | B | 9–10 | 20 → 20 | 70/90 | 0.373 → 0.373 | 1.00 | 1.00 |
| DNOC | uncoupler | A | 6–15 | 7 → 7 | 70/60 | 0.111 → 0.120 | 1.00 | 1.00 |
| DNOC | uncoupler | B | 8–15 | 19 → 19 | 70/60 | 0.313 → 0.326 | 1.00 | 1.00 |
| DNOC | uncoupler | C | 12–15 | 13 → 12 | 70/60 | 0.218 → 0.206 | 0.37 | 0.58 |
| 2,4-dichlorophenol | uncoupler | A | 8–19 | 52 → 47 | 70/70 | 0.852 → 0.810 | 0.47 | 0.69 |
| 2,4-dichlorophenol | uncoupler | B | 12–19 | 43 → 40 | 70/70 | 0.729 → 0.690 | 0.31 | 0.57 |
| **2,4-dichlorophenol** | **uncoupler** | **C** | **16–19** | **46 → 42** | 70/70 | 0.793 → 0.724 | **0.035** | 0.124 |
| 1,2,3-trichlorobenzene | narcotic | A | 8–19 | 33 → 24 | 70/70 | 0.579 → 0.490 | 0.16 | 0.35 |
| 1,2,3-trichlorobenzene | narcotic | B | 12–19 | 16 → 13 | 70/70 | 0.296 → 0.265 | 0.37 | 0.58 |
| 1,2,3-trichlorobenzene | narcotic | C | 16–19 | 25 → 24 | 70/70 | 0.490 → 0.490 | 1.00 | 1.00 |
| 2,4-dichloroaniline | narcotic | A | 9–19 | 16 → 13 | 70/70 | 0.258 → 0.224 | 0.15 | 0.35 |
| 2,4-dichloroaniline | narcotic | B | 12–19 | 10 → 8 | 70/70 | 0.167 → 0.138 | 0.095 | 0.27 |
| 2,4-dichloroaniline | narcotic | C | 16–19 | 29 → 29 | 70/70 | 0.492 → 0.500 | 1.00 | 1.00 |
| aldicarb | carbamate | A | 6–10 | 44 → 39 | 70/90 | 0.775 → 0.727 | 0.29 | 0.57 |
| aldicarb | carbamate | B | 9–10 | 41 → 40 | 70/90 | 0.764 → 0.745 | 0.37 | 0.58 |
| carbofuran | carbamate | A | 6–10 | 55 → 50 | 70/90 | 0.969 → 0.932 | 0.50 | 0.70 |
| carbofuran | carbamate | B | 9–10 | 46 → 44 | 70/90 | 0.857 → 0.820 | 0.16 | 0.35 |
| **carbaryl** | **carbamate** | **A** | **13–15** | **2 → 1** | 50/50 | 0.049 → 0.024 | **0.047** | 0.145 |
| **carbaryl** | **carbamate** | **B** | **13–15** | **6 → 2** | 50/50 | 0.146 → 0.049 | **$8 \times 10^{-5}$** | **0.001** |
| **carbaryl** | **carbamate** | **C** | **13–15** | **6 → 3** | 40/50 | 0.183 → 0.091 | **0.0012** | **0.009** |
| **carbaryl** | **carbamate** | **D** | **13–22** | **29 → 14** | 60/20 | 0.604 → 0.333 | **0.012** | **0.046** |
| carbaryl | carbamate | E | 17–22 | 32 → 30 | 60/20 | 0.667 → 0.714 | 0.59 | 0.79 |
| **malathion** | **organophosphate** | **A** | **10–22** | **40 → 20** | 70/70 | 0.702 → 0.392 | **$2 \times 10^{-5}$** | **0.001** |
| **malathion** | **organophosphate** | **B** | **13–22** | **37 → 24** | 70/70 | 0.661 → 0.471 | **0.0027** | **0.015** |
| malathion | organophosphate | C | 20–22 | 41 → 41 | 70/70 | 0.788 → 0.804 | 1.00 | 1.00 |
| **diazinon** | **organophosphate** | **A** | **6–22** | **29 → 8** | 70/60 | 0.487 → 0.190 | **$4 \times 10^{-4}$** | **0.003** |
| **diazinon** | **organophosphate** | **B** | **11–22** | **22 → 11** | 70/60 | 0.429 → 0.262 | **0.010** | **0.046** |

**Reading the table.** A treatment "fires" when $q_{BH} < 0.05$. The three uncouplers fire in none of eight testable treatments; 2,4-dichlorophenol C, on a three-day window, reaches $p$ =

0.035 and $q$ = 0.124. The narcotics fire in none of six. The three carbamates fire in three of nine, all carbaryl, the only carbamate whose design leaves a window longer than four days; malathion fires in two of three, on its windows of nine and twelve days, and diazinon fires in both treatments, on the longest windows in the set. Carbaryl treatment A ($q$ = 0.145) had two animals alive when its window opened.

**The comparison at comparable window length.** DNOC treatment B and carbaryl treatment D supply seven and nine days of clean water to nineteen and twenty-nine animals; the uncoupler loses none and the carbamate loses half. DNOC's elimination half-life in the same species and laboratory is 0.62 d,[25] so most of the compound had left the animals by the time the window opened.

**Table S7. Intensity of the pulse, for every treatment with a window.** "Acute kill" is the control-corrected fraction of exposed animals already dead when the window opens; "post-window kill" is the further fraction of survivors lost across it. Multiples of the two-day median lethal concentration use acute tests reported in the same data file and measured in the same laboratory; the aldicarb and carbofuran acute series do not bracket 50% survival at two days, so no multiple can be formed. Windows are the days between the two counts of Table S6.

| Compound | Trt | Window (d) | $C_{max}$ (nmol L⁻¹) | × LC50(2 d) | Acute kill | Post-window kill |
|---|---|---|---|---|---|---|
| 2,4,5-TCP | A | 4 | 385 | 0.17 | 0.74 | 0.01 |
| 2,4,5-TCP | B | 1 | 364 | 0.16 | 0.63 | 0.00 |
| DNOC | A | 9 | 2581 | 1.43 | 0.89 | − 0.08 |
| **DNOC** | **B** | **7** | **2612** | **1.45** | **0.69** | **− 0.04** |
| DNOC | C | 3 | 2611 | 1.45 | 0.78 | 0.06 |
| 2,4-DCP | A | 11 | 16094 | 1.04 | 0.15 | 0.05 |
| 2,4-DCP | B | 7 | 16337 | 1.06 | 0.27 | 0.05 |
| 2,4-DCP | C | 3 | 15946 | 1.03 | 0.21 | 0.09 |
| 1,2,3-TCB | A | 11 | 8788 | 1.99 | 0.42 | 0.15 |
| 1,2,3-TCB | B | 7 | 9815 | 2.22 | 0.70 | 0.10 |
| 1,2,3-TCB | C | 3 | 8547 | 1.93 | 0.51 | 0.00 |
| 2,4-DCA | A | 10 | 54855 | 1.05 | 0.74 | 0.13 |
| 2,4-DCA | B | 7 | 58768 | 1.13 | 0.83 | 0.17 |
| 2,4-DCA | C | 3 | 57514 | 1.10 | 0.51 | − 0.02 |
| aldicarb | A, B | 4, 1 | 3643, 3757 | — | 0.23, 0.24 | 0.06, 0.02 |
| carbofuran | A, B | 4, 1 | 49, 49 | — | 0.03, 0.14 | 0.04, 0.04 |
| carbaryl | A, B, C | 2 | 336, 339, 502 | — | 0.95, 0.85, 0.82 | 0.50, 0.67, 0.50 |
| **carbaryl** | **D** | **9** | **141** | — | **0.40** | **0.45** |
| carbaryl | E | 5 | 85 | — | 0.33 | − 0.07 |
| malathion | A | 12 | 4 | 1.16 | 0.30 | 0.44 |
| malathion | B | 9 | 4 | 1.16 | 0.34 | 0.29 |
| malathion | C | 2 | 4 | 1.16 | 0.21 | − 0.02 |

**The uncoupler pulse was the more damaging of the two.** By the time its window opened DNOC B had killed 69% of the exposed animals, against 40% for carbaryl D, at 1.45 times its two-day median lethal concentration; it then killed none, while carbaryl D killed nearly half of the survivors.

**Table S8. Post-exposure decline rate, every treatment with a window of at least one day.** $h_{\mathrm{post}} = -T_{\mathrm{window}}^{-1}\ln\,[S(t_{\mathrm{close}})/S(t_{\mathrm{open}})]$ on control-corrected survival. Sources: the thirteen-compound archive (ref 17) and the chlorpyrifos–pentachlorophenol series (ref 21). The rate is comparable between designs; the binary reading is not. The two diazinon rows are shown for comparison and are not pooled into the summaries below or into Figure 3: their windows are placed by the fitted damage maximum of §S6 rather than by the census rule of §S3, and rates measured under two different window rules do not belong in one median. The pooled set is therefore thirty-two treatments in eleven compounds. Intervals are profile-likelihood limits from the two binomials — treatment and concurrent control — over the window, which stay finite when an arm loses nobody; the delta method does not.

| Source | Compound | Mode of action | Trt | Window (d) | Alive | $h_{\mathrm{post}}$ ($d^{-1}$) | $t_{1/2}$ (d) | $p$ | 95% CI |
|---|---|---|---|---|---|---|---|---|---|
| 2015 | 2,4,5-TCP | uncoupler | A | 4.0 | 15 → 14 | 0.003 | 219 | 1.00 | −0.024 to 0.063 |
| 2015 | 2,4,5-TCP | uncoupler | B | 1.0 | 20 → 20 | − 0.000 | ∞ | 1.00 | −0.028 to 0.096 |
| 2015 | DNOC | uncoupler | A | 9.0 | 7 → 7 | − 0.009 | ∞ | 1.00 | −0.020 to 0.022 |
| **2015** | **DNOC** | **uncoupler** | **B** | **7.0** | **19 → 19** | **− 0.006** | ∞ | **1.00** | −0.017 to 0.010 |
| 2015 | DNOC | uncoupler | C | 3.0 | 13 → 12 | 0.020 | 34.5 | 0.37 | −0.017 to 0.112 |
| 2015 | 2,4-DCP | uncoupler | A | 11.0 | 52 → 47 | 0.005 | 150 | 0.47 | −0.005 to 0.016 |
| 2015 | 2,4-DCP | uncoupler | B | 7.0 | 43 → 40 | 0.008 | 87.9 | 0.31 | −0.004 to 0.025 |
| 2015 | 2,4-DCP | uncoupler | C | 3.0 | 46 → 42 | 0.030 | 22.9 | 0.035 | 0.008 to 0.070 |
| 2007a | pentachlorophenol | uncoupler | D-I | 2.0 | 20 → 20 | − 0.000 | ∞ | 1.00 | −0.020 to 0.048 |
| 2015 | 1,2,3-TCB | narcotic | A | 11.0 | 33 → 24 | 0.015 | 45.6 | 0.16 | −0.004 to 0.040 |
| 2015 | 1,2,3-TCB | narcotic | B | 7.0 | 16 → 13 | 0.016 | 43.9 | 0.37 | −0.013 to 0.064 |
| 2015 | 1,2,3-TCB | narcotic | C | 3.0 | 25 → 24 | 0.000 | 2546 | 1.00 | −0.033 to 0.049 |
| 2015 | 2,4-DCA | narcotic | A | 10.0 | 16 → 13 | 0.014 | 49.2 | 0.15 | −0.004 to 0.048 |
| 2015 | 2,4-DCA | narcotic | B | 7.0 | 10 → 8 | 0.027 | 25.6 | 0.095 | −0.002 to 0.094 |
| 2015 | 2,4-DCA | narcotic | C | 3.0 | 29 → 29 | −0.006 | ∞ | 1.00 | −0.025 to 0.017 |
| 2015 | aldicarb | carbamate | A | 4.0 | 44 → 39 | 0.016 | 43.1 | 0.29 | −0.011 to 0.053 |
| 2015 | aldicarb | carbamate | B | 1.0 | 41 → 40 | 0.025 | 28.1 | 0.37 | −0.013 to 0.109 |
| 2015 | carbofuran | carbamate | A | 4.0 | 55 → 50 | 0.010 | 71.2 | 0.50 | −0.015 to 0.039 |
| 2015 | carbofuran | carbamate | B | 1.0 | 46 → 44 | 0.045 | 15.6 | 0.16 | −0.001 to 0.137 |
| 2015 | carbaryl | carbamate | A | 2.0 | 2 → 1 | 0.347 | 2.0 | 0.047 | 0.019 to 1.634 |
| 2015 | carbaryl | carbamate | B | 2.0 | 6 → 2 | 0.549 | 1.3 | $10^{-4}$ | 0.165 to 1.368 |
| 2015 | carbaryl | carbamate | C | 2.0 | 6 → 3 | 0.347 | 2.0 | 0.0012 | 0.085 to 0.928 |
| **2015** | **carbaryl** | **carbamate** | **D** | **9.0** | **29 → 14** | **0.066** | **10.5** | **0.012** | 0.020 to 0.119 |
| 2015 | carbaryl | carbamate | E | 5.0 | 32 → 30 | − 0.014 | ∞ | 0.59 | −0.071 to 0.023 |
| 2007a | chlorpyrifos | OP | A-I | 2.0 | 5 → 4 | 0.099 | 7.0 | 0.21 | −0.020 to 0.483 |
| 2007a | chlorpyrifos | OP | A-II | 2.0 | 10 → 8 | 0.099 | 7.0 | 0.094 | −0.004 to 0.334 |
| 2007a | chlorpyrifos | OP | A-III | 2.0 | 23 → 19 | 0.083 | 8.3 | 0.052 | 0.005 to 0.211 |
| 2007a | chlorpyrifos | OP | B-I | 4.7 | 20 → 18 | 0.007 | 101 | 1.00 | −0.033 to 0.065 |
| 2007a | chlorpyrifos | OP | B-II | 1.0 | 8 → 6 | 0.288 | 2.4 | 0.028 | 0.048 to 0.895 |
| 2015 | malathion | OP | A | 12.0 | 40 → 20 | 0.048 | 14.3 | $2 \times 10^{-5}$ | 0.025 to 0.079 |
| 2015 | malathion | OP | B | 9.0 | 37 → 24 | 0.038 | 18.4 | 0.0027 | 0.013 to 0.070 |
| 2015 | malathion | OP | C | 2.0 | 41 → 41 | −0.010 | ∞ | 1.00 | −0.043 to 0.016 |
| **2015** | **diazinon** | **organophosphate** | **A** | **16.0** | **29 → 8** | **0.059** | **11.8** | **$4 \times 10^{-4}$** | 0.025 to 0.104 |
| **2015** | **diazinon** | **organophosphate** | **B** | **11.0** | **22 → 11** | **0.045** | **15.5** | **0.010** | 0.010 to 0.093 |

**Fifteen of seventeen anticholinesterase treatments have a positive rate, and one of nine uncoupler treatments exceeds 0.020 $d^{-1}$.** The anticholinesterase exceptions are carbaryl E, which loses two of thirty-two over five days against two of sixteen in its control, and malathion C, which loses none over two days. The uncoupler is 2,4-dichlorophenol C

(0.030 d⁻¹ over three days, $p$ = 0.035), the most slowly eliminated uncoupler in the archive (half-life 3.3 d against 0.4–0.7 d for the other three; Table 1). Medians are 0.003 d⁻¹ for nine uncoupler treatments in four compounds, 0.048 d⁻¹ for seventeen anticholinesterase treatments in five compounds and 0.015 d⁻¹ for six narcotic treatments in two compounds. The chlorpyrifos treatments are not among the twenty-eight tests of Table S6; their smallest unadjusted $p$ is 0.028 (B-II, a one-day window).

**The same compound gives opposite binary verdicts at different window lengths.** Diazinon treatment A falls from 0.487 to 0.190 over its full sixteen-day window ($p = 4 \times 10^{-4}$) and only to 0.400 over its first four days ($p$ = 0.12; $h_{\text{post}}$ = 0.049 d⁻¹, profile interval −0.005 to 0.123). Treatment B behaves the same way ($p$ = 0.010 over eleven days, $p$ = 0.15 over four). This is the design argument of §4.4 demonstrated within a single treatment, with the compound, the species, the animals and the exposure all held fixed.

**Table S9. Post-exposure decline summarised by compound.** $h_{\text{post}}$ as defined above; “fires” counts treatments with $q_{\text{BH}}$ < 0.05 under the binary reading. This is the compound-level summary of Table S8.

| Compound | Mode of action | Fate of the lesion | Treatments | Window range (d) | Median $h_{\text{post}}$ (d⁻¹) | Range | Fires |
|---|---|---|---|---|---|---|---|
| 2,4,5-trichlorophenol | uncoupler | none | 2 | 1–4 | 0.002 | 0.000, 0.003 | 0 |
| DNOC | uncoupler | none | 3 | 3–9 | − 0.006 | − 0.009 to 0.020 | 0 |
| pentachlorophenol | uncoupler | none | 1 | 2 | 0.000 | — | 0 |
| 2,4-dichlorophenol | uncoupler | none | 3 | 3–11 | 0.008 | 0.005 to 0.030 | 0 |
| **uncouplers** | | | **9** | **1–11** | **0.003** | − 0.009 to 0.030 | **0** |
| 1,2,3-trichlorobenzene | narcotic | reverses | 3 | 3–11 | 0.015 | 0.000 to 0.016 | 0 |
| 2,4-dichloroaniline | narcotic | reverses | 3 | 3–10 | 0.014 | − 0.006 to 0.027 | 0 |
| **narcotics** | | | **6** | **3–11** | **0.015** | − 0.006 to 0.027 | **0** |
| aldicarb | carbamate | reactivates | 2 | 1–4 | 0.020 | 0.016, 0.025 | 0 |
| carbofuran | carbamate | reactivates | 2 | 1–4 | 0.027 | 0.010, 0.045 | 0 |
| carbaryl | carbamate | reactivates | 5 | 2–9 | 0.347 | − 0.014 to 0.549 | 3 |
| chlorpyrifos | OP | does not reactivate | 5 | 1–5 | 0.099 | 0.007 to 0.288 | 0 |
| malathion | OP | does not reactivate | 3 | 2–12 | 0.038 | − 0.010 to 0.048 | 2 |
| diazinon* | OP | does not reactivate | 2 | 11–16 | 0.052 | 0.045, 0.059 | 2 |
| **anticholinesterases** | | | **17** | **1–12** | **0.048** | **− 0.014 to 0.549** | **5** |

* Windows placed by the fitted damage maximum of §S6, not by the census rule; the row is shown for comparison and excluded from the summary. Both its treatments fire under that rule (Table S6).

##### *S4.1 The same ordering in another species, on a different endpoint*

The rates of Table S9 order the compound classes by how long their lesion persists. An older experiment orders them the same way in a different species, on a different endpoint, and over hours rather than days. Kallander and colleagues gave fourth-instar *Chironomus riparius* two one-hour pulses at each compound’s one-hour EC20, separated by 0, 2, 6, 12 or 24 h in clean water, and scored the failure to respond to prodding.[22]

**Table S10. Effect of splitting a two-hour exposure, by compound class.** Mean adjusted percent of midges affected, from Table 1 of ref 22.

| Compound | Class | Continuous 2 h | 6 h apart | 24 h apart |
|---|---|---|---|---|
| aldicarb | carbamate | 81% | **12%** | 27% |
| carbaryl | carbamate | 68% | **12%** | 27% |
| propoxur | carbamate | 59% | **10%** | 12% |
| carbofuran | carbamate | 58% | **10%** | 10% |
| parathion | organophosphate | 93% | 48% | 58% |
| malathion | organophosphate | 100% | **99%** | 97% |

**Six hours of clean water is enough for a carbamate and not for malathion.** All four carbamates lose most of their effect; malathion loses none. Acetylcholinesterase measured in the same animals returns to control activity about six hours after a carbaryl pulse and is inhibited again by the next, with no cumulative loss across pulses twenty-four hours apart.[22]

**Parathion falls between the carbamate and malathion**, from 93% to 48%, and that is informative rather than awkward: the ordering is a gradient in adduct persistence rather than a dichotomy between classes. A diethyl phosphoryl adduct, as parathion leaves, ages more slowly than the dimethyl adduct of malathion and can still reactivate; a carbamylated enzyme reactivates in under an hour. The ordering here is therefore the same one Table S9 recovers from survival counts in *G. pulex*, obtained on a different species, a sublethal endpoint and a timescale three orders of magnitude shorter.

What it does not do is discriminate the hazard structures, for the reason §4.5 gives: it is a statement about how long damage lasts, not about how damage becomes death.

##### *S4.2 The compound as the experimental unit*

The thirty-two treatments of Table S8 are nested in eleven compounds, and the twelfth, diazinon, is measured on windows placed by the fitted trajectory of §S6. Because the claim under test is that $h_{\text{post}}$ is set by the compound, treatments within a compound are not independent replicates: the unit is the compound. Summarising each compound by the median of its treatments gives the twelve values of Table S11.

**Table S11. Compound medians of the post-exposure decline rate.** Diazinon is included here because the compound median is not affected by the window rule that excludes it from the pooled treatment set of Table S9.

| Class | Compound | Treatments | Median $h_{\text{post}}$ ($d^{-1}$) |
|---|---|---|---|
| uncoupler | DNOC | 3 | − 0.006 |
| uncoupler | pentachlorophenol | 1 | − 0.000 |
| uncoupler | 2,4,5-trichlorophenol | 2 | 0.002 |
| uncoupler | 2,4-dichlorophenol | 3 | 0.008 |
| narcotic | 2,4-dichloroaniline | 3 | 0.014 |
| narcotic | 1,2,3-trichlorobenzene | 3 | 0.015 |
| anticholinesterase | aldicarb | 2 | 0.020 |
| anticholinesterase | carbofuran | 2 | 0.027 |
| anticholinesterase | malathion | 3 | 0.038 |
| anticholinesterase | diazinon | 2 | 0.052 |
| anticholinesterase | chlorpyrifos | 5 | 0.099 |
| anticholinesterase | carbaryl | 5 | 0.347 |

**The separation is complete**: every anticholinesterase compound lies above every uncoupler, with no overlap. With six compounds against four the two-sided Mann–Whitney test returns $p = 0.0095$, which is the smallest value it can return at those sample sizes — complete separation cannot yield a smaller one. The two narcotics lie between the classes and are not part of this comparison.

Replacing the median by the mean within each compound, or discarding treatments with fewer than ten animals alive when the window opens, leaves every compound on the same side. Replacing it by the largest treatment rate does not: 2,4-dichlorophenol C (0.030 $d^{-1}$) exceeds the largest aldicarb rate (0.025 $d^{-1}$). Reproduced by `s04_compound_test.py` (§S16).

**What the intervals show.** Nine of the thirty-two intervals exclude zero: eight are anticholinesterase treatments and one is 2,4-dichlorophenol C (0.008 to 0.070 $d^{-1}$). The other uncoupler intervals are narrow — DNOC's seven- and nine-day windows admit rates no larger than 0.010 and 0.022 $d^{-1}$ — so their silence is a measured bound rather than a failure to look. Among the anticholinesterases the intervals that include zero are the short windows: aldicarb and carbofuran on one to four days admit rates up to 0.14 $d^{-1}$. Reproduced by `s04_rate_intervals.py` (§S16).

##### *S4.3 Depletion of the frail*

If individuals differ in tolerance, a treatment that has already killed many animals opens its window on a selected, hardier remainder, which would decline more slowly afterwards for reasons that have nothing to do with the compound. The prediction is a negative association between mortality at the opening of the window and the rate measured across it. Regressing the rate on that mortality and on mode of action over the thirty-two treatments of Table S8 gives the opposite sign (Table S12).

**Table S12. The decline rate against prior mortality and mode of action.** $R^2 = 0.50$, $n = 32$. Mortality at the opening of the window is control-corrected, as in Table S8. Mode of action enters as two indicators against the uncouplers, anticholinesterase and narcotic; entering carbamate and organophosphate separately changes the mortality slope to $+0.291$ and $R^2$ to 0.58 without changing its sign.

| Term | Coefficient ($d^{-1}$) | 95% CI | *p* |
|---|---|---|---|
| intercept | − 0.145 | − 0.244 to − 0.046 | 0.006 |
| mortality at window opening | **+ 0.275** | + 0.139 to + 0.411 | 0.0003 |
| anticholinesterase | **+ 0.133** | + 0.051 to + 0.214 | 0.0025 |
| narcotic | − 0.015 | − 0.119 to + 0.090 | 0.78 |

**Treatments are nested in compounds, and the slope survives that.** The row-level fit treats thirty-two treatments as independent, which they are not: they come from eleven compounds. Clustering the standard error on the compound leaves the slope significant — on the three-level specification above, SE = 0.123, $t = 2.37$, $p = 0.040$ with $G = 11$ clusters. Taking the compound as the unit instead of the treatment weakens it to a slope of $+0.142$ with Spearman $\rho = +0.01$ ($p = 0.98$). Neither reading is negative, and depletion of the frail predicts a negative slope, so the confounder is excluded at the treatment level and not supported at the compound level.

Treatments that have lost more animals decline faster, not more slowly, and the mode-of-action term survives the adjustment. The uncouplers are the harsher exposures (median mortality at window opening 63% against 33% for the carbamates) and lose nobody afterwards (Figure 3c).

A subset matched on prior mortality is not available: eleven treatments fall between 25% and 60%, but only two of them are uncouplers. The carbamates reach the highest rates at peak concentrations an order of magnitude below the uncouplers'. Reproduced by `s04_frailty_regression.py` (§S16).

---

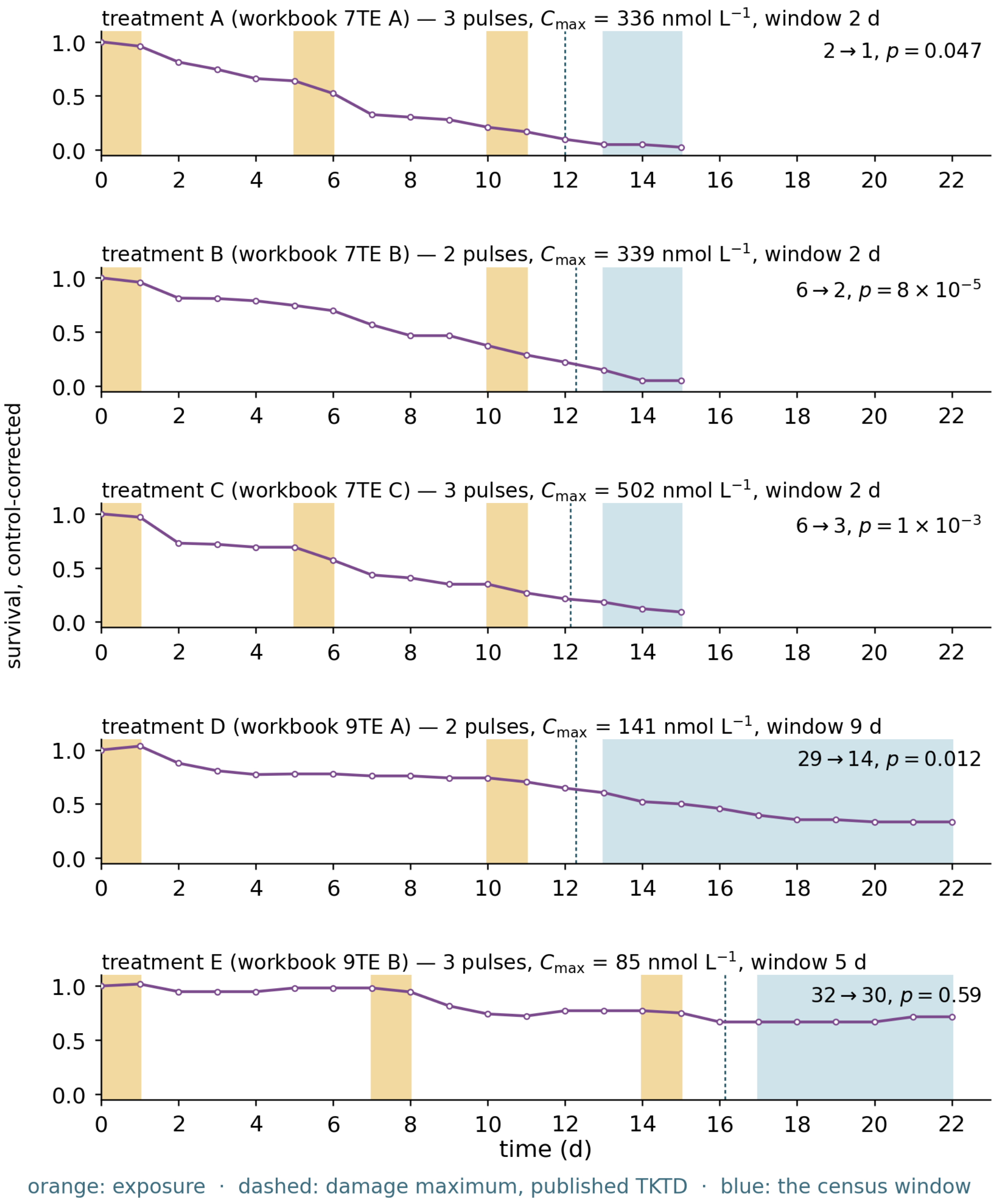


**Figure S2. The carbamate of the matched pair, every treatment.** Carbaryl in *Gammarus pulex*, the five treatments of the 2015 archive: A–C at 50 or 40 animals in a fifteen-day test, D and E at 60 animals in a twenty-two-day test, each with its concurrent control. Bands and dashed line as in Figure S3 (§S6); the damage maximum is computed under the published two-compartment model for this compound and species ($k_{out}$ = 0.27, $k_r$ = 0.97 $d^{-1}$),[26] driven by the measured profile, and falls 1.0–1.3 d after the pulse ends, before the census window of §S3 opens in every treatment. In treatment E the running maximum is set by the second pulse on day 9.2. Survival is control-corrected; annotations give the counts at the window's opening and close and the Fisher $p$ of Table S6. Axes share a two-day tick spacing with Figure S3.

#### S5. The Rate Outside the Archive

Nineteen pulsed-exposure studies were assembled and each was scored against the requirements of §4.7: a clean-water window after the last pulse, the number alive at both of its ends, a measured exposure profile so that the window can be placed, and a concurrent control. Five studies supply all four requirements; the four in *G. pulex* are used here and the neonicotinoid interval series in §S7. Seven are ruled out, and seven could not be scored because the source was not available to this analysis.

**Table S13. Nineteen pulsed-exposure studies against the requirements of §4.7.** The full scoring, with the reason recorded for each study, is in `pulsed_exposure_inventory_C.csv` (§S16).

| Study | Species | Window after the last pulse | Counts at both ends | Exposure | Rate computable |
|---|---|---|---|---|---|
| Ashauer *et al.* 2015[17] | *G. pulex* | 1–9 d | yes | measured | **yes** |
| Ashauer *et al.* 2007a[21] | *G. pulex* | 1–4.7 d | yes | measured | **yes** |
| Ashauer *et al.* 2007b[26] | *G. pulex* | varied | yes | measured | **yes** |
| Ashauer *et al.* 2010[20] | *G. pulex* | 11–16 d | yes | measured | **yes** |
| Zhao & Newman 2006[23] | *H. azteca* | to no further mortality | **figures only** | measured | no |
| Naddy *et al.* 2000[32] | *D. magna* | 7, 14, 18 d | **21-d totals only** | measured | no |
| Andersen *et al.* 2006[24] | *D. magna* | about 20 d | 21-d survival only | **nominal** | no |
| Focks *et al.* 2018[5] | five species | 2–17 d | yes | measured | **yes** |
| Kallander *et al.* 1997[22] | *C. riparius* | **none** | — | measured | no |
| three modelling studies | various | — | none | — | no |
| seven further studies | various | not seen | not seen | not seen | unknown |

**Two of the studies ruled out were designed to carry the test but do not report the counts it needs.** Zhao and Newman followed latent mortality in clean water until it stopped, and Naddy and colleagues left seven to eighteen days between the last pulse and the end of a twenty-one-day test; both measured exposure and ran concurrent controls, but one reports cumulative-mortality curves and the other twenty-one-day totals.

What can still be read from these studies is the ordering, and it agrees. Zhao and Newman estimated the recovery interval needed for an amphipod to return to its original resistance at 83 h for dissolved copper against 18 h for pentachlorophenol, a fivefold difference between two compounds in one species — and pentachlorophenol is the uncoupler whose rate in *G. pulex* is $-0.000\ \mathrm{d}^{-1}$ (Table S9).[23] Andersen and colleagues report that *Daphnia magna* regain mobility within a day of transfer to clean water after a dimethoate pulse but are killed by a second pulse forty-eight hours later, so the lesion outlasts the visible impairment.[24] Neither observation is a rate, and neither enters Table S9 or any summary in this paper; they are recorded because, with the midge data of §S4.1, they place the short-lived uncoupler lesion below longer-lived ones in three further species.

---

#### S6. The Diazinon Experiment: Window Placement, Test Results, and Interval Ordering

**The experiment.** Diazinon in *Gammarus pulex* has one published pulsed experiment:[20] treatments A, B and C of two one-day pulses of about 100 nmol/L separated by 2, 7 or 15 days, seven beakers of ten animals each, six concurrent control beakers, counted daily to day 22. The same counts are the diazinon sheet of the 2015 data supplement.[17] The original analysis fitted the three treatments together under the Threshold Damage Model, a cumulative hazard.[20]

**Where the window opens.** Here the window opens where the fitted damage trajectory stops rising: there the running maximum freezes, and beyond it a peak-based hazard predicts survival flat apart from background mortality. The trajectory is the companion paper's three-compartment model — internal parent, activated metabolite and damage — with the toxicokinetics measured in the same study and the companion's peak-based calibration fit.[4] Its recovery rate sits at the lower bound of the fit, so damage reaches 99.9% of its maximum 1.6 d after the last pulse ends and changes by less than 0.1% thereafter; the window opens at the first count after that point.

**Why a second window rule, and whether it changes the verdict.** The archive-wide census of §S3 cannot use a fitted trajectory, because most sheets have not been fitted, so it opens the window two days after the last pulse instead; on the diazinon treatments the two rules agree to within a day, the resolution at which the counts are recorded. The placement depends on the damage model, but the verdict does not. Under the reduced GUTS form, a single scaled-damage compartment that stops rising within hours of the pulse, the windows of treatments A and B open on days 4 and 9 rather than 6 and 11, and the test returns smaller $p$-values, $1 \times 10^{-4}$ and $3 \times 10^{-6}$. The three-compartment windows are the ones reported because they are the conservative choice: opening later, they leave out the first days of clean water and give the test less to resolve. Opened at the strict maximum of the trajectory instead, days 8 and 13, the test returns $p = 9 \times 10^{-4}$ and 0.004.

**Background mortality.** The concurrent controls give $h_b = 0.0232\ \mathrm{d}^{-1}$ (60 animals reduced to 36 over 22 d); that value is shown for comparison only, since the tests use the controls directly.

**Table S14. Window definitions and results.** "Damage at 99.9% of maximum" is the day the fitted trajectory reaches 99.9% of its maximum; the window opens at the next count. The test is Fisher's exact test against the concurrent controls over the same window.

| Trt | End of the preceding pulse | Damage at 99.9% of maximum | Window (d) | Alive, open → close | Controls, open → close | $p$ |
|---|---|---|---|---|---|---|
| A | day 4.02 | day 5.66 | 6–22 | 29 → 8 | 51 → 36 | $4 \times 10^{-4}$ |
| B | day 9.01 | day 10.63 | 11–22 | 22 → 11 | 44 → 36 | 0.010 |
| C* | day 1.03 | day 2.88 | 3–16 | 55 → 38 | 51 → 38 | 0.67 |

* The window between the two pulses, the direct analogue of the imidacloprid window of §4.2. For this row the pulse named is the first; damage reaches 99.9% of its maximum 1.9 d after it, on day 2.88, and the window opens at the next count, day 3. Being a between-pulse window rather than a post-exposure one, it is not part of the clean-water test set of Table S6.

**Table S15. Interval ordering in the diazinon experiment.** Same total exposure, the second pulse following a clean-water interval of 2, 7 or 15 days.

| Interval | Survivors at 22 d (of 70) | Control-corrected survival at 22 d |
|---|---|---|
| 2 d | 8/70 | 0.190 |
| 7 d | 11/70 | 0.262 |
| 15 d | 19/70 | 0.452 |

**A two-pulse interval design gives ordinary recovery for diazinon.** Survival at 22 d rises with the interval (0.190, 0.262, 0.452; Fisher's exact test, 2 d against 15 d, $p = 0.031$), the direction both hazard structures predict; the neonicotinoid series of §4.6 runs the other way.

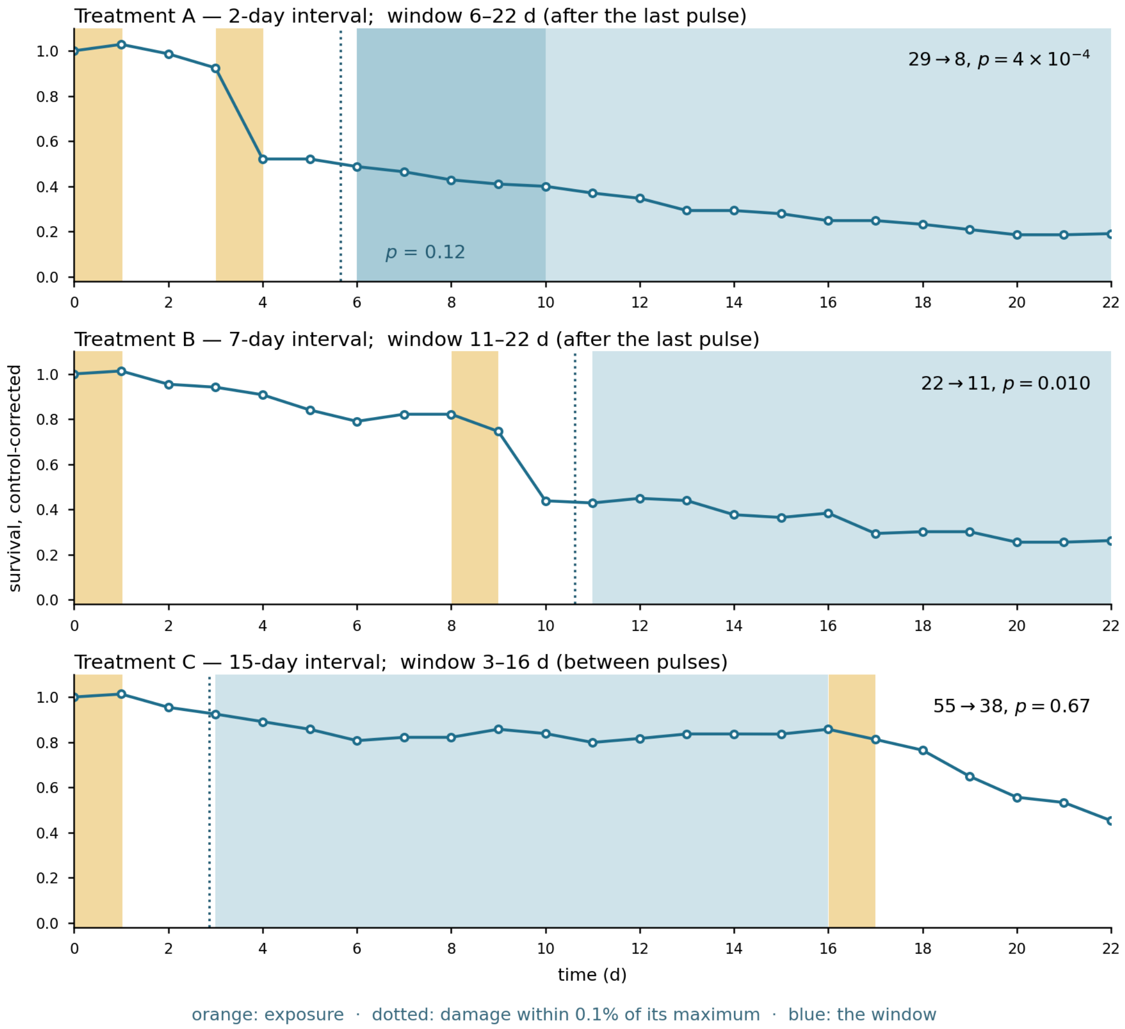


**Figure S3. One compound, one experiment, every treatment.** Diazinon in *Gammarus pulex*, the three treatments of ref 20 at intervals of two, seven and fifteen days; survival is control-corrected. Orange bands are measured exposure, the dotted line the day the fitted damage reaches 99.9% of its maximum, where the running maximum freezes, and the blue band the window it opens. In treatment A the darker blue is the first four days of the window, over which the same test gives $p = 0.12$ against $4 \times 10^{-4}$ for the full window. In treatment C the window lies between the pulses. The clean-water test fires on both treatments with a post-exposure window (A, B). Reproduced by `s06_diazinon_one_experiment.py` (§S16).

#### S7. The Five-Species Imidacloprid Interval Series: Survivor Counts, Immobilisation, and the Window after the Second Pulse

**The design.** Focks and colleagues exposed five macroinvertebrates — *Asellus aquaticus*, *Caenis horaria*, *Chaoborus obscuripes*, *Cloeon dipterum* and *Plea minutissima* — to one total imidacloprid exposure delivered three ways: a single 48-hour pulse, or two 24-hour pulses starting 7 or 14 days apart, each arm five replicates of fifteen animals beside a concurrent control, with concentrations measured in every replicate and counts to day 21–28.[5,16] Days are counted from the start of exposure, so the first census is day 1. In *P. minutissima* all three arms received a third pulse on day 21,[5] so its comparison of equal exposures ends at the day-21 census. This is the design of §4.6. This section gives the survivor counts behind the reversal (Table S16), what the immobilisation record shows between the pulses (Table

S17), the decline after the second pulse (Table S18), and the developmental explanation that §4.6 sets aside.

**Survivor counts: one species reverses.** *Chaoborus obscuripes* and *Cloeon dipterum* lose every exposed animal in all three arms and so cannot show either ordering; the other three are in Table S16.

**Table S16. Survivor counts at each species' last census.** Day 22 for *A. aquaticus*, 21 for *C. horaria* and 21 for *P. minutissima*, the last census before its third pulse. Read from the raw records; these are the counts behind Figure S4b. The *P. minutissima* single-pulse arm was set up with 76 animals rather than 75, as recorded in the source. The last column gives a Cochran–Armitage test for trend across the three splittings and, in parentheses, Fisher's exact test between the single pulse and the fourteen-day treatment.

| Species | Control | 48-h single pulse | 24-h pulses, 7 d apart | 24-h pulses, 14 d apart | Trend *p* (Fisher *p*) |
|---|---|---|---|---|---|
| *A. aquaticus* | 65/75 | **51** | **40** | **29** | $3 \times 10^{-4}$ ($5 \times 10^{-4}$), falling |
| *P. minutissima* | 75/75 | 61 (of 76) | 57 | 53 | 0.17 (0.19), falling, not significant |
| *C. horaria* | 67/75 | **16** | **29** | **31** | 0.010 (0.013), rising |

As the pulses are moved further apart, *A. aquaticus* declines and *C. horaria* recovers, both significantly; *P. minutissima* declines in the same direction as *A. aquaticus* without significance. The tests treat each arm as a single binomial: the source records fifteen animals in each of five replicates, and replicate-level overdispersion cannot be estimated from the published tables; allowing for it could only make these *p*-values larger, so they are lower bounds. Reproduced by `s07_trend_tests.py` (§S16).

**What the first pulse leaves: the immobilisation record.** The data file records dead, alive and immobile animals separately, so immobilisation can be read alongside survival without any further experiment. Its usefulness varies by species: *A. aquaticus* controls are scored immobile at fractions between 0.17 and 1.00 throughout, because isopods rest motionless, so the column carries no information there; *P. minutissima* and *C. horaria* controls show no immobilisation. Table S17 gives the record for *P. minutissima* in the fourteen-day treatment, where survival is complete between the pulses.

**Table S17. *Plea minutissima*, fourteen-day interval treatment.** Selected census days up to day 21, before the third pulse, survivors and immobilised animals against the concurrent control (75 per arm). The full record is in the source data file.

| Day | 1 | 2 | 3–14 | 15 | 16 | 17 | 21 |
|---|---|---|---|---|---|---|---|
| control, alive | 75 | 75 | 75 | 75 | 75 | 75 | 75 |
| control, immobile | 0 | 0 | 0 | 0 | 0 | 0 | 0 |
| treatment, alive | 75 | 75 | **75** | 69 | 64 | 61 | **53** |
| treatment, immobile | 55 | 6 | **0** | 69 | 64 | 61 | 0 |

The first pulse immobilises fifty-five animals on day one and six on day two; from day three to day fourteen immobilisation is zero and survival is complete. The second pulse then immobilises every survivor and mortality follows. For the same total exposure delivered as a single 48-hour pulse, survival at day twenty-one is sixty-one rather than fifty-three.

**The window after the second pulse.** In the fourteen-day treatment the second 24-hour pulse ends on day 15 and observation continues to day 21 in all five species, so the treatment leaves six days of clean water — longer than the 4.6 d that §4.7 requires for imidacloprid. The window is counted from day 16, the first census after the pulse, to day 21, the last census common to all five species. Three species have enough survivors to test, and all three decline against the control (Table S18).

**Table S18. Post-exposure decline in the fourteen-day interval treatment.** Window from day 16 to day 21 (five days). Survival is control-corrected; $h_{post}$ and its profile-likelihood interval are computed as in Table S8, and $p$ is Fisher's exact test against the concurrent control, as in Table S6. Reproduced by `s07_second_pulse_window.py` (§S16).

| Species | Alive, day 16 → 21 | Control, day 16 → 21 | Control-corrected survival | $h_{post}$ ($d^{-1}$) | 95% interval | $p$ |
|---|---|---|---|---|---|---|
| *Asellus aquaticus* | 43 → 29 | 68 → 66 | 0.69 | 0.073 | 0.037 to 0.123 | $2 \times 10^{-5}$ |
| *Caenis horaria* | 42 → 31 | 70 → 67 | 0.77 | 0.052 | 0.020 to 0.096 | 0.002 |
| *Plea minutissima* | 64 → 53 | 75 → 75 | 0.83 | 0.038 | 0.020 to 0.065 | $1 \times 10^{-4}$ |
| *Chaoborus obscuripes* | 4 → 0 | — | — | not testable | — | — |
| *Cloeon dipterum* | 0 | — | — | not testable | — | — |

**These three rates are the bracketed entry of Table 1.** Their median, 0.052 $d^{-1}$, falls between the carbamate and organophosphate medians. Each pulse was dosed and removed on a set schedule, so here the window opens at the first census after removal.

**The census rule applied to all fifteen treatments.** Opening the window two days after measured exposure ends, as §S3 does for the archive, eleven of the fifteen treatments meet the four requirements of §4.7: every arm of *A. aquaticus* and *C. dipterum*, the single-pulse and seven-day arms of *C. horaria* and *P. minutissima*, and the single-pulse arm of *C. obscuripes*. Survival falls against the control in eight of the eleven — every arm of *A. aquaticus* ($p \leq 0.009$) and *C. dipterum*, and the single-pulse arms of *C. horaria* ($p = 5 \times 10^{-6}$) and *C. obscuripes* — and not in the seven-day arm of *C. horaria* ($p = 0.17$) or either *P. minutissima* arm ($p = 0.45$ and 1.00). The measured profiles of *C. dipterum* record no second pulse in its interval arms, which are therefore scored as single pulses. Opened instead three upstream half-lives of §4.3 (4.6 d) after exposure ends, seven treatments qualify and four still decline: the single-pulse and seven-day arms of *A. aquaticus* ($p = 0.022$ and 0.002) and the single-pulse arms of *C. horaria* and *C. obscuripes*. Reproduced by `s07_census_rule_window.py` (§S16).

**An alternative explanation set aside.**

*A developmental explanation does not fit.* Receptor density in aquatic insects changes three- to sevenfold between larval and adult stages,[28] so a developmental account predicts the reversal in species that develop within the test. *A. aquaticus* and *P. minutissima*, the first of which reverses, neither emerge nor reproduce over the twenty-eight days, whereas *C. horaria*, which emerges, recovers in the ordinary way.

**Exposure relative to each species' own sensitivity.** The reversing species, *A. aquaticus*, received 5.8 times its own four-day median effect concentration, *P. minutissima*, which falls in the same direction without significance, 1.3 times, and the recovering *C. horaria* ten times its own.[9] With the sensitivities the same laboratory measured two years earlier, in May and June 2012,[34] the multiples become 3.8, 7.0 and 34: the numbers move with the season, but the recovering species again received the most. If this is more than coincidence, the reversal is a low-exposure phenomenon and should disappear as the multiple rises — a prediction these three species cannot test.

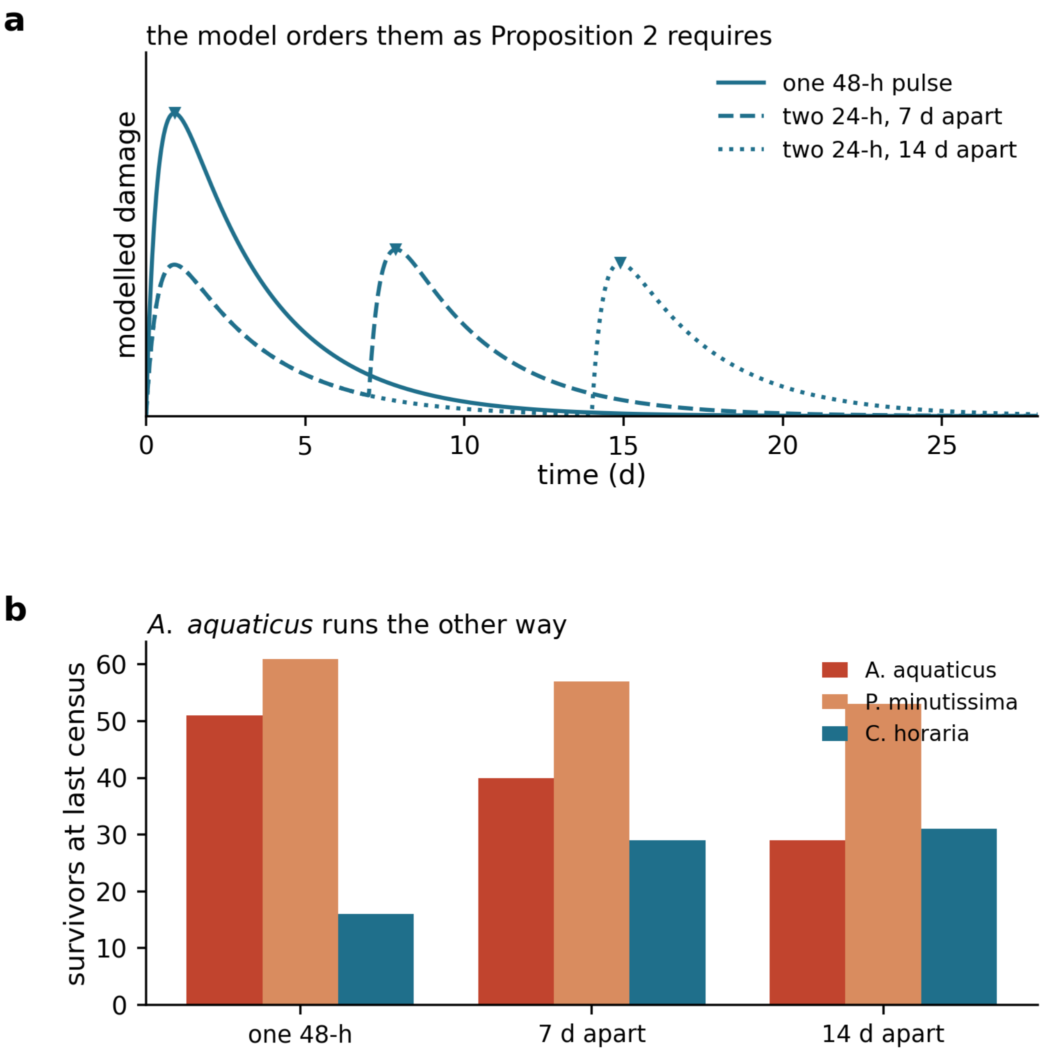


**Figure S4. Splitting a fixed exposure, and what it says about recovery.** (a) Modelled damage on a two-stage cascade for one exposure delivered as a single 48-hour pulse or as two 24-hour pulses seven or fourteen days apart; markers show each maximum. (b) Observed survivors at each species' last census (Table S16) for the three species with survivors. The model orders the three designs as Proposition 2 requires; *A. aquaticus* runs the other way; *P. minutissima*, at day 21, falls in the same direction without significance. The corresponding figure for the bound of Proposition 3 is Figure S5.

#### S8. How Much Three Neonicotinoids Recover Between Pulses, Read from Survival Counts Alone

**The design.** In a second experiment from the same study, Focks and colleagues exposed *Cloeon dipterum* to imidacloprid, thiacloprid and thiamethoxam in two 24-hour pulses, the second starting on day 7 or on day 14, each pulse set to the compound's 48-hour $LC_{30}$; three replicates of ten animals per arm, a common control of ninety, counts to day 28.[5,16] The two arms of a compound differ only in the interval, so comparing their survival asks how much the animals recovered in the seven extra days between a second pulse on day 7 and one on day 14.

**Why a ratio of survival counts bounds the recovery.** Under a peak-based hazard, survival is $S = \exp(-k[M-\theta]^{+})$ up to the background, so $-\ln S$ is proportional to the damage held above the threshold. The ratio of $-\ln S$ between the 14-day and the 7-day arm is therefore the ratio of that excess — the excess-damage ratio — and the killing rate $k$

cancels from it. The threshold does not cancel, but it can only push the ratio down: with $M$ the smaller peak, from the 14-day arm, $[M-\theta]^+/[M'-\theta]^+ \leq M/M'$ whenever $M' > M > \theta > 0$. The excess-damage ratio is therefore a lower bound on the ratio of peak damage, whatever the tolerance distribution and whatever repair law is at work. That makes it informative in one direction only. A ratio near one forces the peak ratio near one: the animals recovered almost nothing in the seven extra days. A ratio well below one shows that the second peak was lower, so some recovery occurred, but not how much, because a threshold close to the peaks turns a small fall in peak damage into a large fall in excess damage.

**Table S19. Recovery inferred from survival counts alone.** $S$ is survival at day 28 in the arm whose second pulse started on day 7 or day 14; the excess-damage ratio is $-\ln S(14\text{ d})/-\ln S(7\text{ d})$, a lower bound on the ratio of peak damage.

| Compound | $S$(7 d) | $S$(14 d) | $-\ln S$ (7 d) | $-\ln S$ (14 d) | Excess-damage ratio | Reading |
|---|---|---|---|---|---|---|
| imidacloprid | 8/30 | 9/30 | 1.32 | 1.20 | 0.91 | almost no recovery in seven further days |
| thiacloprid | 0/30 | 10/30 | ≥ 4.09* | 1.10 | ≤ 0.27* | recovery detected; its size not bounded |
| thiamethoxam | 0/30 | 0/30 | ≥ 4.09* | ≥ 4.09* | undetermined* | both arms lose every animal |

* A zero count is censored at half an animal, so $-\ln S$ is a lower bound there; for thiacloprid the ratio is then an upper bound, and for thiamethoxam, with both arms at zero, no ratio can be formed. The ratios are computed on raw survival; the concurrent control loses 17 of 90 animals over the 28 days, and correcting for it gives 0.89 and ≤ 0.23 in place of 0.91 and ≤ 0.27, leaving each reading unchanged. The bound on thiacloprid depends on the censoring rule only in the third decimal: one third of an animal gives ≤ 0.21, the Jeffreys value $0.5n/(n+1)$ gives ≤ 0.23 and a whole animal ≤ 0.28, so the reading — recovery detected, unlike imidacloprid — does not turn on the convention.

**Converting the imidacloprid ratio to a half-life requires a damage model.** Under the two-stage model of §4.3, a peak ratio of 0.91 corresponds to an upstream half-life of 5.3 d (Figure S5). Because 0.91 is a lower bound on the peak ratio, 5.3 d is a lower bound on the half-life, reached when the threshold is zero; the threshold itself is not identified by two treatments. The value is within an order of magnitude of the 1.5 d fitted in a different species, *D. magna* (§4.3).

**The compound that recovers is the one with the cyano group.** Thiacloprid is the only one of the three whose counts show recovery between the two intervals; it carries a cyano-substituted heterocycle conferring higher detoxification potential, whereas imidacloprid and thiamethoxam carry a nitro group.

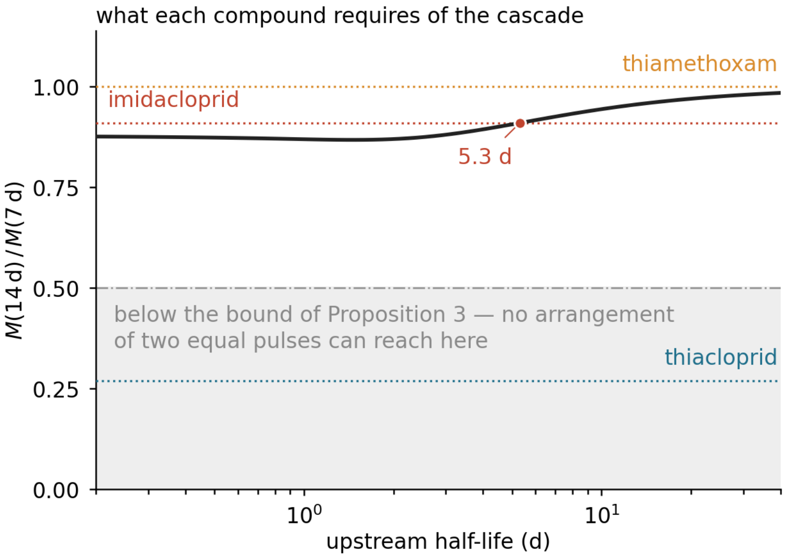


**Figure S5. Recovery timescale per compound, and the bound of Proposition 3.** The ratio of peak damage between the 14-day and 7-day designs as a function of the upstream half-life on a two-stage cascade. Horizontal lines mark the excess-damage ratios of Table S19 — lower bounds on the peak ratio — and a point marks the half-life at which the curve reaches each; the thiamethoxam line at 1.00 records only that both arms lose every animal, not a measured ratio. The shaded region lies below the bound of Proposition 3, which no pair of equal pulses can reach under convex repair; see §S8 and §S10.2.

### Part IV. Model Fits and Controls

#### S9. Materials and Methods for the Model Fits

Three results in the main text rest on fitted models: the *Daphnia magna* comparison of §4.3, the uncoupler comparison of §4.5, and the window placement for diazinon (§S6). This section gives the model configurations, the estimation procedure and the comparison metrics as applied to the *D. magna* series of Li *et al.*;[6] the DNOC fit uses the same configurations with toxicokinetics from the measured half-life (§S12), and the diazinon windows use the companion paper's fitted three-compartment model rather than a fit made here (§S6). The fitted parameters are in Tables S29–S31 and S34, and the identifiability checks in §S15.2.

##### *S9.1 Model configurations*

**Toxicokinetics.** One compartment, with uptake $k_u$ and elimination $k_e$,

$$\frac{\mathrm{d}C_b}{\mathrm{d}t} = k_u C_w - k_e C_b,$$

both held fixed at the published values.

**Damage.** The damage formation rate is $k_a$. Damage is either single-stage,

$$\frac{\mathrm{d}D}{\mathrm{d}t} = k_a C_b - R(D),$$

or two-stage, with an upstream state $D_1$ that persists after the parent has cleared and is lost at the upstream clearance rate $k_{\mathrm{off}}$,

$$\frac{\mathrm{d}D_1}{\mathrm{d}t} = k_a C_b - k_{\mathrm{off}} D_1, \qquad \frac{\mathrm{d}D_2}{\mathrm{d}t} = k_{12} D_1 - R(D_2)$$

with repair either proportional, $R(D) = k_r D$, or Michaelis–Menten, $R(D) = R_{\max} D/(K{+}D)$. In the notation of §2 the damage input is $u = k_a C_b$ on the single-stage model and $u = k_{12} D_1$ on the two-stage one, $k_{12}$ being the transfer rate from $D_1$ to $D_2$. Only the second input can remain positive once the parent has cleared, which is the difference §4.3 turns on.

**Hazard.** Cumulative,

$$H_A(t;z) = \int_0^t b_w \, [D(\tau) - z]^+ \, \mathrm{d}\tau + h_b t$$

or peak-based,

$$H_B(t;\theta) = k[M(t) - \theta]^+ + h_b t, \qquad M(t) = \max_{\tau \le t} D(\tau).$$

The killing rate constants $b_w$ and $k$ have different dimensions and are not comparable directly; only the dimensionless products $b_w z T$ and $k\theta$ can be compared between structures, and no such comparison is made here.

**Background.** The term $h_b t$ above is what the source implementation supplies as a multiplicative control series, $S = e^{-H} S_b$ with $S_b = e^{-h_b t}$.[6] That series is not reproduced in the source Supporting Information, so an exponential fixed by the reported 76% control survival at 22 d is used. Because the likelihood is conditional binomial, a constant factor on $S$ cancels in the ratios $S_i/S_{i-1}$ and only the shape of the control curve matters; no intermediate control counts are reported, so an exponential is the weakest assumption available.

**Which combinations are fitted.** Damage stage and repair law are not crossed freely. Three pairings — single-stage with proportional repair, two-stage with proportional repair, and two-stage with Michaelis–Menten repair — are fitted under each hazard structure, giving the six variants of Table S29; a seventh gives the cumulative structure a threshold read on $D_1$ (§S11). A single-stage Michaelis–Menten model is not fitted: with one compartment the removal term acts on the same state the threshold reads, so its shape parameter would be estimated from the same information as the threshold.

**Table S20. Symbols in this paper and in the companion's full cascade.** The persisting state $D_1$ is a damage precursor, not the companion's activated metabolite $A$.

| **Quantity** | **This paper (reduced)** | **Companion (full cascade)**[4] |
|---|---|---|
| external concentration | $C_w(t)$ | $C_{\mathrm{ext}}(t)$ |
| internal parent | $C_b$ | $C_i$ ($\equiv C_b$) |
| toxicokinetic rates | $k_u$ (uptake), $k_e$ (elimination) | $k_{\mathrm{in}}, k_{\mathrm{out}}$ |
| damage input | $k_a C_b$, or $k_{12} D_1$ with the persisting state | $k_3 A$ |
| repair | $k_r$, or $R_{\max}, K$ | $k_r$ |
| hazard and tolerance | $b_w, z$ (cumulative); $k, \theta$ (peak-based); $h_b$ | the same |

##### *S9.2 Parameters held fixed*

**Table S21. Constants held fixed in every fit.** Rate constants are per hour throughout, following the units of the source implementation, whose model code is reproduced in its Supporting Information;[6] half-lives are converted to days where reported.

| Symbol | Value | Basis |
|---|---|---|
| $k_u$, $k_e$ | published | measured uptake and elimination of the source study[6] |
| $k_{12}$ | 1 $h^{-1}$ | degenerate with $k_a$ (below) |
| $h_b$ | fixed by 76% control survival at 22 d | control series not reproduced in the source |

$k_{12}$ is fixed because it is degenerate with $k_a$: scaling $k_{12}$ by $\alpha$ and $k_a$ by $1/\alpha$ scales $D_1$ by $1/\alpha$ and leaves the input $k_{12}D_1$ to $D_2$, and so $D_2$ itself, unchanged; only the product is identified. Because repair acts on $D_2$, this holds for either repair law, and fixing $k_{12}$ loses no generality.

##### *S9.3 Estimation*

Each variant is fitted with one parameter set to all treatments of a series at once, so that the dependence on interval is predicted rather than fitted treatment by treatment. The objective is the conditional binomial negative log-likelihood,

$$\mathrm{NLL} = -\sum_i \left[ n_i \ln \frac{S_i}{S_{i-1}} + (n_{i-1} - n_i) \ln\left(1 - \frac{S_i}{S_{i-1}}\right)\right]$$

with $n_i$ the count alive at the $i$th census. Optima are found by bounded Nelder–Mead from randomised log-uniform starts, with warm starts from every nested special case — the two-stage Michaelis–Menten model reduces to the single-stage proportional model as $k_{\mathrm{off}} \to \infty$ with $k_a/k_{\mathrm{off}}$ fixed and $K \to \infty$ with $R_{\max}/K$ fixed — so that a larger model cannot report a worse optimum than a model it contains through non-convergence alone.

**Why the warm starts matter.** Started cold, the five-parameter cumulative model converges to a likelihood 17 units worse than its own three-parameter special case (Table S29), and the peak-based fit stalls eight units above its own optimum.

Damage is integrated by exponential-Euler on each linear stage, with the Michaelis–Menten rate held constant over the step. Convergence was checked for every reported comparison by halving $\Delta t$; for the decisive comparison of Table S29 the likelihood difference changed by less than 0.3 units between $\Delta t$ = 0.02 d (0.48 h) and $\Delta t$ = 0.01 d.

##### *S9.4 Model comparison metrics*

Variants are compared by the Akaike information criterion in its small-sample form, with Akaike weights, since seventeen observations support four or five parameters. With $n_p$ the number of fitted parameters and $n$ the number of observations,

$$\mathrm{AIC} = 2\,\mathrm{NLL} + 2n_p, \qquad \mathrm{AICc} = \mathrm{AIC} + \frac{2n_p(n_p+1)}{n - n_p - 1}$$

$$w_m = \frac{\exp(-\Delta_m/2)}{\sum_j \exp(-\Delta_j/2)}, \qquad \Delta_m = \mathrm{AICc}_m - \min_j \mathrm{AICc}_j.$$

Out-of-sample performance is reported as the predictive NLL of a treatment withheld from calibration (§S14), and, for comparability with the evaluation practice of this literature, as normalised root-mean-square error and survival-probability prediction error, the EFSA metrics, and Nash–Sutcliffe efficiency on the withheld treatment.

##### *S9.5 How many damage stages a model must carry: the kT rule*

The choice between the single- and two-stage forms above is not arbitrary. Each stage in the chain from exposure to death carries a rate constant, and whether a stage has to be represented explicitly is decided by that rate against the length of the test. A stage whose rate is fast reaches equilibrium and can be absorbed into its neighbours; a stage whose rate is slow does not change within the test. Writing $T$ for the test length, the dimensionless product $kT$ sorts them:

- $kT > 20$ — the stage equilibrates and can be lumped;
- $0.2 < kT < 20$ — the stage must be carried explicitly;
- $kT < 0.2$ — the stage does not decay within the test.

The boundaries are deliberately wide: above $kT = 20$ less than $e^{-20}$ of the transient is left, so the stage is at equilibrium throughout; below $kT = 0.2$ less than 20% of the stage decays within the test, so it acts as a constant.

Applied at the four-day length of the regulatory acute battery, the criterion sorts every rate constant that has been measured for these compound classes.

**Table S22. Every measured stage of the damage chain against the four-day test.** Reproduced by `s09_stage_selection.py` (§S16).

| Stage | $k$ ($h^{-1}$) | Half-life | $kT$ | Verdict | Source |
|---|---|---|---|---|---|
| parent elimination, diazinon / *D. magna* | 0.266 | 2.6 h | 25.5 | lumped | ref 29 |
| **activation to diazoxon**, $k_{act}$ | 0.014 | 2.1 d | **1.3** | **carried** | ref 29 |
| diazoxon with acetylcholinesterase, $k_i$ | 3.3 | 0.21 h | 317 | lumped | ref 29 |
| carbaryl–enzyme reactivation, *in vitro* | 1.9 | 0.36 h | 182 | lumped | ref 31 |
| carbamylated enzyme hydrolysis | 1.19 | 0.58 h | 114 | lumped | ref 22 |
| **enzyme activity recovery, *in vivo*** | 0.12 | 5.8 h | **11.5** | **carried** | ref 31 |
| **thiacloprid receptor dissociation** | < 0.0036 | > 8 d | **0.3** | **carried** | ref 12 |
| **fitted upstream state $D_1$**, imidacloprid | 0.0189 | 1.53 d | **1.8** | **carried** | §4.3 |

**The criterion reproduces the modelling choices already made for these compounds.** One-compartment toxicokinetics suffice because parent elimination is lumped; the companion paper carries an activation compartment because $k_{act}$ is not;[4] binding is treated as instantaneous because $k_i$ is two orders of magnitude past the threshold; and the state this paper interposes is required because receptor dissociation for a neonicotinoid lies at the slow boundary of the band. The value fitted here falls in the middle of the band that has to be carried, which is a further check on it beyond the profile of §S15.2.

**What follows for the two hazard structures.** A slow upstream stage delays the peak of damage and so keeps a running maximum rising after exposure ends, which is what a peak-based hazard reads; a cumulative hazard integrates over the same delay and is less sensitive to it. Figure S6a shows that delay directly: as the upstream rate falls through the values measured for these compounds, the peak of damage moves from within the exposure to beyond the end of a four-day test, so that a running maximum is still being updated when the counts are taken. Figure S6b places every measured stage on the $kT$ axis, with the value fitted here among them. The peak-based structure's sensitivity to that delay, and the cumulative structure's insensitivity, are the whole of the reversal in §4.3. It also fixes the design requirement from the other side: a window shorter than three half-lives of the slowest carried stage, $\max_j 3\ln 2/k_j$ — 4.6 d for the fitted imidacloprid state — cannot show what those stages do (§4.7).

**Table S23. The stage-selection criterion applied by compound class.** Stages are carried or lumped by the $kT$ rule above at the four-day test length; the damage input $u$ is what the carried stages deliver.

| Class | Stages to carry | Damage input |
|---|---|---|
| uncouplers | none | $u = k_a C_b$, ending with exposure |
| carbamates | recovery ($kT \approx 11$) | $u = k_a C_b$ with fast $R$ |
| organophosphates | activation, and an adduct that ages | $u$ from the activated metabolite, activation rate-limiting |
| neonicotinoids | one persisting state ($kT \approx 0.3$) | $u = k_{12} D_1$, $D_1$ near frozen |

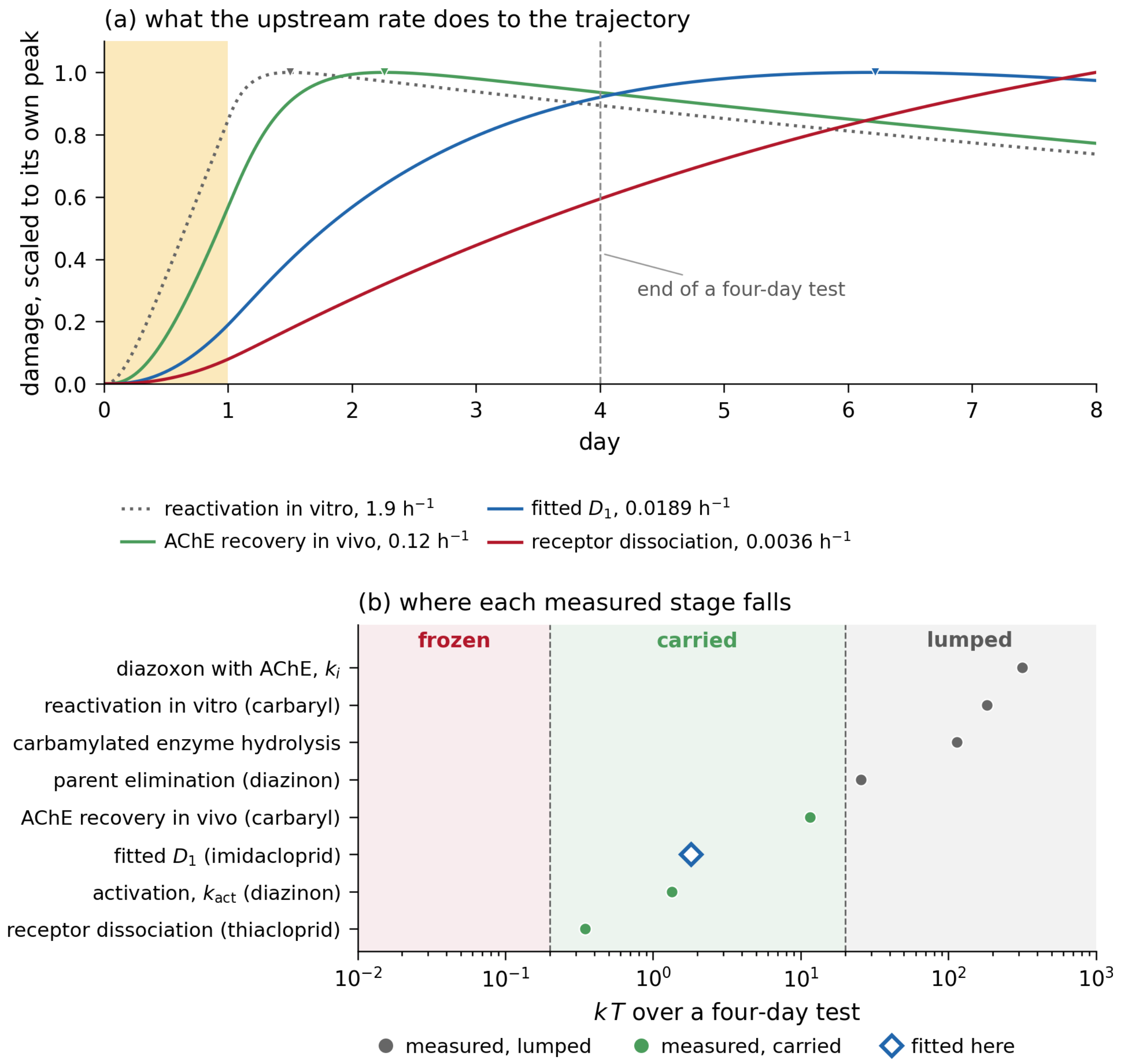


**Figure S6. The stage-selection criterion: simulation against measured rates.** (a) Damage under the two-stage model of §S9.1 following a one-day pulse (shaded), with the upstream clearance rate set in turn to four values that have been measured or fitted for these compounds; each trajectory is scaled to its own peak and the peak marked. As the rate falls the peak moves from within the exposure to beyond the end of a four-day test, which is what a running maximum reads and a cumulative hazard partly absorbs. (b) Every rate constant measured for this chain, placed on the $kT$ axis at the four-day test length. Bands mark the three regimes of §S9.5. Every stage that published models carry explicitly falls in the middle band and every stage they lump falls above it; the value fitted here sits in the middle of the band that has to be carried. Reproduced by `fig_S6_stages.py` (§S16).

##### *S9.6 Software and reproducibility*

All fits and figures were produced with Python 3.13.11 on Ubuntu 24.04 under Windows Subsystem for Linux 2 (NumPy, SciPy, Matplotlib; versions pinned in `requirements.txt` in the archive), by exponential-Euler integration at $\Delta t = 0.02$ d unless stated, with fixed random seeds.

---

#### S10. Numerical Verification of Both Dose Metrics

This section collects the numerical checks of the propositions of §S1 and of the cumulative metric, for which no proof is available (§S1.2), and asks of each whether any configuration reproduces the observed ordering of §4.6, in which mortality rises as the pulses are moved apart. Four checks are reported. At short separations, overlapping designs included, the peak is computed under ten repair laws (next paragraph). Table S24 runs the peak metric under six repair laws on a generic pair of two-day pulses. Table S25 scans both metrics over a parameter grid on the three exposure splittings of §4.6 — one 48-hour pulse, two 24-hour pulses seven or fourteen days apart — 81 combinations per metric, 162 in all. Figure S7 shows both metrics as continuous functions of the separation, from complete overlap to 21 d. §S10.1 and §S10.2 then ask what it would take to break Propositions 2 and 3. Where a law satisfies the hypotheses of §S1, the peak metric is settled by Proposition 2 and the numbers show the size of the effect; for the cumulative metric the numbers are the only evidence. Three of the cases checked lie outside those hypotheses and are covered by the numbers alone: square-root repair, which is not Lipschitz at zero; the band laws of Table S26, which switch on and off discontinuously; and a rectangular exposure applied to damage directly, with no first-order stage, which is log-concave but vanishes after the pulse and so is a limit of condition (V1) rather than an instance of it. Constant removal, which does not vanish as damage goes to zero, is covered by Proposition 2 as a limit of laws that satisfy its hypotheses (ref 35, §1).

**Short separations, overlapping designs included.** Proposition 2 is least obvious where the two pulses overlap (§S1.2). Two rectangular pulses of unit height and duration $\tau = 1$ d were placed with their starts $\Delta = 0$, 0.25, 0.5, 0.75, 1, 2, 4, 7 and 14 d apart — from complete superposition through the abutting design to the separations of §4.6 — under ten repair laws: proportional at four rates from 0.01 to 5 $d^{-1}$, Michaelis–Menten in two regimes, constant removal, square-root, threshold, and non-monotone ($1.5\,D\,e^{-D}$). The peak was non-increasing in $\Delta$ under every law, and for proportional repair the computed ratio $M(0)/M(\tau)$ matches the closed form $2/(1+e^{-k_r\tau})$ to four decimals at all four rates. Reproduced by `s01_prop2_short_separations.py` (§S16).

**Table S24. Repair laws tested for the peak metric.** Two pulses of duration $\tau$ = 2 d with their starts $\Delta$ apart, $\Delta$ from 2 to 30 d; at $\Delta = \tau$ the pulses abut, which is the single-pulse design. $f_{max}$ is the peak reached by one pulse alone, as in §S1. Under every law the peak is non-increasing in $\Delta$. The last two columns are the smallest ratio over the range computed, $M(30)/M(\tau)$, and its limit as $\Delta \to \infty$, $f_{max}/M(\tau)$, which Proposition 3 bounds below by one half for convex repair. Reproduced by `s10_metric_scan.py` (§S16).

| Repair law | $M(\Delta{=}\tau)$ | $M(30)$ | Monotone non-increasing | max $M/f_{max}$ | $M(30)/M(\tau)$ | $f_{max}/M(\tau)$ |
|---|---|---|---|---|---|---|
| proportional, $R = 0.3D$ | 2.330 | 1.504 | yes | 1.55 | 0.65 | 0.65 |
| Michaelis–Menten, saturated ($R_{max} = 0.05$, $K = 0.01 \ll D$) | 3.803 | 2.417 | yes | **2.00** | 0.64 | **0.50** |
| Michaelis–Menten, transition ($R_{max} = 0.2$, $K = 0.5$) | 3.434 | 1.769 | yes | 1.94 | 0.52 | 0.51 |
| quadratic, $R = 0.5D^2$ | 1.404 | 1.270 | yes | 1.12 | 0.90 | 0.89 |
| square-root, $R = 0.3\sqrt{D}$ | 2.630 | 1.493 | yes | 1.76 | 0.57 | 0.57 |
| threshold, $R = 0.4[D-0.2]^+$ | 2.154 | 1.577 | yes | 1.45 | 0.73 | 0.69 |

The last two columns test Proposition 3. For convex repair the floor of one half is proved (§S1.3); concave laws are not covered by the proof. The saturated Michaelis–Menten row, close to constant removal, sits exactly on the floor because the pulses in this table do not overlap and the input has no tail. Overlap (Figure S7a) or a decaying input tail (Table S27: $M(\tau)/f_{max}$ = 2.681, ratio 0.373) takes a concave law across it, which is why Proposition 3 is stated for convex repair.

**Table S25. Combinations tested, and the number reproducing the observed ordering.** "Reduced" is the one-compartment scaled-damage model with dominant rate constant $k_d$; "two-stage" is the model of §S9.1. "Reversals" counts combinations in which mortality rises as the pulses are moved apart. Grids: seven values of $k_d$ log-spaced over the range and three thresholds for the reduced model (7 × 3 = 21); four values of $k_{off}$, five of $R_{max}$ and three of $K/D_{peak}$, at a threshold of 0.2 of the single-pulse peak, for the two-stage model (4 × 5 × 3 = 60); each counted once per metric. Reproduced by `s10_metric_scan.py` (§S16).

| Hazard structure | Damage model | Parameters scanned | Combinations | Reversals |
|---|---|---|---|---|
| peak-based | reduced | $k_d \in [0.02, 5]$ $d^{-1}$, threshold 0.05, 0.2 or 0.5 of peak | 21 | 0 |
| cumulative | reduced | as above | 21 | 0 |
| peak-based | two-stage, MM repair | $k_{off} \in [0.05, 1]$, $R_{max} \in [0.002, 0.2]$, $K/D_{peak} \in \{10^{-6}, 0.1, 1\}$ | 60 | 0 |
| cumulative | two-stage, MM repair | as above | 60 | 0 |
| **total** | | | **162** | **0** |

The $K/D_{peak}$ values scanned in Table S25 run from the fully saturated limit ($K \ll D$, where removal is constant) to $K \approx D$, the regime in which the *D. magna* optimum falls, though $K$ itself is not identified there (§S15.2). The first-order limit, in which Michaelis–Menten repair becomes proportional, is covered by the proportional law of Table S24 and by the reduced-model scans. Figure S7 shows both statistics as continuous functions of the separation, for proportional and Michaelis–Menten repair, from complete overlap to 21 d.

**Slow cascades tested separately.** Because a sufficiently slow second stage delays the damage maximum, and a maximum that occurs after the later pulse might in principle invert the ordering, second-stage rate constants from 0.005 to 0.3 $d^{-1}$ were scanned separately, moving the damage maximum from day 3.7 to day 10.4. The ordering of the peak metric held in all eight cases, as Proposition 2 requires of a linear cascade; delaying the maximum changes when damage is largest, not the monotonicity in the interval between pulses.

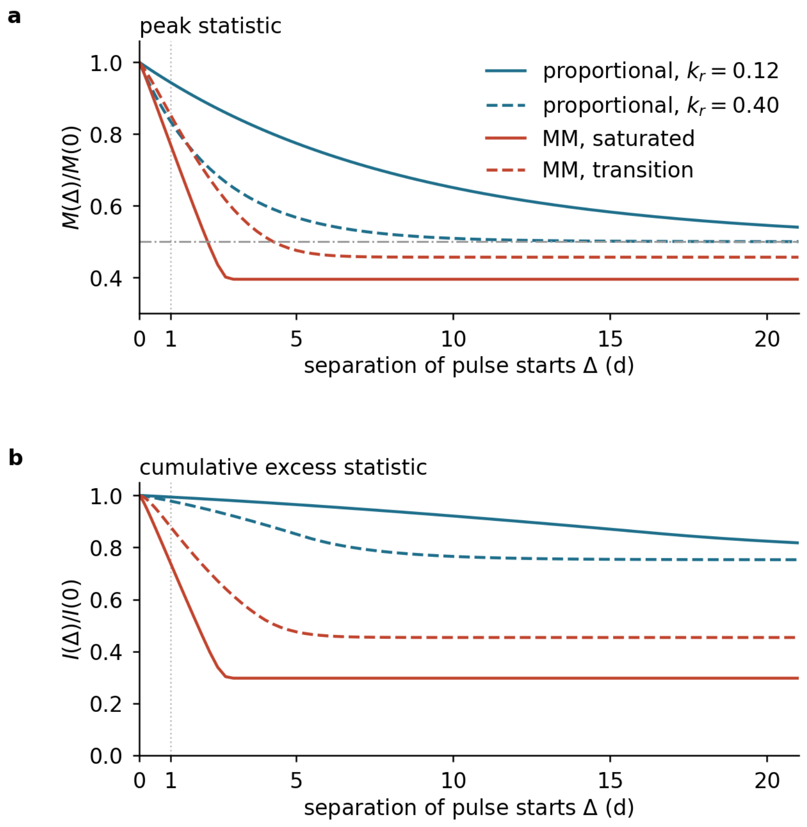


**Figure S7. Numerical verification of the two dose metrics.** (a) The peak statistic and (b) the cumulative excess statistic, $I = \int [D - z]^{+}\,dt$ with $z = 0.15$, as functions of the separation $\Delta$ between the starts of two identical 24-hour pulses, from complete overlap ($\Delta$ = 0) through the abutting design ($\Delta$ = 1 d, dotted) to 21 d, each relative to its value at complete overlap. Repair is proportional at two rates, or Michaelis–Menten with $R_{\max}$ = 0.4 $d^{-1}$ in the saturated and transition regimes; the input is the rectangular exposure itself, a limit of condition (V1) (§S10, opening). Both statistics are non-increasing throughout under all four laws: for the peak this is Proposition 2, and for the cumulative statistic, for which no proof is available (§S1.2), the panel is the evidence. The dash-dotted line is the one-half bound of Proposition 3. Proportional repair approaches it as recovery between pulses becomes complete; both Michaelis–Menten curves cross it, because that law is concave and two overlapping pulses raise damage above twice the single peak (§S1.3, §S10.2) — at a maximum repair rate far above the one fitted (Table S28). Drawn by `fig_S5_S7.py` (§S16).

##### *S10.1 What it would take to break Proposition 2, and whether it is enough*

Proposition 2 holds under (V1)–(V3), and (V3), log-concavity of the damage input, is where a real system can escape it: two-compartment kinetics make the input decay as a sum of two exponentials, $A_1 e^{-k_f t} + A_2 e^{-k_s t}$, which is log-convex rather than log-concave in the tail. Three results bound how large a violation that escape can produce, a fourth shows why no qualitative rule can exclude it, and a grid illustrates them.

**Linear repair closes the escape.** For linear repair, $R(D) = \kappa D$, Proposition 2 needs only a unimodal single-pulse input, which a rectangular pulse through one- or two-compartment elimination provides (ref 35, Theorem 1 and Lemma 1): the peak is non-increasing in the separation, log-convex tails included. A violation therefore requires a repair law that is nonlinear over the range the trajectory visits, which is the first thing to check of any candidate.

**A sandwich bound turns that into a test.** If the single-pulse input is unimodal and, on $[0, M^{\kappa_-}(\Delta)]$, the law is enclosed between two linear ones, $\kappa_- D \le R(D) \le \kappa_+ D$, where $M^{\kappa}$ is the peak under linear repair at rate $\kappa$, then comparison with the two linear trajectories gives

$$\frac{M_R(\Delta')}{M_R(\Delta)} \;\le\; \rho \equiv \frac{M^{\kappa_-}(\Delta')}{M^{\kappa_+}(\Delta)}$$

(proof: ref 35, Theorem 2). When $\rho \le 1$ the pair of separations cannot reverse; the test takes three linear integrations and applies to one pair of separations at a time (ref 35, Corollary 2.1). For the threshold law with an added linear term, $R(D) = 0.1\,D + 0.5\,[D-c]^+$, $\rho(1\text{ d}, 14\text{ d}) = 0.96$, so under that law the fourteen-day design cannot exceed the single 48-hour pulse; the test does not decide the pairs (1 d, 7 d) and (7 d, 14 d) ($\rho = 1.028$ and $1.020$ at the tightest admissible $\kappa_+$; ref 35, §8).

**Threshold repair has a ceiling computable from linear integrations.** Where the test gives $\rho > 1$, the excess above the switch level $c$ is itself governed by a linear process, and ref 35, Proposition 3, bounds the peak ratio by $(c+M^r(\Delta'))/M^{\kappa_+}(\Delta)$, where $M^r$ is the peak of that linear process, with rate $r$, and $M^{\kappa_+}$ the peak under a linear law that bounds the repair law from above; for a law that removes nothing below $c$ it is at most $1 + M^r(\Delta')/c$. For the hard threshold law $0.5\,[D-c]^+$ with $c = 0.553$, on the input that gives the largest violation found, this is 1.515 for the fourteen-day design and 1.553 for the seven-day design, against computed ratios of 1.241 and 1.165; both bounds rest on linear integrations alone.

**No qualitative rule separates violating repair laws from safe ones.** Adding a linear term $\varepsilon D$ to the hard threshold law leaves the violation in place at small $\varepsilon$: the computed ratio is 1.240, 1.236 and 0.906 at $\varepsilon = 10^{-3}$, $10^{-2}$ and $10^{-1}$, against 1.241 for the hard threshold, and the sandwich certificate applies at $10^{-1}$. These values are numerical. Ref 33, Proposition 4, proves the qualitative fact behind them: given one violating law, every law close enough to it that removes at least as much damage also violates, so no property of where repair is zero separates violating laws from safe ones.

**Table S26. Proposition 2 violations under two-compartment kinetics, by repair family.** $M(1\,\mathrm{d})$ is the peak under the single 48-hour pulse, $M(14\,\mathrm{d})$ under two 24-hour pulses fourteen days apart; a ratio above one is a violation. Numerical illustration of the results above, over 5,568 combinations of a two-compartment input (slow-pathway dose share $\varphi$, rates $k_s$ and $k_f$) with five repair families, on the design of §4.6. The log-concave control, $\varphi = 0$, gives a largest ratio of 1.0007 over 58 configurations at the time step used and 1.0003 at half that step. The excess comes only from the band laws, which switch on and off discontinuously and so fall outside the Lipschitz condition of Proposition 2; every continuous law in the control stays at 1.0000, as the proposition requires. A refined grid ($\varphi \le 0.8$, $k_f \le 20\ \mathrm{d}^{-1}$) raises the largest violation with the observed monotone ordering to 1.241, stable to 1.240 on a quarter step — below the bound of 1.515 that ref 35, Proposition 3, gives from linear integrations (§S10.1).

| Repair family | Violations | Largest $M(14\,\mathrm{d})/M(1\,\mathrm{d})$ |
|---|---|---|
| proportional | 0 | — |
| Michaelis–Menten | 0 | — |
| non-monotone | 0 | — |
| threshold, $r[D-c]^+$ | 517 | **1.168** |
| band, $r\,\mathbf{1}[c<D<c+w]$ | 638 | 1.046 |

The two repair laws of the classification never violate it, as the linear theorem requires of the proportional one and the grid confirms of the other. The violation grows with $\varphi$ to a maximum near 0.6–0.7 and falls again (Figure S8a).

**A small violation of the peak is enough to reproduce the observed ordering.** Taken at face value a peak ratio of 1.241 is small, but the observed quantity is the excess above a threshold, and the amplification near $\theta = M$ is steep (Figure S8b). Reaching the observed excess-damage ratio of 3.3 needs $\theta = 0.895\,M(1\,\mathrm{d})$; at $\theta = 0.90$ the ratio is 3.41 and the *A. aquaticus* mortality goes from 0.215 to 0.563 against the 0.554 observed. The ordering of §4.6 is therefore reproducible outside (V3) — by a two-compartment elimination with roughly two-thirds of the dose on a slow pathway, a fast-to-slow rate ratio near 400, threshold repair, and a killing threshold at nine-tenths of the smaller peak.

**What reproducing the ordering outside (V3) requires.** Each of those four is a claim about the animal, and three are checkable: a slow pathway carrying most of the dose is a toxicokinetic statement, the rate ratio is measurable, and a threshold at $0.9\,M$ fixes the slope of the concentration–response. None has been measured for these species. The conclusion of §4.6 is therefore that the ordering lies outside the framework as fitted — with the one-compartment kinetics and the two repair laws the classification carries — rather than outside anything the framework could be extended to express.

Reproduced by `s10_prop2_violation_search.py` and `s10_sandwich_certificate.py` (§S16), which carry the control at two time steps, the refined grid, the step-size check, the certificate example and the softening series.

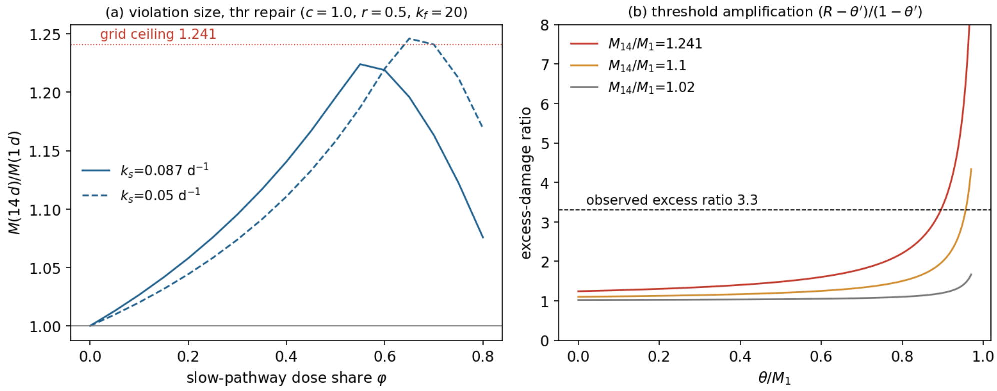


**Figure S8. How large a Proposition 2 violation two-compartment kinetics can produce.** (a) The peak ratio $M(14\text{ d})/M(1\text{ d})$ under threshold repair as a function of the slow-pathway dose share $\varphi$, for two slow rates; the violation grows to a maximum near $\varphi$ = 0.6–0.7 and falls again, and the refined grid ceiling is 1.241; the $k_s$ = 0.05 curve, sampled more finely in $\varphi$, reaches 1.246 at $\varphi$ = 0.65. (b) The excess-damage ratio $(R-\theta')/(1-\theta')$ as the threshold approaches the smaller peak, $\theta' = \theta/M(1\text{ d})$, for three peak ratios, with the observed 3.3 dashed. The dotted line in (a) is the grid ceiling of 1.241; the bound of 1.515 from linear integrations (§S10.1) lies above the axis. The observed 3.3 is reached at $\theta'$ = 0.895 for the largest violation the grid allows, and not at all for a peak ratio of 1.02. Drawn by `fig_S8_violation.py` from the model of `s10_prop2_violation_search.py` (§S16); see §S10.1.

##### *S10.2 Where the factor-two step fails, and whether the fits reach it*

Proposition 3 rests on an intermediate step, $M(\Delta) \le 2f_{\max}$, proved in §S1.3 for convex repair: such a law removes at least as much from a combined level as from its parts, $R(a+b) \ge R(a) + R(b)$, so two pulses never push damage above the sum of two single-pulse trajectories. For proportional repair the inequality is an equality. A concave law removes less from a combined level than from its parts, so the step can fail, in two ways. Overlapping pulses double the input while concave removal rises less than twofold, which is why both Michaelis–Menten curves of Figure S7a cross the bound. Pulses that do not overlap breach it only through the input tail: with a tail-free input the zero-order limit sits exactly on the bound (Table S24), while a decaying tail costs the single-pulse peak more removal than it costs the double, and the excess appears (Table S27).

**Table S27. The factor-two step and the one-half bound, by the shape of the repair law.** A single 24-hour pulse against two such pulses on a one-compartment input, so that (V1)–(V3) hold throughout; $\tau$ is the contiguous design. Values are those printed in section [4] of the output of `s10_sandwich_certificate.py` and are stable to a quarter step. Bold marks a bound breached.

| Repair law | Shape | $M(\tau)/f_{\max}$ | $M(14)/M(\tau)$ |
|---|---|---|---|
| proportional, $k_r$ = 0.05 | linear | 1.988 | 0.769 |
| threshold, $r[D-c]^+$ | convex | 1.648 | 1.000 |
| square-root, $R = 0.3\sqrt{D}$ | concave | **2.212** | **0.452** |
| constant removal, $R$ = 0.05 | concave | **2.206** | 0.629 |
| Michaelis–Menten, $K = 0.01\,D_{\text{peak}}$ | concave | **2.681** | **0.373** |

**The fitted repair rate is far from the failure.** Michaelis–Menten repair is concave, and it is one of the two laws the classification carries, so the question is whether the breach is reachable at the repair rates this archive supports. It is not: in deep saturation — the worst case for this law, and a regime the fits cannot exclude since $K$ is not identified (§S15.2) — the one-half bound is breached only for $R_{max}$ between 0.05 and 0.2 $d^{-1}$ or above (Table S28), and the value fitted to the *Daphnia magna* series is $8 \times 10^{-5}$ $h^{-1}$, or 0.0019 $d^{-1}$, at least twenty-six times smaller, where the ratio is 0.99.

**Table S28. The one-half bound against the maximum repair rate, Michaelis–Menten in deep saturation.** The rate fitted in §S11 sits at the right-hand end.

| $R_{max}$ ($d^{-1}$) | **0.200** | **0.050** | **0.020** | **0.005** | **0.002** |
|---|---|---|---|---|---|
| $M(14)/M(\tau)$ | **0.36** | 0.63 | 0.87 | 0.97 | 0.99 |

The bound §4.6 uses is therefore safe for this archive, and the convexity hypothesis records what would have to be true of a compound for it not to be. Reproduced by `s10_prop3_factor_two.py` (§S16).

#### S11. The *Daphnia magna* Series: Fits, Parameters, and What Each Extension Contributes

**The series and what this section gives.** Li *et al.* exposed *Daphnia magna* to imidacloprid in three treatments that share a first 24-hour pulse and differ only in when the second arrives — 2, 7 or 15 days later — with seven replicates of ten animals and seventeen survival observations over 22 d.[6] This section gives every model variant fitted to that series (Table S29), the parameters of the two that Figure 2 shows (Table S30), what each extension contributes (Table S31), where the fitted persistence sits among measured ones (§S11.1), and the parameter-free comparison of §4.2 step by step (§S11.2).

**Table S29. All model variants, fitted to treatment C alone and to all three treatments with a single parameter set.** All values are warm-started optima; the likelihood surface has multiple local optima and warm starts are decisive (§S9), only one restart in thirty reaching the best basin unaided. With seventeen points and four or five parameters the small-sample criterion is the appropriate one: AICc and the Akaike weights *w* are given for the three-treatment fit. The ordering is the same under AIC and AICc; the margin over the best cumulative variant is 10.5 AICc units, with 89% of the weight on the two-stage peak-based model. Bold marks the best value in each column.

| Damage | Repair | Hazard | Par. | Treatment C only: NLL | AIC | All three: NLL | AIC | AICc | ΔAICc | *w* |
|---|---|---|---|---|---|---|---|---|---|---|
| single | proportional | cumulative | 3 | **112.84** | **231.67** | 284.97 | 575.94 | 577.79 | 10.5 | 0.005 |
| single | proportional | peak-based | 3 | 162.30 | 330.61 | 363.09 | 732.19 | 734.03 | 166.8 | < 0.001 |
| two-stage | proportional | cumulative | 4 | 113.73 | 235.46 | 285.25 | 578.50 | 581.83 | 14.6 | 0.001 |
| two-stage | proportional | peak-based | 4 | 118.83 | 245.65 | **277.97** | **563.95** | **567.27** | **0** | **0.891** |
| two-stage | MM | cumulative | 5 | 113.68 | 237.36 | 285.34 | 580.69 | 586.13 | 18.9 | < 0.001 |
| two-stage | MM | peak-based | 5 | 111.31 | 232.62 | 278.06 | 566.13 | 571.57 | 4.3 | 0.104 |
| two-stage | proportional | cumulative on $D_1$ | 4 | — | — | 284.97 | 577.94 | 581.27 | 14.0 | 0.001 |

**Giving the cumulative structure its own extension does not change the result.** The last row of Table S29 reads the cumulative threshold on the upstream state $D_1$. With the same four parameters and warm starts it returns NLL 284.97, the single-stage cumulative value, with the transfer rate at the upper bound of its search range, and falls 14 AICc units behind. Reproduced by `s11_symmetric_extension.py` (§S16).

**Fitted to one treatment alone, the verdict reverses.** Fitted to treatment C alone, the simplest cumulative model has the lowest information criterion and the four-parameter peak-based model — which wins by 12 information criterion units on the three treatments together — trails every cumulative variant and its own saturating-repair form, ahead only of the single-stage peak-based model.

The published parameter set of Li *et al.* gives NLL 289.06 on the three treatments, so the fitted single-stage cumulative model improves on it by 4.1 units before any structural question is asked. Table S30 gives the fitted parameters of the two-stage fit under each hazard structure.

**Table S30. Fitted parameters of the two-stage fit under each hazard structure.** Proportional repair for the cumulative structure, Michaelis–Menten for the peak-based one. These are the variants of Figure 2(c, d) up to the repair law, which moves every other parameter by less than 2% (Table S31). Half-lives are converted from the fitted rate constants, which are per hour as in the source implementation.

| | Upstream half-life | Terminal repair half-life | Damage maximum | Effect of the extension |
|---|---|---|---|---|
| cumulative, two-stage proportional | 0.7 h (at the bound of the search) | 6.1 d | day 1.2 | collapses onto the single-stage model |
| **peak-based, two-stage Michaelis–Menten** | **1.50 d** | — | **still rising at day 22** | improves on the single-stage model by 85 units |

A dash in the table means the profile puts the rate below the resolution of the test: terminal repair is localised only to half-lives beyond $10^2$ d (§S15.2). The cumulative fit discards the extension — its upstream state clears within the hour, so $D_1$ simply follows the parent — which is why its likelihood matches the three-parameter value to within 0.3 units. The peak-based fit uses it, and uses it in one specific way: a state that outlasts the parent compound by more than a day, feeding the damage $D_2$ that the hazard reads, which is not repaired within the test.

**How much of each test the running maximum spends frozen.** On the fitted trajectory of the two-stage peak-based variant the maximum is reached on day 15.0, 19.9 and 22.0 for the two-, seven- and fifteen-day treatments, so the hazard is frozen for 32%, 10% and 14% of the three tests — in the last of these only between the pulses. Where the maximum is still updating the two hazard structures are the same functional of the exposure (§S12), so the twelve-unit margin of Table S29 measures the damage model rather than a property the peak-based structure has and the cumulative one lacks. Reproduced by `s11_frozen_time.py` (§S16).

**Table S31. Contribution of each extension, evaluated against the cumulative structure refitted at the same damage model.** The likelihood columns decompose the comparison of Table S29; “vs cumulative” is the cumulative NLL minus the peak-based NLL, so a positive value favours the peak-based structure. The fitted parameters are added here. Rate constants are per hour, as in the source implementation.

| Damage model | Repair | Par. | Peak-based NLL | vs cumulative | $k_{off}$ ($h^{-1}$) | $k_r$ or $R_{max}$ ($h^{-1}$) | $k$ | $\theta$ |
|---|---|---|---|---|---|---|---|---|
| single stage | proportional | 3 | 363.09 | − 78.1 | — | $1.0 \times 10^{-4}$ | 1.00 | 0.0067 |
| **two stage** | **proportional** | **4** | **277.97** | **+ 7.3** | **0.0189** | $\mathbf{7.0 \times 10^{-5}}$ | **2.64** | **0.449** |
| two stage | MM | 5 | 278.06 | + 7.3 | 0.0193 | $8.0 \times 10^{-5}$ | 2.69 | 0.446 |

**The extension does not help every variant.** Given the same upstream stage, the cumulative structure ends 0.3 likelihood units *worse* than its own single-stage special case under the repair law used in Figure 2: a metric that integrates over time gains nothing from a later peak (Figure 2a,c). Adding saturating repair on top of the two-stage model changes the likelihood by 0.09 and costs a parameter, so the information criterion worsens.

The two-stage rows differ in the fifth parameter only: the Michaelis–Menten fit adds $K = 1.0$, which the profile of §S15.2 shows to be unidentified. Every other parameter moves by less than 2%. Here $k$ is per unit damage and $\theta$ is in damage units; the damage scale is arbitrary and fixed by $k_a$, so neither is comparable with $b_w$ and $z$ (§S9).

##### *S11.1 Where the fitted persistence sits among measured ones*

The state $D_1$ is inferred from survival counts, not measured. Whether the number it takes is physically reasonable can still be asked, by placing it on the axis of persistences that have been measured in these animals.

**Table S32. The fitted persistence against measured ones.** Half-lives, not rate constants, for comparability.

| **Quantity** | **Half-life** | **Species, compound** | **Source** |
|---|---:|---|---|
| parent internal residue | 0.11–4.62 d | *G. pulex*, five classes | refs 25–27 |
| parent internal residue | 0.04–0.05 d (0.60–0.78 $h^{-1}$) | *D. magna*, imidacloprid | ref 6 |
| **measured** recovery of inhibited enzyme, *in vivo* | **0.24 d** (0.12 $h^{-1}$) | *D. magna*, carbaryl | ref 31 |
| **fitted upstream state $D_1$** | **1.53 d** (1.13–2.41) | *D. magna*, imidacloprid | §4.3 |
| upstream state inferred from a ratio of counts | ≥ 5.3 d (lower bound) | *C. dipterum*, imidacloprid | §S8 |
| receptor-bound fraction | > 8 d | *G. pulex*, thiacloprid | ref 12 |

**The fitted value falls between the parent and the receptor, which is where a damage precursor belongs.** It is slower than the compound leaves the animal and faster than it dissociates from the receptor, so it cannot be either of those two things and is not required to be. The one directly measured entry — enzyme activity recovering *in vivo* at 0.12 $h^{-1}$ in the same species[31] — is for a carbamate, the class whose lesion is shortest-lived, and sits below the fitted neonicotinoid value as the ordering of §4.5 requires.

**For anticholinesterases the precursor is measured directly, and the measured range spans three orders of magnitude in the order of §4.5.** Enzyme activity is itself the state that persists after the parent has cleared: after a carbaryl pulse it returns to control activity within hours in *Chironomus riparius*[22] and recovers at 0.12 $h^{-1}$ *in vivo* in *D. magna*,[31] whereas for the organophosphates of the midge experiment no reactivation was detected and the inhibited enzyme ages into a form that cannot reactivate.[22] At the persistent end there is no rate to recover, because the enzyme does not return.

**What is still missing.** No study we found measures $D_1$, or a quantity standing for it, for imidacloprid in *D. magna*; the persistence reported here is inferred from counts.

##### *S11.2 The parameter-free comparison of §4.2, step by step*

The prediction of §4.2 uses no fitted quantity, and the steps are these.

**Table S33. The four numbers behind the 68% prediction.**

| Step | Value | Source |
|---|---|---|
| survival one day after the first pulse | 89% | ref 6, Figure 2 |
| survival at day 10, constant thereafter until the second pulse | 24% | ref 6, Figure 2 |
| day on which the running maximum becomes fixed | 1.14 d | published trajectory (ref 6); the refits of Figure 2 freeze at 1.1–1.3 d |
| control survival at 22 d | 76% | ref 6 |

While the running maximum is fixed, the peak-based hazard is constant, so the predicted survival ratio between any two times in the interval equals the ratio in the controls. Allocating the whole of the 22-day background mortality to the interval from day 1 to day 10 is the assumption most generous to the peak-based structure, because it puts as much decline as the controls allow into that interval; even so the prediction is 68% where 24% was observed, a factor of 2.8. The threshold, the killing rate and the tolerance spread cancel from the ratio whatever values they take, which is why no fit can close the gap; the fitted counterpart (Table S29, Figure 2b) is reported as a check rather than as the result.

#### S12. The Two-Stage Extension Applied to Two More Neonicotinoid Series and to an Uncoupler

**What this section adds.** §4.3 and §4.5 refer to two comparisons they do not tabulate: the two-stage extension applied to two more species exposed to imidacloprid, and the four variants fitted to the uncoupler DNOC. The two species are *Asellus aquaticus* and *Caenis horaria* from the interval series of §S7, each fitted with one parameter set across its three exposure profiles, damage driven by the nominal external concentration, and background mortality from the concurrent control (`s12_two_species_fits.py`, §S16).

**Table S34. Single- and two-stage damage under both hazard structures, one parameter set per species.** Values are negative log-likelihoods. The deficit is the single-stage peak-based value minus the single-stage cumulative one, so a positive deficit means the peak-based structure is behind. The *D. magna* row is from Table S29.

| Species | Single-stage deficit of the peak-based structure | Two-stage cumulative | Two-stage peak-based, proportional repair |
|---|---|---|---|
| *A. aquaticus* | 59.2 | 430.14 | 431.10 |
| *C. horaria* | − 3.5 | 526.36 | 520.95 |
| *D. magna* (for comparison) | 78.1 | 285.25 | 277.97 |

**In *A. aquaticus* the two structures coincide.** On a single damage stage the peak-based structure falls 59 units behind the cumulative one; with the upstream stage it reaches 431.10, the single-stage cumulative value to two decimal places and within one unit of the two-stage cumulative fit. The reason is structural rather than statistical. Both fits drive the recovery rates to zero, and in that limit a peak-based hazard on a twice-integrated damage cascade and a cumulative hazard on a once-integrated one are the same functional of the exposure — the isomorphism of the companion paper, the two structures coinciding once repair vanishes,[4] reached here exactly. No likelihood can then separate the structures, and a saturating repair term has nothing to act on.

**In *C. horaria* the peak-based structure leads at either depth.** It is ahead by 3.5 units on one damage stage and by 5.4 on two, in line with the structure the literature assigns this species (Figure 1); the cumulative structure gains nothing from the added stage.

**What the data require is the upstream stage, not saturating repair.** Saturation changes the log-likelihood by 0.00, 0.00 and $-0.09$ units in the three series, in each case less than the one likelihood unit (two AIC units) the extra parameter costs.

**The uncoupler: what the upstream stage is worth when damage does not outlast the compound.** The four variants — single- and two-stage damage under each hazard structure, the two-stage form with saturating repair — are fitted to the three DNOC interval treatments simultaneously, with the elimination rate constant estimated for this compound and species ($k_e = 1.11$ d$^{-1}$, a half-life of 0.62 d;[25] Table 1 there), and give the values behind Figure 4. Under the single-stage trajectory the cumulative structure gives NLL 136.34 and the peak-based one 120.78 (three parameters each, AIC 278.7 and 247.6); with the upstream stage the two give 136.34 and 120.47 (five parameters each, AIC 282.7 and 250.9). Both two-stage fits are warm-started from the single-stage optimum (§S9.3).

The upstream stage improves the peak-based log-likelihood by 0.3 units for DNOC, less than its two extra parameters cost (AIC +3.4), against a gain of 166 AIC units for imidacloprid in *D. magna* (single stage to two-stage Michaelis–Menten, Table S29); it leaves the cumulative fit unchanged (AIC +4.0). At its best the cumulative structure is 31 units behind the peak-based one with the same damage model. With damage that does not outlast the compound, there is nothing for the extension to carry.

---

#### S13. Why an Eroding Tolerance Cannot Produce the Interval Reversal, and What Could

**What is being tested.** A tolerance that erodes during the test, so that an animal becomes easier to kill as time passes, is the one remaining candidate inside the threshold itself; feeding inhibition is documented for this compound class at concentrations roughly a fifth of the lethal one.[10,11,33] This section asks whether erosion can make the fourteen-day treatment of §S7 deadlier than the single 48-hour pulse, in either of the two ways it can enter the dose metric.

**Applied after the running maximum, erosion changes nothing.** Let $g(t)$ be the declining factor by which the tolerance erodes. Scoring $M(T)/g(T)$ divides every exposure profile by the same constant at the end of the test, which is a monotone rescaling and preserves every ordering. Placing the erosion outside the running maximum is, however, an approximation; the definition puts it inside, as the next paragraph does.

**Applied inside the running maximum, it turns the ordering over by only 1.3%.** An individual with threshold $\theta_i\, g(t)$ dies when $D(\tau) > \theta_i\, g(\tau)$ for some $\tau \le t$, which is equivalent to $\theta_i < \max_\tau[D(\tau)/g(\tau)]$, so the correct dose metric is the running maximum of the *ratio*. With $g(t) = e^{-\lambda t}$ this is $\max_\tau[D(\tau)e^{\lambda\tau}]$, and as $\lambda$ rises the maximum moves towards the end of the observation window. What happens there depends on how fast damage recovers. With the recovery rate the *A. aquaticus* fit returns — driven to zero (§S12) — the profile whose exposure ended latest ends the window marginally higher in the ratio metric: the fourteen-day profile exceeds the single pulse by 1.3%, so the ordering does turn over, but only just. The observed excess-damage ratios of the fourteen-day treatment to the single pulse, control-corrected, are 3.3 for *A. aquaticus* and 1.6 for *P. minutissima* at day 21. A 1.3% effect cannot account for either, and the turnover does not grow with $\lambda$ above about 0.03 d$^{-1}$, so no choice of erosion rate improves it.

**What reaching the observed ratios would take.** The effect becomes large enough only if damage is also allowed to recover appreciably: with a terminal repair half-life near six days ($k_r \approx 0.12$ d$^{-1}$) and an erosion rate near $\lambda = 0.12$ d$^{-1}$, the predicted ratios reach the observed ones. The repair half-life required is some sixty times shorter than the value the *Daphnia magna* profile allows — about 350 days, $k_r \approx 0.002\ \mathrm{d}^{-1}$ (§S15.2) — and the fit to *A. aquaticus* itself returns no measurable repair (§S12). An eroding tolerance therefore reproduces the ordering only by assuming a recovery rate the same data exclude.

**What the data require instead.** Once the repair laws and an eroding threshold are excluded, what is left is a state depleted during exposure and only partly restored in clean water — non-monotone, and therefore an explicit energy budget rather than a damage cascade or a rescaled threshold. Two measurements from the same laboratory support that reading: carbaryl consumes 26% of the available energy budget in *Daphnia magna*,[31] and the diazinon recovery experiment was run without feeding, which its authors note would have hastened recovery.[29] It is also consistent with the sublethal literature, in which starvation mortality is reported as an indirect cause adding to direct mortality rather than modifying it.[10,11]

**How to test it: vary the food.** The state carried between pulses is invisible in both recorded endpoints, so neither survival nor immobilisation can be read for it directly. If depleted reserves are what the first pulse leaves, raising food should weaken the interval reversal and restricting food strengthen it, whereas an explanation internal to the damage cascade predicts no effect either way. That experiment has not been run.

---

#### S14. Out-of-Sample Prediction: Each Treatment Predicted from the Other Two

The three treatments share a first pulse and differ only in when the second arrives, so each can be held out in turn and predicted from the other two. This tests whether a fitted structure extrapolates to an interval it has not seen, which is the property the calibration–validation exercises in this literature are built to assess. Alongside the predictive likelihood, the two metrics used in EFSA Tier-2C model evaluation are reported, normalised root-mean-square error and survival-probability prediction error,

$$\mathrm{NRMSE} = \frac{1}{\bar{S}_{\mathrm{obs}}}\sqrt{\frac{1}{n}\sum_i \left(S_{\mathrm{obs},i} - S_{\mathrm{pred},i}\right)^2}, \qquad \mathrm{SPPE} = \frac{S_{\mathrm{obs}}(T) - S_{\mathrm{pred}}(T)}{S_{\mathrm{obs}}(0)} \times 100.$$

Calibration used the warm-started procedure of §S9.3; the vectorised integrator used here reproduces the three-treatment optima of Table S29 to within 0.05 NLL units (`s14_integrator_check.py`).

**Table S35. Leave-one-treatment-out cross-validation.** Each model is calibrated on two treatments and used to predict the third with no refitting. Italicised rows give the Nash–Sutcliffe efficiency (NSE) for the in-sample best variant of each hazard structure (Table S29) and for the single-stage peak-based form, discussed below. The model that wins in sample (Table S29) is in bold. Predictive likelihoods are evaluated on the held-out treatment only, and are not comparable with the calibration likelihoods of Table S29.

| Model | A (2 d) | | | B (7 d) | | | C (15 d) | | |
|---|---|---|---|---|---|---|---|---|---|
| | Predictive NLL | NRMSE | SPPE | Predictive NLL | NRMSE | SPPE | Predictive NLL | NRMSE | SPPE |
| single stage, prop., cumulative | 146.6 | 0.176 | + 8.0 | 87.4 | 0.169 | − 1.5 | 121.6 | 0.151 | − 10.3 |
| | *NSE 0.966* | | | *0.970* | | | *0.957* | | |
| single stage, prop., peak-based | 78.8 | 0.175 | − 7.1 | 121.4 | 0.430 | − 8.1 | 166.1 | 0.472 | − 6.9 |
| | *NSE 0.967* | | | *0.807* | | | *0.576* | | |
| two stage, prop., cumulative | 137.7 | 0.170 | + 8.0 | 87.1 | 0.175 | − 1.2 | 122.0 | 0.151 | − 10.4 |
| **two stage, prop., peak-based** | 86.7 | 0.187 | + 7.4 | 79.8 | 0.062 | − 1.4 | 125.9 | 0.137 | − 4.2 |
| | *NSE 0.962* | | | *0.996* | | | *0.964* | | |
| two stage, MM, cumulative | 156.2 | 0.170 | + 8.0 | 87.1 | 0.176 | − 1.2 | 122.1 | 0.151 | − 10.4 |
| two stage, MM, peak-based | 86.7 | 0.188 | + 7.4 | 79.8 | 0.064 | − 1.3 | 124.5 | 0.143 | − 3.6 |

**Table S36. Aggregated across the three folds of Table S35.** Lower predictive NLL is better. “Worst fold” is the largest single-fold predictive NLL, so a model can lead on the total and still fail one treatment. Bold marks the model that wins in sample (Table S29); its saturating-repair form leads out of sample by 1.3 units.

| Model | Total predictive NLL | Worst fold |
|---|---|---|
| two stage, MM, peak-based | 291.0 | 124.5 |
| **two stage, proportional, peak-based** | **292.3** | **125.9** |
| two stage, proportional, cumulative | 346.8 | 137.7 |
| single stage, proportional, cumulative | 355.6 | 146.6 |
| two stage, MM, cumulative | 365.4 | 156.2 |
| single stage, proportional, peak-based | 366.2 | 166.1 |

**What the cross-validation shows.** The peak-based hazard is the worst of the six models out of sample on a single-stage damage trajectory and gives the best two once the damage model carries an upstream stage — a swing of 73.9 predictive likelihood units produced entirely by the damage model, with the hazard structure unchanged.

Adding saturating repair to the winning model improves the out-of-sample total by 1.3 units while costing 2.2 AIC units in sample (Table S29) — a difference too small to change any conclusion, and in opposite directions. The single-stage peak-based model predicts 60% survival on day 10 of the held-out 15-day treatment against 24% observed (NRMSE 0.472) — the fitted counterpart of the 68% that §4.2 obtains without fitting — which is the failure of §4.2 reproduced prospectively rather than retrospectively.

**Nash–Sutcliffe efficiency, reported for comparability with the evaluation practice of this literature, does not discriminate here.** The worst fold gives 0.957 for the in-sample best cumulative variant and 0.962 for the in-sample best peak-based one (italicised rows above); only the single-stage peak-based form, the worst model on every other measure, falls to 0.807 and 0.576 on the seven- and fifteen-day folds. The reason is in the statistic: survival in these treatments falls from complete to zero, so the variance in the denominator is large and any model tracking the broad decline scores above 0.95, whereas the predictive likelihood and NRMSE respond to the shape of the curve between pulses: the total predictive likelihood separates the out-of-sample best variant of each structure by 54 units (Table S36), and NRMSE on the fifteen-day fold separates them by a factor of 1.1.

**Two qualifications.** In fold A the single-stage peak-based model has the lowest predictive NLL of the six, while no better on NRMSE. The upstream half-life is less well determined by two treatments than by three, 1.3 to 16 d across the folds against 1.53 d (1.13–2.41 d) on the full series, because only the 15-day treatment leaves a long clean-water interval.

#### S15. Robustness: Digitisation Error, Identifiability, and the Window Rule

Three analyses follow. The first two target the decisive comparison of §4.3 — the 11.99-unit AIC margin of the two-stage peak-based model over the single-stage cumulative one in Table S29 — through the error in reading the survival points (§S15.1) and the identifiability of the parameters that carry its mechanistic reading (§S15.2). The third targets the window census of §S3 (§S15.3).

##### *S15.1 Error in reading survival off the published figure: a bootstrap*

The comparison that carries §4.3 is between the single-stage cumulative model at three parameters and the two-stage proportional peak-based model at four, with $\Delta\mathrm{AIC} = +11.99$ on the data as read. Survival points taken from Figure 2 of the source carry an estimated error of $\pm 0.03$ in surviving fraction (§S2). Each replicate perturbs the read points by $\varepsilon \sim \mathrm{Unif}(-0.03,+0.03)$, restores monotonicity, re-rounds to counts and refits both models from the unperturbed optima. Because the perturbation moves each read point by at most two animals, the third version below needs no random draws: treatment B's day-22 point cannot move, the count before it being already zero, and treatment A's can become only 4, 5 or 6 animals of 70, with probabilities 5/21, 5/21 and 11/21. Its column is that exact distribution.

Three versions were run (Table S37), in increasing order of how accurately they describe where the error actually lies. The first perturbs every point except $t = 0$, which assumes the whole series was digitised. The second holds fixed the three values of treatment C stated in the source text. The third perturbs only the two points that are genuinely read from the plotted curve alone — treatment A on day 22 and treatment B on day 22 — which Table S2 shows to be the correct description.

**Table S37. Digitisation bootstrap.** $\Delta$AIC > 0 favours the peak-based structure; $|\Delta\mathrm{AIC}| < 2$ is the tie band used throughout.

| | Whole series (120 rep.) | Text points fixed (100 rep.) | Two read points (exact) |
|---|---|---|---|
| unperturbed $\Delta$AIC | + 11.99 | + 11.99 | + 11.99 |
| median | + 4.17 | + 4.31 | **+ 11.99** |
| interquartile range | − 0.93 to + 8.06 | + 1.27 to + 7.49 | + 8.28 to + 11.99 |
| 2.5th–97.5th percentile | − 7.18 to + 17.66 | − 4.02 to + 16.38 | **+ 4.32** to + 11.99 |
| $P(\Delta\mathrm{AIC}>0)$ | 0.717 | 0.820 | **1.000** |
| $P(\Delta\mathrm{AIC}>2)$ | 0.617 | 0.700 | **1.000** |
| $P(\Delta\mathrm{AIC}<-2)$ | 0.200 | 0.090 | **0.000** |

**Under the correct description the comparison is robust**: perturbing the two read points leaves the peak-based structure ahead by at least 4.3 AIC units whatever the error, by more than 8 with probability 0.76 and by the full 11.99 with probability 0.52, and the sign never turns. The first two columns are retained because they bound the answer from the pessimistic side: had more of the series been read from the plot, the median would fall to 4.2–4.3 and the comparison would reverse beyond the tie band in one replicate in five or one in eleven, because perturbation degrades the shape-dependent peak-based fit more than the endpoint-dependent cumulative one.

##### *S15.2 Which parameters the data identify: profile likelihoods*

Three quantities carry the mechanistic reading of §4.3 — the upstream clearance rate $k_{\mathrm{off}}$, the half-saturation constant $K$ and the terminal repair rate $k_r$ — and were profiled by fixing each on a grid and re-optimising the remainder from several starts. $k_{\mathrm{off}}$ and $k_r$ are profiled on the four-parameter proportional-repair model that §4.3 quotes, $K$ on the five-parameter Michaelis–Menten model, the only one that has it, with starts including each of two independently obtained optima (Table S29).

**Table S38. Profile likelihood in the upstream clearance rate, two-stage proportional peak-based model.** Each row fixes $k_{\mathrm{off}}$ and re-optimises the remaining three parameters; the bold row is the optimum, and $\Delta$ is the excess over it. Selected rows of a 37-point grid (`s15_profile_koff.py`, §S16). Rate constants per hour, as in the source implementation.

| $k_{\mathrm{off}}$ ($h^{-1}$) | upstream half-life (d) | profile NLL | $\Delta$ |
|---|---|---|---|
| 0.0060 | 4.81 | 283.73 | 5.75 |
| 0.0101 | 2.87 | 281.03 | 3.06 |
| 0.0120 | 2.41 | 279.90 | 1.93 |
| 0.0151 | 1.92 | 278.58 | 0.60 |
| **0.0189** | **1.53** | **277.97** | **0** |
| 0.0226 | 1.28 | 278.56 | 0.59 |
| 0.0255 | 1.13 | 279.89 | 1.92 |
| 0.0301 | 0.96 | 283.14 | 5.16 |
| 0.0450 | 0.64 | 300.56 | 22.59 |

The 95% profile-likelihood interval, where the profile crosses 1.92 units above the minimum (located by interpolation between grid points), is $k_{\mathrm{off}} \in [0.0120, 0.0255]$ $h^{-1}$, that is, an upstream half-life between 1.13 and 2.41 days. The parameter is well determined, and the likelihood rises more steeply towards shorter half-lives than towards longer ones: a half-life of 0.64 d costs 22.6 units, one of 4.8 d costs 5.8.

**Table S39. Profile likelihood in the half-saturation constant, peak-based structure.** Each row fixes $K$ and re-optimises the rest.

| $K$ | profile NLL |
|---|---|
| $3 \times 10^{-8}$ | 278.108 |
| $8 \times 10^{-7}$ | 278.108 |
| $2 \times 10^{-5}$ | 278.108 |
| $10^{-4}$ | 278.108 |
| $2.5 \times 10^{-3}$ | 278.108 |
| $1.3 \times 10^{-2}$ | 278.108 |
| $6.3 \times 10^{-2}$ | 278.104 |

**The half-saturation constant is not identified.** The profile is flat across seven orders of magnitude, because the fitted maximum repair rate is small enough ($8 \times 10^{-5}$ h$^{-1}$, a removal half-life of order $10^2$ d) that the shape of a negligible term cannot be recovered from the data. Any statement locating $K$ relative to peak damage would be a statement about one point on a flat ridge, and none is made here. The same conclusion is reached from the other direction in Table S29: the Michaelis–Menten model does not improve on the proportional one, so the parameter that distinguishes them is doing no work.

**Terminal repair rate.** The repair rate of the four-parameter model was profiled in the same way (Table S40). The likelihood is flat from $k_r = 10^{-6}$ to $2 \times 10^{-4}$ h$^{-1}$ — repair half-lives from $10^2$ to $10^4$ days, all indistinguishable — then rises by up to 8 units for half-lives between 5 and 46 days, then falls again to within 0.4 units of the optimum at a half-life near 1.8 days.

**The repair rate is not identified**, and the admissible set is not even connected: the data are consistent with no terminal repair at all and with fast repair, but not with the intermediate rates. This is the third independent line reaching the same conclusion as Table S29 and Table S39, that the repair axis is doing no work in this dataset.

**Table S40. Profile likelihood in the terminal repair rate.** Each row fixes $k_r$ and re-optimises the rest; the bold row is the optimum.

| $k_r$ (h$^{-1}$) | repair half-life (d) | profile NLL |
|---|---|---|
| $10^{-6}$ | 28881 | 278.15 |
| $8.6 \times 10^{-6}$ | 3367 | 278.10 |
| $7.4 \times 10^{-5}$ | 393 | **277.97** |
| $2.2 \times 10^{-4}$ | 134 | 278.19 |
| $6.3 \times 10^{-4}$ | 46 | 279.39 |
| $1.9 \times 10^{-3}$ | 16 | 282.82 |
| $5.4 \times 10^{-3}$ | 5.3 | 285.92 |
| $1.6 \times 10^{-2}$ | 1.8 | 278.36 |

##### *S15.3 Whether the census depends on how the window is defined*

The window rule of §S3 contains two free choices: the fraction of peak exposure below which exposure is deemed to have ended, and the allowance for a multi-stage cascade to reach its maximum. Both were varied over the full census and the tests repeated with the same multiplicity control. The compound the choices bear on hardest is carbaryl, whose five treatments and their default windows are drawn in Figure S2.

**Table S41. Treatments firing at $q_{BH}$ < 0.05 out of treatments testable, for each combination.** Multiplicity control is over the same set as Table S6: every treatment of the archive that the combination leaves with a window of at least one day and that has concurrent controls, plus the two diazinon treatments, whose windows are set by the fitted damage maximum (§S6) and do not move with the allowance. The number of tests therefore varies from 18 to 30. Uncouplers are 2,4,5-trichlorophenol, 2,4-dichlorophenol and DNOC; narcotics 1,2,3-trichlorobenzene and 2,4-dichloroaniline; reactive compounds 4-nitrobenzyl chloride and Sea-Nine.

| Exposure end | Cascade allowance | Median window (d) | Tests | uncouplers | narcotics | aldicarb, carbofuran | carbaryl | malathion | diazinon | reactive |
|---|---|---|---|---|---|---|---|---|---|---|
| 1% | 1 d | 5.0 | 23 | 0/5 | 1/5 | 0/3 | 3/5 | 1/2 | 2/2 | 1/1 |
| 1% | 2 d | 4.0 | 21 | 0/4 | 0/5 | 0/3 | 3/5 | 1/2 | 2/2 | 0/0 |
| 1% | 3 d | 3.0 | 18 | 0/4 | 0/4 | 0/2 | 0/4 | 1/2 | 1/2 | 0/0 |
| 5% | 1 d | 5.0 | 29 | 0/8 | 1/6 | 0/4 | 4/5 | 2/3 | 2/2 | 1/1 |
| **5%** | **2 d** | **4.0** | **28** | **0/8** | **0/6** | **0/4** | **3/5** | **2/3** | **2/2** | **0/0** |
| 5% | 3 d | 3.0 | 22 | 0/6 | 0/4 | 0/2 | 0/5 | 1/3 | 1/2 | 0/0 |
| 10% | 1 d | 5.0 | 30 | 0/8 | 1/6 | 0/4 | 4/5 | 2/3 | 2/2 | 2/2 |
| 10% | 2 d | 4.0 | 29 | 0/8 | 0/6 | 0/4 | 3/5 | 2/3 | 2/2 | 0/1 |
| 10% | 3 d | 3.0 | 23 | 0/6 | 0/4 | 0/2 | 0/5 | 1/3 | 1/2 | 0/1 |

**Varying the rule changes how many treatments fire, and at a one-day allowance which compounds.** In all nine combinations the three uncouplers, aldicarb and carbofuran fire in no treatment, and malathion treatment A and diazinon treatment A fire in every one. A one-day allowance, which opens each window a day earlier, adds 1,2,3-trichlorobenzene treatment B and the 4-nitrobenzyl chloride treatment, and at 10% the Sea-Nine treatment. Carbaryl fires in four of five at a one-day allowance (A–D), three at the default (B–D) and none at three days (Table S41). That is a power effect: a longer allowance shortens the two-day windows of treatments A–C to a single day and removes deaths from them, and with those three small $p$-values gone the remaining treatment D ($p$ = 0.012) no longer clears the Benjamini–Hochberg threshold. The same loss of accompanying small $p$-values takes diazinon treatment B ($p$ = 0.010) below the threshold at a three-day allowance, although the diazinon windows, sixteen and eleven days, are untouched by the allowance; treatment A fires in every combination. No combination makes an uncoupler or a short-window carbamate fire.

### Part V. Availability and References

#### S16. Code and Data Availability

Every number and figure in this paper is reproduced by the scripts below, deposited with the data at Zenodo as Technical Report C0 (https://doi.org/10.5281/zenodo.23488395); on acceptance this Supporting Information joins the deposit as version 1.0 under the same concept DOI. Each analysis script begins with the number of the section it belongs to, and its first line states the tables or figures it reproduces; `run_all.py` runs them in order, and `MANIFEST.csv` and the README of the archive list what each reads and writes, the shared `lib_` modules, and where to obtain the published source data files, which are not redistributed. Table, figure and section numbers with an S refer to this document; section numbers without one refer to the main text.

Technical report with complete proofs: ref 35 (version 1.0, https://doi.org/10.5281/zenodo.23356185), deposited with `s10_prop2_violation_search.py`, `s10_sandwich_certificate.py`, `s01_prop2_short_separations.py` and `T0_sigma_ordering_check.py` and their outputs.

**Table S42. Script inventory, in the order of this document.** `MANIFEST.csv` in the archive gives the files each script reads and writes.

| Script | Reproduces |
|---|---|
| ***Part I. Theory*** | |
| `s01_prop2_short_separations.py` | Proposition 2 at short separations, ten repair laws (§S10) |
| `s01_v3_logconcavity.py` | Hypothesis (V3) for the fitted inputs (§S1) |
| ***Part II. Data and designs*** | |
| `s03_window_census.py` | The window census and the clean-water tests of the 2015 archive, every spreadsheet layout (Tables S3 and S6) |
| `s03_window_census_2007a.py` | The same for the 2007a archive (Tables S3, S6 and S8) |
| `s03_design_requirements.py` | §S3.1, Table S4 |
| `s03_class_window.py` | §S3.2, Table S5 |
| ***Part III. What the test measures*** | |
| `s04_pulse_intensity.py` | §S4, Table S7 |
| `s04_cleanwater_q.py` | The *q*-values of Table S6, over all twenty-eight tests |
| `s04_decline_rates.py` | §S4, Tables S8 and S9 |
| `s04_rate_intervals.py` | Profile-likelihood intervals on the decline rate (Tables S8 and S18) |
| `s04_compound_test.py` | §S4.2, Table S11 |
| `s04_frailty_regression.py` | §S4.3, Table S12 and the compound-level checks after it |
| `s04_half_lives.py` | Internal half-lives of Table 1 and the DNOC half-life of §S4 and §S12 |
| `s05_external_studies.py` | §S5, Table S13 |
| `s06_diazinon_one_experiment.py` | §S6, Tables S14 and S15, Figure S3 |
| `s06_diazinon_reduced_model.py` | Windows and tests under the reduced model (§S6) |
| `s07_immobilisation_and_counts.py` | §S7, Tables S16 and S17 |
| `s07_trend_tests.py` | §S7, the trend and Fisher tests of Table S16 |
| `s07_second_pulse_window.py` | §S7, Table S18 |
| `s07_census_rule_window.py` | The census rule applied to the interval series, at two allowances (§4.7, §S7) |
| `s08_recovery_ratios.py` | §S8, Table S19 and Figure S5 |
| ***Part IV. Model fits and controls*** | |
| `s09_stage_selection.py` | §S9.5, Table S22 |
| `s10_metric_scan.py` | §S10, Tables S24 and S25 |
| `s10_slow_cascade.py` | §S10, the slow-cascade check |
| `s10_prop2_violation_search.py` | §S10.1, Table S26 and the values drawn in Figure S8 |
| `s10_sandwich_certificate.py` | §S10.1, the sandwich certificate and the softening series; §S10.2, Table S27 |
| `s10_prop3_factor_two.py` | §S10.2, Table S28 |
| `s11_dmagna_fits.py` | §S11, the Michaelis–Menten rows of Table S29; the model shared by §S11–§S15 |
| `s11_dmagna_warm_starts.py` | §S11, the nested warm starts and the parameters of Figure 2 and Table S30 |
| `s11_dmagna_single_stage.py` | §S11, the single-stage rows of Table S29 |
| `s11_dmagna_two_stage_proportional.py` | §S11, the two-stage proportional rows of Tables S29 and S31, and the *D. magna* row of Table S34 |
| `s11_dmagna_treatment_C_only.py` | §S11, the "Treatment C only" columns of Table S29 |

| Script | Reproduces |
|---|---|
| `s11_symmetric_extension.py` | The cumulative hazard on the upstream state (§S11, Table S29, last row) |
| `s11_frozen_time.py` | Time the running maximum spends frozen on the fitted trajectory (§S11) |
| `s11_peak_times.py` | Damage peak times behind Figure 2 |
| `s11_parameter_free_check.py` | The parameter-free comparison of §4.2 and §S11.2 (Table S33), and the exoskeleton depot test |
| `s12_two_species_fits.py` | §S12, the *A. aquaticus* and *C. horaria* rows of Table S34 |
| `s12_dnoc_fits.py` | §S12, the DNOC variants |
| `s13_eroding_tolerance.py` | §S13 |
| `s14_cross_validation.py` | §S14, Tables S35 and S36, including NSE, and the full refit behind Tables S29–S31 |
| `s14_integrator_check.py` | §S14, the vectorised integrator against the loop |
| `s15_bootstrap.py` | §S15.1, Table S37 |
| `s15_profile_koff.py` | §S15.2, Table S38, and the interval quoted in §4.3 |
| `s15_profile_K.py` | §S15.2, Table S39 |
| `s15_profile_kr.py` | §S15.2, Table S40 |
| `s15_window_sensitivity.py` | §S15.3, Table S41 |
| ***Figures*** | |
| `fig_data.py` | The data drawn in the figures (`C-figure_data.json`) |
| `fig_1_2_3_S1_S4.py` | Figures 1, 2 and 3, and Figures S1 and S4 |
| `fig_4_dnoc.py` | Figure 4 (arguments `A` and `B` draw the other two DNOC treatments) |
| `fig_S2_carbaryl.py` | Figure S2 |
| `fig_S5_S7.py` | Figures S5 and S7 |
| `fig_S6_stages.py` | Figure S6 |
| `fig_S8_violation.py` | Figure S8 |
| `fig_toc.py` | TOC graphic |
| ***Shared modules*** | |
| `lib_paths.py` | Where every script finds its inputs and writes its outputs |
| `lib_fast_integrator.py`, `lib_guts_core_v2.py` | Numerical integrators |
| `lib_neonic_cascade.py` | The two-stage cascade of §S8 and §S13 |
| `tktd_core.py`, `case_diazinon.py` | The diazinon model and data reader of the companion paper (ref 4), for §S6 |

#### S17. References Cited Only in the Supporting Information

Reference numbers throughout this document follow the main text (references 1–30). The five references below are cited only here and continue that numbering.
